# Tandem repeat variation shapes immune cell type-specific gene expression

**DOI:** 10.1101/2024.11.02.621562

**Authors:** Hope A. Tanudisastro, Anna S.E. Cuomo, Ben Weisburd, Matthew Welland, Eleanor Spenceley, Michael Franklin, Angli Xue, Hao Lawrence Huang, Blake Bowen, Jianan Fan, Oscar A. Dong, Albert Henry, Peter C. Allen, Kristof Wing, Owen Tang, Michael Gray, Andre L.M. Reis, Jonathan Margoliash, Nehir E. Kurtas, Jeffrey M. Pullin, Arthur S. Lee, Harrison Brand, Michael Harper, Katalina Bobowik, Michael Silk, Amy Miniter, John Marshall, Vivian Bakiris, Bindu Swapna Madala, Caitlin Uren, Caitlin Bartie, Rachael A. McCloy, Anne Senabouth, Harriet Dashnow, Liam Fearnley, Alejandro Martin-Trujillo, Egor Dolzhenko, Zhen Qiao, Stuart M. Grieve, Tung Nguyen, Eyal Ben-David, Ling Chen, Kyle Kai-How Farh, Michael Talkowski, Stephen I. Alexander, Owen M. Siggs, Leonhard Gruenschloss, Hannah R. Nicholas, Jennifer Piscionere, Cas Simons, Chris Wallace, Melissa Gymrek, Ira W. Deveson, Alex W. Hewitt, Gemma A. Figtree, Katrina M. de Lange, Joseph E. Powell, Daniel G. MacArthur

**Affiliations:** Centre for Population Genomics, Garvan Institute of Medical Research, and UNSW Sydney, Sydney, NSW, Australia; Centre for Population Genomics, Murdoch Children’s Research Institute, Melbourne, VIC, Australia; Faculty of Medicine and Health, University of New South Wales, Sydney, NSW, Australia; Faculty of Medicine and Health, University of Sydney, Sydney, NSW, Australia; Garvan-Weizmann Centre for Cellular Genomics, Garvan Institute of Medical Research, Sydney, NSW, Australia; UNSW Cellular Genomics Futures Institute, University of New South Wales, Sydney, NSW, Australia; Program in Medical and Population Genetics, Broad Institute of MIT and Harvard, Cambridge, MA, USA; Analytic and Translational Genetics Unit, Massachusetts General Hospital, Boston, MA, USA; Faculty of Science, University of Sydney, Sydney, NSW, Australia; Menzies Institute for Medical Research, University of Tasmania, Hobart, TAS, Australia; Department of Ophthalmology, Royal Hobart Hospital, Hobart, TAS, Australia; Centre for Eye Research Australia, University of Melbourne, East Melbourne, VIC, Australia; Charles Perkins Centre, The University of Sydney, Sydney, NSW, Australia; Kolling Institute of Medical Research, Royal North Shore Hospital, Sydney, NSW, Australia; Genomics and Inherited Disease Program, Garvan Institute of Medical Research, Sydney, NSW, Australia; Department of Medicine, University of California San Diego, CA, USA; Massachusetts General Hospital, Boston, MA, USA; The Broad Institute of MIT and Harvard, Cambridge, MA, USA; MRC Biostatistics Unit, University of Cambridge, UK; Department of Biomedical Informatics, University of Colorado Anschutz Medical Campus, Aurora, CO, USA; Population Health and Immunity Division, The Walter and Eliza Hall Institute of Medical Research, Parkville, VIC 3052, Australia; Department of Medical Biology, University of Melbourne, Parkville, VIC, Australia; Bruce Lefroy Centre, Murdoch Children’s Research Institute, Parkville, VIC, Australia; Department of Genetics and Genomics Sciences and Minidich Child Health and Development Institute, Icahn School of Medicine at Mount Sinai, New York, NY, USA; Pacific Biosciences of California, Menlo Park, CA, USA; Imaging and Phenotyping Laboratory, Charles Perkins Centre, Faculty of Medicine and Health, The University of Sydney, Australia; Lumus Imaging, St George Private Hospital, Kogarah NSW 2217; Illumina Artificial Intelligence Laboratory, Illumina, San Diego, CA, USA; Centre for Kidney Research, Kids Research Institute and the Department of Nephrology, The Children’s Hospital at Westmead, Westmead, NSW, Australia; Cambridge Institute of Therapeutic Immunology & Infectious Disease, University of Cambridge, UK; Department of Computer Science and Engineering, University of California San Diego

## Abstract

Tandem repeats (TRs) - highly polymorphic, repetitive sequences across the human genome - are important regulators of gene expression but remain underexplored due to challenges in accurate genotyping and analysis^1^. Here, we generate new whole genome and single-cell RNA sequencing from >5.4 million blood-derived cells across 1,925 individuals in two cohorts [Cuomo *et al.*, accompanying manuscript], and perform meta-analysis to characterize the impact of variation in >1.7 million TR loci on immune cell type-specific gene expression. We identify >69,000 single-cell expression TR loci (sc-eTRs), 30.7% of which are specific to one of 28 immune cell types, and reveal dynamic regulatory effects using cell-state inference. Matched single-cell ATAC sequencing profiles from >3.4 million nuclei in 922 individuals [Xue *et al.*, accompanying manuscript]. uncover chromatin accessibility QTLs for nearly one-third of expression-associated TRs, supporting coordinated effects on cis-regulatory architecture. Fine-mapping implicates 1,490 TRs as candidate causal drivers of gene expression in 6.1% of tested genes, and colocalization analyses highlight >200 genes in which TRs likely mediate genetic associations with immune and hematological traits. Together, these results provide a genome-wide, multiomic view of TR-mediated regulation in the human immune system, establishing TRs as key contributors to cell type-specific regulatory variation and complex trait architecture.

## Introduction

Tandem repeats (TRs) - highly polymorphic repetitive sequences characterized by 1-6 bp motifs (short tandem repeats [STRs]) and >6bp motifs (variable number tandem repeats [VNTRs]) - represent one of the most mutable classes of human genetic variation with profound biological effects^1^. They occupy a larger proportion of the genome than protein-coding genes^2,3^, can mutate rapidly^4^, and influence gene expression by regulating transcription factor binding^2^, altering nucleosome occupancy^5^, and modulating the activity of promoters^6^ and enhancers^7^. Despite their functional potential, TRs have been understudied in population genetics because short-read sequencing and alignment historically performed poorly in repetitive regions, limiting accurate genotyping^1^. However, methodological improvements have begun to address these challenges, enabling more precise and comprehensive characterization of TRs^1^.

Although rare repeat expansions are well-established causes^8^ of Mendelian disorders, such as Huntington’s disease and Fragile X Syndrome, emerging evidence indicates that common TR variation may contribute more broadly to complex traits. Common TR variation has been associated with gene expression^9^, epigenetic modification^10^, serum biomarker levels^11^, cancer susceptibility^12^, and complex disease risk^13^. However, inconsistent locus definitions^3^ and reliance on imputation from surrounding SNVs^11,14^ have hampered systematic comparison across studies and obscured the true contribution of TRs to regulatory variation.

While most analyses of TR-mediated expression effects have relied on bulk RNA-seq^3,9^ (which averages signals across heterogeneous cell populations), single-cell RNA sequencing (scRNA-seq)^15^ now enables the systematic mapping of expression quantitative trait loci (eQTLs) at a finer resolution at population scale. Single-cell analysis provides insight into cellular heterogeneity^16^, differentiation pathways^16^, and disease progression^17^. Given their high mutability and regulatory potential, TRs represent a compelling but largely unexplored class of variants that may tune state-dependent transcriptional programs in ways not captured by SNV-based approaches.

Here we present a genome-wide catalog of single-cell expression quantitative trait TR loci (sc-eTRs) from peripheral blood in Phase 1 of TenK10K, the largest paired human whole genome and single-cell RNA-seq resource to date [Cuomo *et al.*, accompanying manuscript (https://doi.org/10.1101/2025.03.20.25324352)]. We quantify their cell type specificity, reveal dynamic effects through cell state inference, assess evidence for causality, uncover coordinated chromatin accessibility changes in matched single-cell ATAC-seq chromatin accessibility QTL (caQTL) analyses, and evaluate colocalization with immune and hematological traits, providing an integrated view of TR-mediated regulation in the human immune system.

## Results

### Genome-wide polymorphic TR discovery

To quantify how TR length variation influences gene expression across 28 immune cell types, we first identified polymorphic TR loci in 1,925 individuals from Phase 1 of TenK10K, each with matched high-coverage whole genome sequencing (WGS) and scRNA-seq data (**Fig. 1a**). The 1,925 individuals analyzed in this study consists of individuals from the Tasmanian Ophthalmic Biobank cohort (950 European ancestry individuals with no signs of ocular disease) and the BioHEART cohort (975 European ancestry individuals with diagnosed or suspected cardiac disorders) (**Methods**) (**Supplementary Fig. 1**).

**Figure 1:**
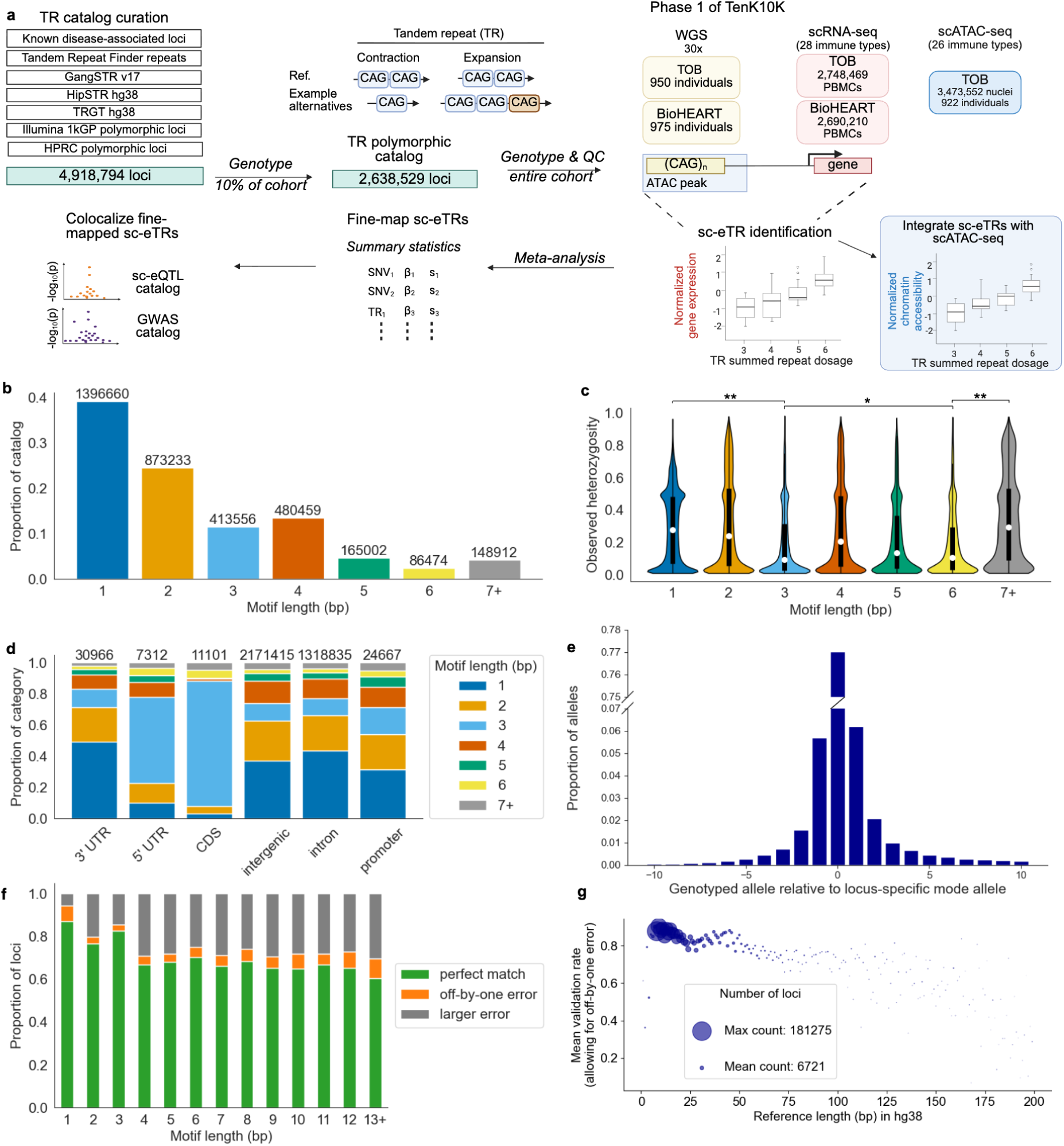
Genome-wide polymorphic TR curation and validation. **a,** Schematic overview of the study workflow: tandem repeat (TR) catalog curation, single-cell expression quantitative trait TR loci (sc-eTR) identification, integration with scATAC-seq, fine-mapping, and colocalization analyses. TOB, Tasmanian Ophthalmic Biobank; PBMCs, peripheral blood mononuclear cells. **b**, Distribution of motif lengths for polymorphic TR loci included in the curated catalog. **c**, Observed heterozygosity of polymorphic TR loci stratified by motif length. Violin plots depict the distribution of locus-specific heterozygosity, highlighting median values (white circles), interquartile ranges (IQR; boxes indicate 25th to 75th percentiles), and whiskers extending to ±1.5×IQR. Motif-length comparisons were tested using two-sided Mann-Whitney U tests. \*\**P* < 1 x 10^-30^; \**P* = 6.5 x 10^-32^ **d**, MANE gene annotation distribution for polymorphic TR loci, categorized by motif length. CDS, coding sequence. **e**, Frequency distribution of genotyped alleles relative to the locus-specific modal allele, considering alleles within ±10 repeats of the mode. **f**, Validation of ExpansionHunter genotypes compared against TRGT genotypes obtained from PacBio HiFi long-read sequencing (n = 25 samples). Concordance categories include perfect matches (exact dosage agreement), off-by-one errors (±1 repeat difference), and larger discrepancies (labeled ‘larger error’), stratified by motif length. **g**, Mean genotype validation rates for ExpansionHunter calls (allowing off-by-one errors) benchmarked against TRGT long-read genotypes, plotted by TR locus reference length (in base pairs, bp). The horizontal axis is truncated at 200 bp, capturing >99.9% of the analyzed loci.

Most TR genotyping tools require loci to be specified *a priori*, meaning that catalog curation is crucial for sensitivity. To generate a genome-wide TR catalog, we merged several reference-based^18–21^ and population-derived polymorphic TR catalogs, using samples from the 1000 Genomes Project and the Human Pangenome Reference Consortium^22^, generating a unified catalog of 4,918,794 loci^23^, with some loci containing multiple TRs (**Methods**). To select for variation informative for association testing, we genotyped ∼10% of individuals and retained 2,638,529 polymorphic loci. Homopolymer and dinucleotide motifs comprised the majority of the polymorphic loci (**Fig. 1b**), whereas tri- and hexanucleotides exhibited the lowest heterozygosity, consistent with these motifs having lower mutation rates^24^ (**Fig. 1c)**. Most polymorphic TRs were intergenic or intronic, while coding sequences were strongly enriched for frame-preserving trinucleotide motifs (**Fig. 1d**).

We next genotyped all individuals at polymorphic loci using ExpansionHunter. Across loci, allele lengths were tightly distributed around the modal allele, with 97% of calls falling within 10 repeat units, consistent with stepwise mutation dynamics^25^ (**Fig. 1e, Extended Data Fig. 1**). After applying stringent locus- and genotype-level quality filters, the final callset comprised ∼1.7 million high-confidence autosomal TRs with a median of six alleles per locus (**Methods**).

To assess accuracy we benchmarked ExpansionHunter TR genotypes against two truth sets. Concordance with capillary electrophoresis genotypes was 95.0% (**Supplementary Fig. 2**, **Supplementary Note 1**, **Methods**) and comparison with PacBio HiFi long-read sequencing in 25 individuals showed 81% exact agreement, rising to 86.5% with ±1 repeat tolerance (**Fig. 1f-g**, **Supplementary Note 1**).

### Immune cell landscape of sc-eTRs

We mapped *cis* single-cell expression quantitative trait TR loci (sc-eTRs) by testing associations between 1,732,938 common autosomal TRs and the expression of 22,213 genes across 28 immune cell types. Using associaTR, we performed pseudobulk eQTL mapping within 100 kb of each gene while controlling for technical and biological covariates, and combined results across the two cohorts via fixed-effects meta-analysis to mitigate cohort-specific sequencing effects (**Fig. 1a**, **Supplementary Fig. 4-9**, **Methods**). Associations involving tandem repeats are referred to as sc-eTRs, and those involving single nucleotide variants/indels as sc-eSNVs.

We identified 69,210 unique sc-eTRs associating with 15,889 eGenes at a gene-level false discovery rate (FDR) of 5% (**Fig. 2a**, **Extended Data Fig. 2**, **Supplementary Table 1**). Type 1 error was well-controlled by permutation (**Fig. 2a, Methods**). The number of sc-eTRs correlated with the number of cells sequenced per cell type (Pearson *r* = 0.90, *P* = 7.94×10^-11^), but we detected sc-eTRs in every cell type (**Fig. 2a**). Meta-analysis substantially increased discovery power, revealing sc-eTRs for 349 eGenes that did not reach significance in either cohort alone (**Supplementary Fig. 10a**). Sensitivity analyses confirmed an approximately linear relationship between sample size and eGene discovery (**Supplementary Fig. 10b**).

**Figure 2:**
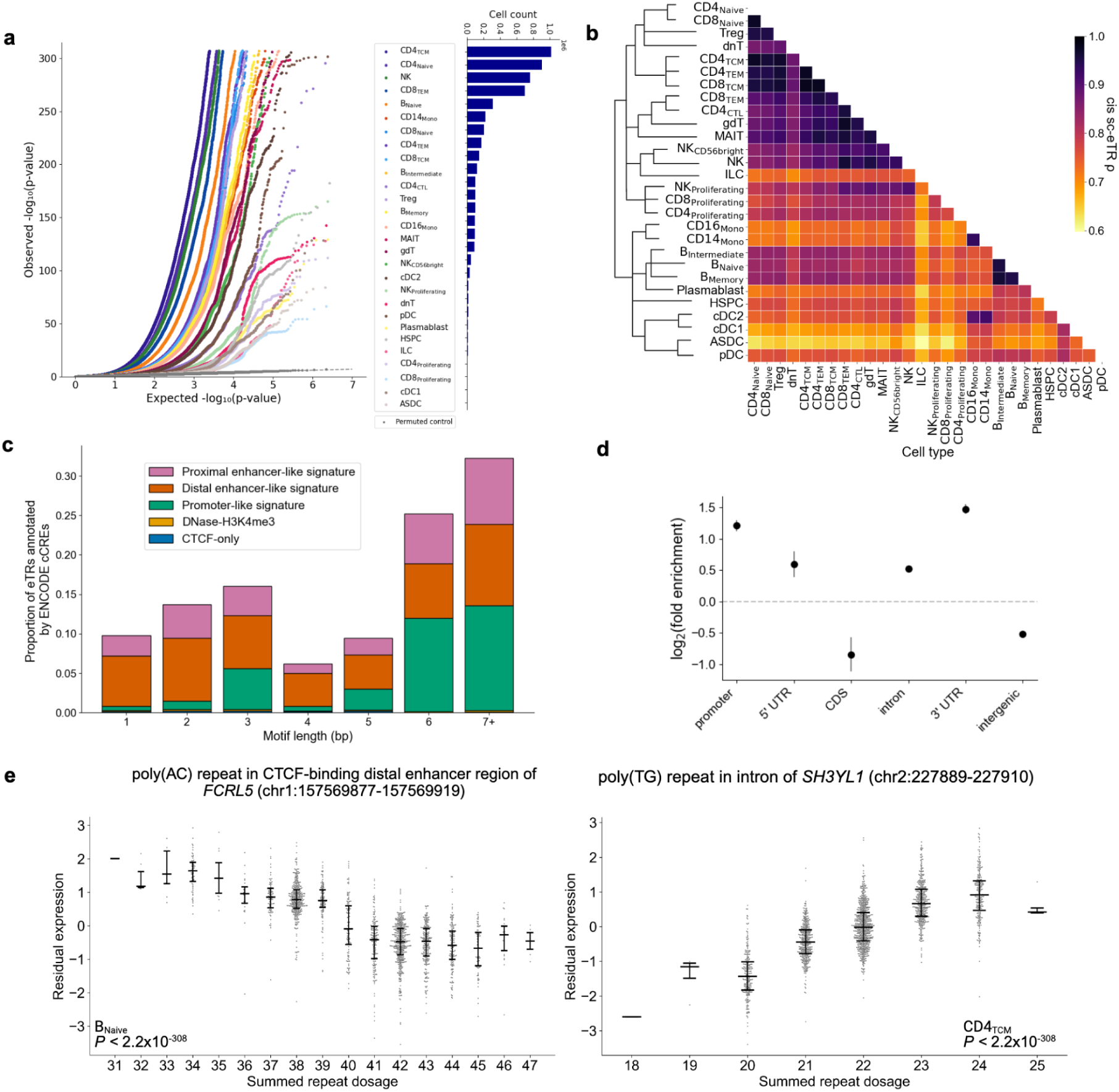
single-cell eTR discovery. **a,** Quantile-quantile (QQ) plot comparing observed two-sided *P* values for each TR x gene test (colored by cell type) against the expected uniform distribution (dashed gray line). Permuted control is highlighted in dark gray (n=1,925). Bar plot on the right shows the number of cells sequenced per cell type in the study cohort (n=1,925 individuals). **b**, Heatmap of pairwise correlations of *cis* sc-eTR posterior effect sizes across cell types, adjusted for global effect size sharing and power differences with multivariate adaptive shrinkage^26^. Row ordering reflects scRNA-seq cell type classification by scPred^57^ (**Methods**). **c**, ENCODE candidate *cis* regulatory element (cCRE) annotation of sc-eTRs, by motif length. **d**, Log_2_(fold enrichment) of sc-eTRs relative to all TRs genotyped (n = 2,638,529 loci), by MANE gene annotation. Points indicate observed log₂(fold enrichment) and error bars indicate 95% confidence intervals. **e**, Examples of eTRs previously identified with whole blood bulk RNA-seq^9,34^, now resolved to specific immune cell types using TenK10K Phase 1 data (n = 1,925 individuals). Horizontal bars indicate the first quartile, median, and third quartile. P values were calculated using two-sided Wald tests of the regression coefficient (β) and are uncorrected for multiple testing; for both comparisons, *P* < 2.2 x 10^-308^.

To assess cell type similarity, we used multivariate adaptive shrinkage^26^ (mashR) to estimate posterior effect sizes across cell types. Pairwise correlations recapitulated known immunological hierarchies: B-cell subsets showed strong convergence, whereas lymphoid and myeloid lineages were more distinct (**Fig. 2b, Supplementary Note 2**). type-2 conventional dendritic cells (cDC2) and monocytes showed strong concordance, consistent with shared marker expression^27^. The pairwise correlation of posterior effect sizes of sc-eTRs was comparable to that of sc-eSNVs (**Supplementary Fig. 11**).

sc-eTRs were enriched in ENCODE candidate *cis* regulatory elements (cCREs) (odds ratio [OR] = 1.96, Fisher’s exact test, *P* < 2.2×10^-16^), particularly in distal enhancer-like signatures (**Fig. 2c**). They were also enriched in proximal regulatory regions and depleted in intergenic regions (**Fig. 2d**), and were more likely to overlap cCREs than sc-eSNVs (**Supplementary Fig. 12**). Motif-enrichment analyses revealed marked dependence on genomic context: longer motifs (≥6 bp) were enriched in CDS and 5′ UTRs, whereas intergenic and intronic regions showed depletion of 2+ bp motifs (**Supplementary Fig. 13**).

Across all sc-eTRs, increasing repeat length generally corresponded to higher gene expression (median effect size =0.018, Wilcoxon signed-rank test, *P* = 1.78×10^-7^). This trend was strongest in 5′ UTRs, introns, and 3′ UTRs (**Extended Data Fig. 3**). Effect sizes varied by genomic context, with TRs in promoter-like signatures showing larger median effects than in distal enhancers (0.05 vs 0.02, Mann Whitney U, *P* = 3.1×10^-3^), whereas intergenic sc-eTRs exhibited broader effect size distributions compared to those in proximal regulatory elements (Levene’s test, *P* = 1.0×10^-5^). The greatest variability in effect sizes was seen at lower starting dosages (<20 repeats), possibly reflecting higher polymorphism or sampling variance (**Supplementary Fig. 14**).

To evaluate how TR regulatory effects vary across continuous cellular states, we applied scDeepID, a biologically informed deep learning framework (**Methods**). Focusing on B-cell subsets, we inferred three functional axes - protein localization to membrane^28^, activation of NK cells^29^, and response to multicellular organisms^30^ - and stratified cells by low and high activity states (≤15th and >85th percentile of cell function scores, respectively). This revealed 579 cell state eTRs (FDR < 5%), with establishment of protein localization to the membrane producing the greatest number **(Supplementary Table 2**). For example, the effect of a poly(TTTG) repeat on *SYNGR1* expression is modulated by natural killer cell activation activity, with the effect of increasing repeats increasing *SYNGR1* expression being strongest in intermediate B cells with lower levels of NK cell activation activity (*P =* 6.80 x 10^-8^, **Extended Data Fig. 4**). *SYNGR1* encodes a synaptic vesicle protein and has previously been associated with rheumatoid arthritis^31^; its knockdown has been demonstrated to impact cytokine production by NK cells^32^. Finally, cell state eTRs were enriched in enhancer-like signatures, consistent with previous findings of cell state interacting eQTLs^33^ (**Supplementary Fig. 15**).

While mashR provides insights into global sharing patterns among broadly testable sc-eTRs, it does not capture loci that are tested (or genes that are expressed) in only a subset of immune cell types. To address this, we adapted the classification framework described in Cuomo *et al*. [Cuomo *et al.*, accompanying manuscript]] by performing ∼3 million pairwise comparisons, taking into account differential gene expression, significance of association, LD, and effect directionality (**Supplementary Table 1**, **Methods**). On a gene level, 13.3% of eGenes exhibiting specific regulation (max scenario = 1 or 2), whereas the remainder demonstrated broader sharing (max scenario ≥ 3) (**Methods, Extended Data Fig. 5**). The majority of sc-eTRs (69.3%) had concordant effects in at least two cell types, and these tended to lie closer to transcription start sites (Mann Whitney U, *P* = 1.57×10^-32^). In contrast, cell type-specific sc-eTRs (scenarios 1-3 and 5) were enriched in intergenic regions and depleted in proximal regulatory elements (**Supplementary Fig. 16**).

Motif composition also differed by sharing pattern: cell type-specific sc-eTRs were enriched for di- and tetranucleotide motifs (OR = 1.23, 1.10; *P* = 1.15 x 10^-32^, 2.27 x 10^-5^, respectively), repeat classes frequently intersecting transcription factor binding sites^2^, whereas shared sc-eTRs were enriched for homopolymers (OR = 1.13, *P* = 1.82 x 10^-20^). Mean GC content was modestly higher in cell type-specific sc-eTRs compared to shared sc-eTRs (0.130 vs 0.123; Mann-Whitney U, *P* = 7.51×10^-8^), consistent with preferences of distal regulatory or architectural elements like CTCF binding sites.

Finally, comparing our results with bulk whole blood eTRs from GTEx^9,34^ (n = 652 individuals), 91.5% overlapped nominally significant sc-eTRs (*P* < 0.05) in at least one cell type (**Supplementary Table 3**, **Methods**). Single-cell resolution revealed cell type-specific drivers for loci previously observed in bulk. For example, the poly(AC) enhancer repeat near *FCLR5* was resolved to B naive cells, consistent with the role of the gene in B cell receptor signaling^35^. Additionally, the poly(TG) intronic repeat in *SH3YL1* was mapped to CD4_TCM_ cells, reflecting the role of *SH3YL1* in T cell modulation^36^ (**Fig. 2f**). Signals unique to sc-eTRs were enriched for cell type specificity (scenarios 1-3 and 5) compared to those overlapping bulk eTRs identified in GTEx (OR = 3.09, Fisher’s exact test, *P* = 2.3×10^-127^).

### TR-linked epigenetic-expression effects

To investigate how TRs regulate gene expression, we leveraged single-cell ATAC-seq profiles from 922 individuals from Phase 1 of TenK10K [Xue *et al.*, accompanying manuscript (https://doi.org/10.1101/2025.08.27.25334533)]. For each sc-eTR, we performed chromatin accessibility QTL (caQTL) mapping by testing whether TR length variation was associated with chromatin accessibility of nearby ATAC peaks (±10 kb of the TR) in the corresponding cell type.

Across cell types, 19,231 TRs were significantly associated (FDR < 5%) with at least one nearby ATAC peak, indicating that nearly one-third (31.8%) of expression-associated TRs associate with alterations of local chromatin structure (**Supplementary Fig. 17, Supplementary Table 4**). These caQTL+ sc-eTRs tended to exhibit stronger associations with gene expression (median |β_expression_| = 0.17 vs. 0.13 for caQTL- sc-eTRs; Mann Whitney U, *P* < 2.2 x 10^⁻16^) (**Fig. 3a**), likely reflecting both true biological coupling and, in part, improved ascertainment of well-profiled TRs.

**Figure 3:**
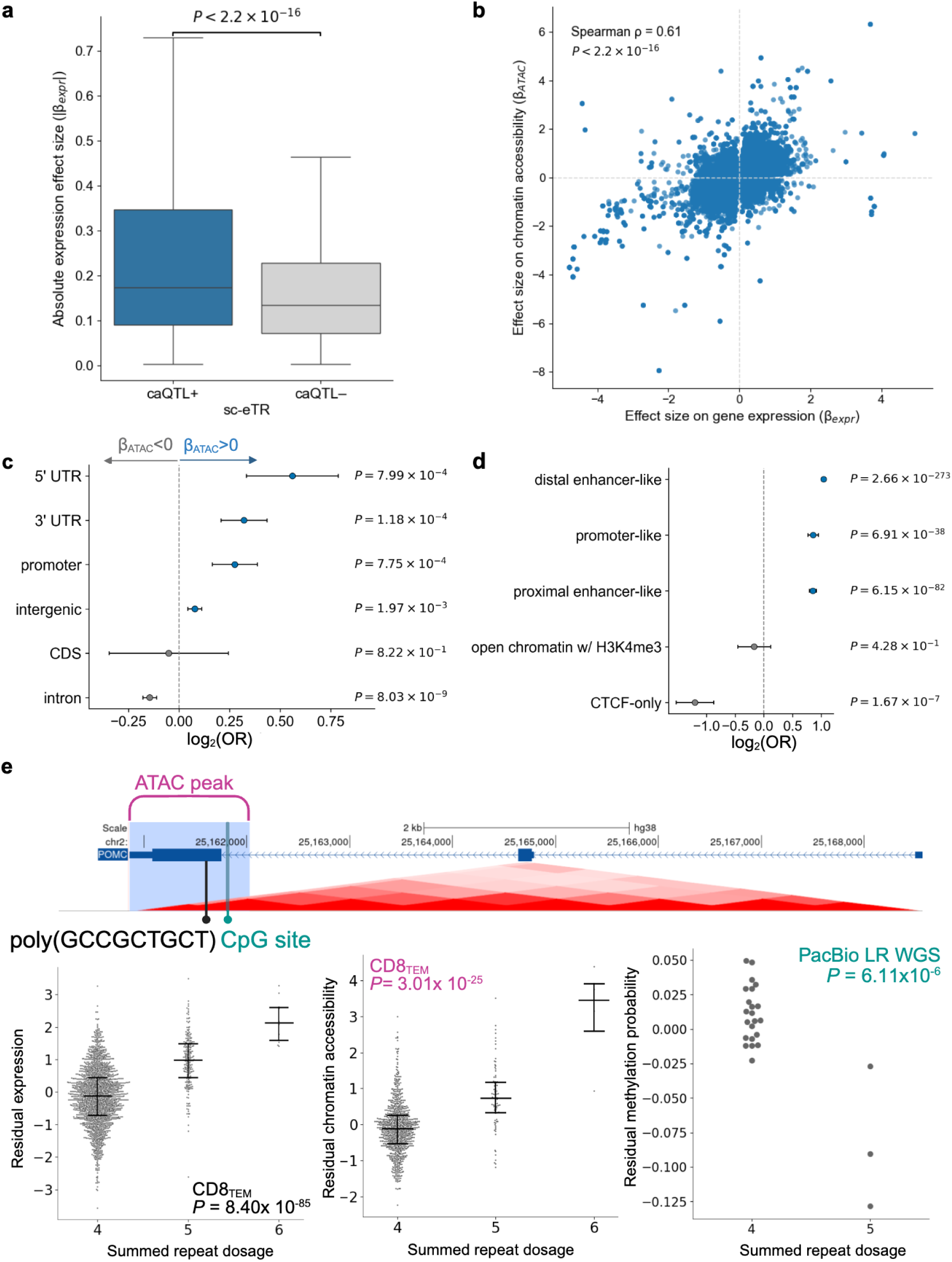
Tandem repeats coordinate *cis*-regulatory effects on chromatin accessibility and transcription at single-cell resolution. **a**, Boxplot showing the distribution of absolute expression effect sizes for TRs associated with chromatin accessibility (caQTL+ sc-eTRs) versus those without (caQTL- sc-eTRs) (n = 112,100 sc-eTRs). Centre lines show medians; box limits show interquartile range (IQR); whiskers extend to 1.5 × IQR; outliers omitted for clarity. caQTL+ sc-eTRs show larger effect sizes (one-sided Wilcoxon *P* < 2.2x 10^⁻16^). **b**, Scatterplot of TR effect sizes on gene expression versus chromatin accessibility across caQTL+ sc-eTRs, showing positive correlation (Spearman ρ = 0.61, two-sided *P* < 2.2 x 10^-16^). **c**, Observed log_2_(odds ratio) (center point) of caQTL+ sc-eTRs with positive (β_ATAC_ > 0) versus negative effect sizes (β_ATAC_ < 0) across MANE genomic annotations (n=19,231 distinct TRs). Two-sided Fisher’s exact P values (uncorrected); error bars denote 95% confidence intervals. CDS, coding sequence. **d**, Observed log_2_(odds ratio) (center point) of caQTL+ sc-eTRs across ENCODE cCRE annotations relative to caQTL- sc-eTRs (n=112,100 sc-eTRs). Two-sided Fisher’s exact P values (uncorrected); error bars denote 95% confidence intervals. CTCF-only denotes CTCF binding sites without overlap with other cCRE annotations. **e**, Locus zoom plot and association swarmplots for a coding poly(GCCGCTGCT) repeat in *POMC* in CD8_TEM_ using scRNA-seq (n = 1,925 individuals), scATAC-seq (n = 922 individuals), and PacBio long-read sequencing methylation data (n = 25 individuals). Horizontal bars indicate the first quartile, median, and third quartile. P values were calculated using two-sided Wald tests of the regression coefficient (β) and are uncorrected for multiple testing. Longer alleles at this TR locus are associated with elevated gene expression, increased chromatin accessibility, and decreased methylation. Locus in **e** and **Extended Data** Fig. 6 were selected to illustrate diverse multiomic regulatory architectures.

Among caQTL+ sc-eTRs, 50.9% showed increased chromatin accessibility with increasing repeat length (β_ATAC_ > 0) (**Fig. 3b**). TRs associated with decreased accessibility were modestly enriched in introns (OR = 1.10, Fisher’s exact, *P* = 1.1 × 10⁻⁸), while those with positive effect sizes were enriched in 5′ UTRs (OR = 1.48, *P* = 6.6 × 10⁻⁴), promoters (OR = 1.21, *P* = 7.0 × 10⁻⁴), and 3′ UTRs (OR = 1.25, *P* = 1.2 × 10⁻⁴) (**Fig. 3c**). These findings suggest that longer TR alleles may preferentially promote chromatin accessibility, and thereby enhance gene expression, when located within regulatory elements.

Amongst caQTL+ sc-eTRs, TR effects on expression and accessibility were positively correlated (Spearman ρ = 0.61, *P* < 2.2× 10^⁻16^), with 78% of caQTL+ sc-eTRs exhibiting concordant effect directions on expression and chromatin accessibility (**Fig. 3b**).

For example, a poly(GCCGCTGCT) repeat within the coding sequence of *POMC* not only associates with expression in CD8_TEM_ cells (*n* = 1,925, *P* = 8.40 × 10^-85^), but also with chromatin accessibility at an overlapping ATAC peak (*n* = 922, *P* = 3.01 × 10^-25^) (**Fig. 3e**). Using DNA methylation data from 25 individuals with PacBio HiFi sequencing in Phase 1 of TenK10K (**Methods**), we further found that repeat length was associated with methylation levels at a nearby CpG site (*n* = 25, *P* = 6.11 × 10^-6^). This tandem repeat has previously been implicated as a causal driver of *POMC* expression across multiple bulk tissues^34^. We extend these findings by demonstrating that the repeat’s regulatory influence persists within CD8 effector memory T cells, consistent with T cells producing POMC, a precursor protein for immune cell signalling^37^.

Similarly, an intronic poly(CTCCTCCCAGTGACCC) repeat in *INF2* associates with gene expression (*n* = 1,925, *P* = 3.72 × 10^-184^; **Extended Data Fig. 6**) and chromatin accessibility (*n* = 922, *P* = 3.08 × 10^-61^) in CD4_TCM_ cells and DNA methylation levels at a nearby CpG site (*n* = 25, *P* = 7.2 × 10^-6^), consistent with INF2 promoting microtubule formation required for centrosome reorganization in T cells^38^. Together, these examples illustrate the layered regulatory architecture of TRs revealed by our multiomic approach.

Genome-wide analyses further revealed that caQTL+ sc-eTRs were strongly enriched in promoter-like, proximal enhancer-like, and distal enhancer-like ENCODE cCREs, and depleted from CTCF-only sites, consistent with their role as constitutive insulators (**Fig. 3d**). Even stronger enrichments were observed for caQTL+ sc-eTRs intersecting CTCF-binding sites, suggesting that some TRs may alter CTCF occupancy and influence long-range enhancer-promoter contacts (**Supplementary Fig. 18**). caQTL+ sc-eTRs were also enriched in topologically associating domain (TAD) boundaries (OR = 1.23, *P* = 2.79 x 10^-8^), consistent with clinically relevant TRs colocalizing in these chromatin domain boundaries^39^.

Motif analysis using HOMER^40^ showed striking enrichments of motifs of caQTL+ sc-eTRs amongst zinc finger transcription factors, including KLF, LRF/ZBTB7A, NRF, and OCT families, pointing to coordinated TR effects within regulatory networks governing lymphocyte identity and activation^41^ (**Supplementary Table 5**).

Finally, integrating chromatin accessibility with our previously defined framework of regulatory sharing revealed that caQTL+ sc-eTRs were strongly enriched among cell type-restricted regulatory events [scenario 1] (OR = 2.53, *P* = 9.48 × 10^-18^) and modestly enriched for shared, concordant effects across multiple cell types [scenario 4] (OR = 1.09, *P* = 4.53 × 10^-16^) (**Methods, Extended Data Fig. 5**). These results indicate that TR-associated chromatin remodeling contributes both to cell type-specific transcriptional programs and to regulatory processes conserved across immune lineages.

### Features of candidate causal sc-eTRs

Despite higher genotyping uncertainty than SNVs, 11,099 genes showed an sc-eTR as the lead (or tied lead) association in their *cis* window. Lead sc-eTRs in the best-powered cell type exhibited weaker LD with the next most significant neighbouring SNV (proxy SNV) than did lead SNVs (**Supplementary Fig. 19**). Generally, LD strength was inversely correlated with the number of distinct TR alleles at the locus (**Extended Data Fig. 7**), suggesting that higher mutability can erode correlation with surrounding SNV haplotypes and that a subset of TR-driven regulatory effects may be poorly captured by nearby SNVs. Lead sc-eTRs also showed large differences in association strength relative to their proxy SNVs (**Supplementary Fig. 20**), suggesting either large underlying effect sizes or particularly robust TR genotyping, marking them as strong candidates for causality.

Fine-mapping across 15,889 eGenes identified 1,490 TRs as candidate causal sc-eTRs (PIP ≥ 0.7), spanning 1,354 eGenes (**Supplementary Table 6**, **Methods**). Among the gene-cell type combinations with fine-mapped signals, 40 supported multiple causal TRs and 577 supported both a TR and a SNV. 4.2% harbored independent SNV and TR signals in distinct credible sets. However, the vast majority of loci contained fine-mapped signals from only one variant type (81.7% with SNVs alone, 14.1% with TRs alone).

Candidate causal sc-eTRs were markedly enriched in proximal regulatory elements, including promoter-like cCREs (OR = 11.59), 5′ UTRs (OR = 9.72), and promoters (OR = 7.55), and were depleted in distal intergenic regions (OR = 0.46, all with Fisher’s exact test *P* < 2.2×10^-16^, **Extended Data Fig. 7).** Enrichments were stronger for candidate causal sc-eTRs than for all sc-eTRs (**Supplementary Fig. 21**), underscoring that TRs most likely to drive expression changes preferentially localize to proximal regulatory regions.

Sequence composition further distinguished candidate causal sc-eTRs. GC-rich repeats (≥80% GC) were enriched (OR = 3.43, *P* < 2.2×10^-16^), consistent with their localization to promoters and CpG islands^42^. Evolutionary analyses revealed that candidate causal sc-eTRs preferentially overlapped transcription factor binding sites (TFBS) within constrained primate-specific elements^43^, Zoonomia^44^ Group 2 regions (characterized by more rapid evolutionary turnover), and deeply conserved mammalian elements (**Supplementary Fig. 22**). These enrichments were strongly GC-dependent: TRs with 61-80% GC showed the highest TFBS enrichment within constrained primate elements (OR = 15.43, *P* = 0.001), whereas those with ≥81% GC were enriched for DNase hypersensitivity summits (OR = 3.77, *P* = 1.5×10^-8^), suggesting that GC-rich candidate causal sc-eTRs may preferentially modulate regulatory architectures shaped by recent selection.

eGenes associated with candidate causal sc-eTRs were enriched in the least constrained decile of gene-level loss-of-function observed/expected upper bound fraction (LOEUF) scores (binomial test *P* = 3.13×10^-87^, **Supplementary Fig. 23-24**), consistent with prior reports that dosage-sensitive genes under stronger purifying selection are relatively depleted for expression-modifying variation^45^.

Direct TR genotyping revealed novel fine-mapped associations: candidate causal sc-eTRs showed only moderate LD with lead SNVs (median r² = 0.57, IQR = 0.48, **Supplementary Fig. 25**), and more than half of them remained significant (FDR < 5%) after conditioning on the lead SNV (**Supplementary Table 7**). These results suggest that a substantial subset of candidate causal sc-eTRs capture regulatory variation not explained by nearby SNVs.

As illustrative cases, a CTCF-binding poly(GTGC) repeat in the 5′ UTR of *RUNDC3B* was fine-mapped across nine lymphoid cell types (minimum *P* = 2.71×10^-89^, **Fig. 4c**), consistent with lymphoid-specific promoter methylation at this locus^46^. A poly(AC) repeat 52 kb upstream of *KIR3DL2* was similarly finemapped in NK cells (*P* = 1.46×10^-116^; **Fig. 4d**), consistent with the gene’s role as an HLA class I receptor^47^ in NK cells. We also fine-mapped a GC-rich 5′ UTR repeat in *PLA2G4C*, spanning B-cell, monocyte, and dendritic lineages (minimum *P* = 2.91×10^-255^; **Fig. 4e**). PLA2G4C encodes a phospholipase generating lipid signalling precursors^48^, suggesting that TR variation at this locus may influence immune-relevant lipid pathways.

**Figure 4:**
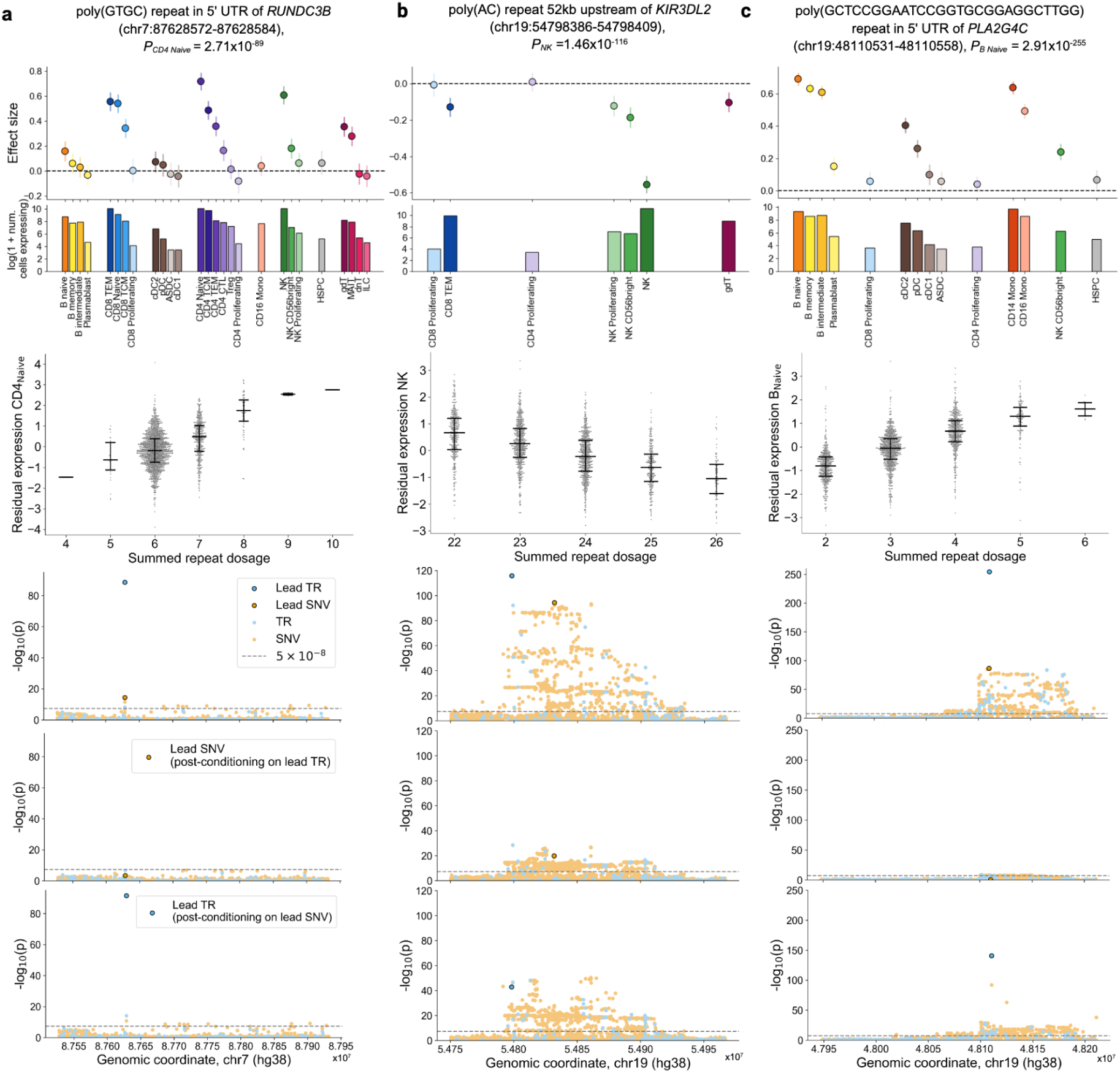
Case examples of candidate causal sc-eTRs. **a-c**, For each locus, the top panel provides the distribution of observed sc-eTR effect sizes (center point) across immune cell types exhibiting sufficient gene expression for association testing, with error bars representing 95% confidence intervals. The accompanying bar plot shows the number of cells expressing the respective gene per cell type. Subsequent plots detail the association between repeat length and residual gene expression (after adjusting for covariates; n = 1,925 individuals) [horizontal bars indicate the first quartile, median, and third quartile], nominal association signals for the candidate causal sc-eTR, lead eSNV, and other variants within ±100 kb of the gene body, and conditional association analyses. Specifically, associations are shown conditioned first on the candidate causal sc-eTR genotype, and then separately conditioned on the genotype of the lead eSNV. Detailed regional LD plots corresponding to these examples are available in **Extended Data** Fig. 6. P values were calculated using two-sided Wald tests of the regression coefficient (β) and are uncorrected for multiple testing.

Finally, we explored disease relevance of candidate causal sc-eTRs. While rare repeat expansions in >60 TR loci cause monogenic disorders^8^, the role of common variation at these loci is less understood. Unexpectedly, we found nominal associations at 52 disease-associated loci, and fine-mapped seven - *C9orf72*, *CACNA1A, CBL, CSTB*, *DIP2B*, *GLS*, and *PPP2R2B* - as candidate causal regulators of expression in blood-derived cells (**Supplementary Table 8**, **Supplementary Note 3**, **Supplementary Fig. 26**). Moreover, 24.9% of eGenes linked to candidate causal sc-eTRs overlapped with genes in PanelApp, a database of clinically relevant gene panels (**Supplementary Table 9**), suggesting potential roles in variable penetrance, modifier effects, or second-hit mechanisms.

Collectively, these findings position TRs as a distinct and underappreciated layer of regulatory variation that shapes human gene expression, with potentially important implications for both complex traits and rare disease.

### sc-eTRs likely drive complex traits

We next asked whether candidate causal sc-eTRs could help explain variation in complex traits. We first intersected candidate causal sc-eTRs with significant TR PheWAS hits from UK Biobank (UKBB)^13^ and found that those associated with hematological traits were enriched in 5’ UTRs (OR = 4.59 Fisher’s exact test, *P* = 0.0003, **Fig. 5a**, **Supplementary Table 10**, **Supplementary Fig. 27**, **Methods**).

**Figure 5:**
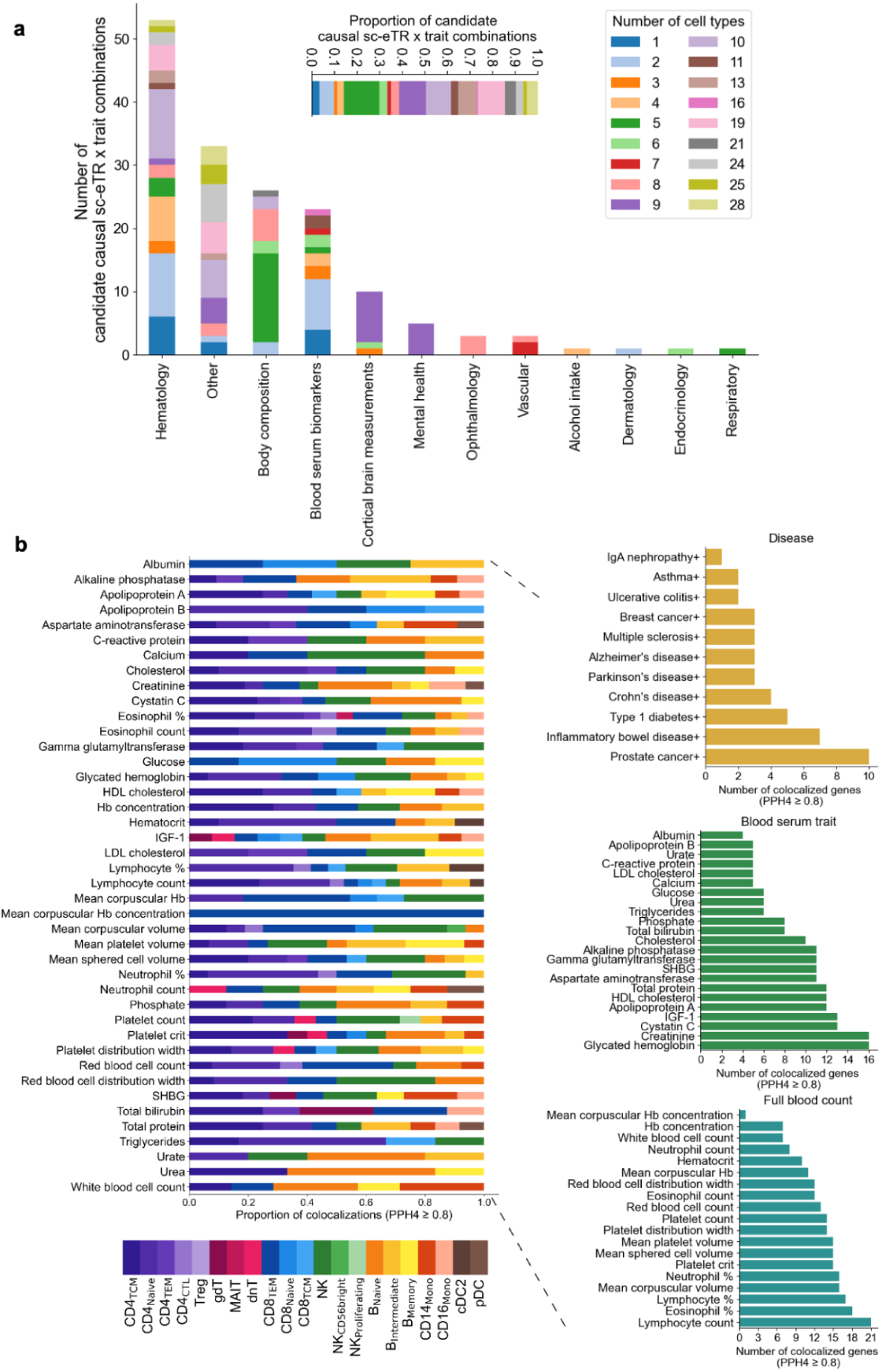
Candidate causal sc-eTRs in cell type-specific gene regulation associate with complex trait variation. **a**, Bar plot showing the number of candidate causal sc-eTRs that intersect TR UKBB PheWAS hits, colored by the number of cell types sharing each sc-eTR. Inset: bar plot depicting the proportion of candidate causal sc-eTRs overlapping UKBB PheWAS traits, colored by the number of cell types sharing the sc-eTR. **b,** Barplot of the number of colocalized genes (PPH4 ≥0.8) associated with a candidate causal sc-eTR, per GWAS catalog, divided into disease, blood serum trait, and full blood count categories (right panel). Only phenotypes with at least one colocalization were plotted. Phenotypes marked with + were colocalized using only SNVs, while unmarked phenotypes were colocalized using SNVs and imputed TRs. The cell type specificity of serum traits and full blood count phenotypes are illustrated in the left panel.

To test whether sc-eTRs help drive GWAS signals, we performed colocalization to compare our sc-eQTL data (sc-eSNV+sc-eTR) with two types of GWAS resources: (1) SNV-only GWAS for asthma, autoimmune, cancer, infectious, and neuroinflammatory conditions; and (2) joint SNV and imputed STR GWAS for 44 UKBB blood and serum traits^11^ (**Fig. 5b**, **Methods**). We used coloc^49^ to compute the posterior probability that the trait and gene expression were driven by the same variant, and considered an sc-eTR a plausible driver of the trait when (i) the gene-trait pair showed strong colocalization (PPH4 ≥ 0.8), (ii) the sc-eTR was confidently fine-mapped (PIP ≥ 0.7), and (iii) the sc-eTR was in at least moderate LD (r² ≥ 0.5) with a variant in the GWAS window.

Across traits, we identified 215 genes (502 gene-trait pairs) satisfying these criteria (**Supplementary Table 11**). The greatest number of colocalizations was observed for full blood count traits, followed by blood serum traits, and finally disease phenotypes (**Fig. 5b**). The proportion of loci showing strong colocalization varied by trait (e.g., 8.1% for lymphocyte percentage, 13.6% for Parkinson’s disease; **Supplementary Fig. 28**). Importantly, these results underestimate the true contribution of TRs to complex traits, as robust colocalization with SNV-only GWAS signals requires TRs to be well tagged by nearby SNVs.

Several loci illustrate how sc-eTRs can anchor trait associations to specific immune contexts using multiomic data. A poly(TG) repeat ∼3kb upstream of *LIME1* lies within a locus that colocalizes with asthma in CD4 naive cells (PP H4 = 0.87, *P* = 9.8 x 10^-164^, **Extended Data Fig. 9**) and was also associated with nearby chromatin accessibility variation (*P* = 3.62 x 10^-4^), consistent with *LIME1*’s role in T cell receptor signalling and its hypermethylation in atopy^50^. A poly(T) repeat in the *KIF16B* promoter lies in a locus that colocalizes with mean platelet volume across 13 immune cell types (maximum PP H4 = 1, minimum *P <* 2.2×10^-308^, **Extended Data Fig. 10**), supported by chromatin and methylation changes; KIF16B encodes a kinesin motor protein involved in trafficking secretory granules^51^, suggesting that TR variation may modulate immune crosstalk^52^.

Additionally, a poly(AAAC) intronic repeat in *ADH5* lies within a locus colocalizing with total bilirubin (PP H4 = 0.84, *P* = 3.75x 10^-53^, **Extended Data Fig. 11**) and associates with altered chromatin accessibility (*P* = 4.7x 10^-51^) in CD16 monocytes; ADH5 regulates STING-mediated innate responses in monocytes^53^ which can subsequently alter hepatic function^54^. Finally, a poly(CG) repeat in a distal enhancer of *UBA7* lies within a locus that colocalizes with inflammatory bowel disease in NK cells (PP H4 = 0.92, *P* = 8.58 x 10^-19^, **Extended Data Fig. 12**) and showed coordinated effects on chromatin accessibility (*P* = 2.25×10^-4^); UBA7 mediates interferon-stimulated gene conjugation, a pathway that can promote colonic inflammation^55^.

Together, these colocalizations suggest that tandem repeat variation can be a molecular driver of complex trait associations, with effects that emerge only in the appropriate immune cell context. By anchoring trait-linked variants to specific regulatory mechanisms in immune cells, we show that incorporating TRs into causal architecture can deepen our understanding of the pathways underlying blood, serum, and immune-mediated traits.

## Discussion

Our single-cell atlas of tandem repeat (TR) variation across 28 immune cell types reveals a regulatory landscape far more extensive than previously appreciated. With more than 69,000 sc-eTRs and 1,490 fine-mapped candidate causal TRs, our results show that repeat variation contributes substantially to gene regulatory architecture, often in ways that are poorly captured by surrounding SNVs. These findings underscore the need for direct genotyping of TRs, as imputation- and SNV-based approaches likely miss many potentially causal TR loci.

A central theme emerging from our study is that TRs act as context-sensitive regulators. Their effects are not only quantitative but can shift qualitatively with repeat length or cellular environment. For instance, at the *FRA10AC1* locus, we observed that repeat length positively associates with gene expression within the spectrum of common variation, while larger pathogenic expansions induce gene silencing^56^. Indeed, some sc-eTRs achieving statistical significance across multiple cell types may appear shared but can still exert distinct, cell type-specific effects on expression, as captured in our cell type specificity scoring framework. When considered alongside mashR and cell state inference results, these findings suggest that “specificity” is an emergent property shaped by both cellular context and the underlying distribution of alleles.

Our study further highlights the utility of multiomic data to elucidate how TR variation influences gene regulation. Nearly a third of sc-eTRs were associated with matched changes in local chromatin structure, providing evidence for TR-mediated remodeling of proximal regulatory elements. Notably, we found examples of sc-eTRs supported by ATAC-seq data modeling both cell type-specific regulatory landscapes as well as broadly shared transcriptional programs. Building a comprehensive TR-caQTL atlas will be essential for resolving how repeat variation shapes the regulatory genome in different immune contexts.

The substantial number of novel associations we discovered underscores the need for directly genotyped repeat-inclusive genomic catalogs. Such resources will be crucial for downstream inference - from Mendelian randomization to improved colocalization with complex trait GWAS - particularly in loci where TRs are poorly tagged by SNVs. Although TR genotyping has improved, short-read sequencing still struggles to accurately capture subsets of TR loci, such as those found in low complexity regions. Long-read genome sequencing will be key for establishing truth sets, calibrating genotyping models, and identifying contexts in which short-read sequencing is sufficient and those where the additional resolution of long-read sequencing is helpful.

More broadly, modeling TRs raises conceptual and analytic challenges that extend beyond those encountered for SNVs: multi-allelic dosage encodings, length-dependent instability, and the potential for length-dependent allele-specific regulatory effects. Addressing these issues will require tailored statistical frameworks, improved population structure correction, and functional validation using scalable CRISPR or synthetic biology approaches. As long-read genome and transcriptome datasets expand in well-phenotyped cohorts, the detection and dissection of allele-specific TR effects will become tractable at scale.

Overall, our results position TRs as a material component of human regulatory variation, influencing cell type-specific gene expression and contributing to the genetic basis of immune traits. Incorporating TRs into the broader landscape of genomic variation, together with SNVs, indels, and structural variants, will be essential for building a more comprehensive picture of human regulatory architecture and its role in health and disease.

## Supporting information

Supplementary Information Guide

## Methods

### TenK10K Phase 1 dataset processing

The TenK10K Phase 1 resource, including sample quality control, whole-genome sequencing, and single-cell RNA processing, is detailed in the accompanying manuscript by Cuomo *et al.* Phase 1 comprises 1,925 individuals with matched WGS and scRNA-seq from two cohorts: the Tasmanian Ophthalmic Biobank [TOB] (950 individuals of European ancestry recruited without ocular disease; cardiovascular phenotypes were not recorded) and BioHEART (975 individuals of European ancestry presenting with diagnosed or suspected cardiac conditions, including acute myocardial infarction, suspected coronary artery disease, and cardiothoracic or vascular surgery).

Peripheral blood was collected from all participants. TOB and BioHEART samples were prepared using KAPA Hyper PCR-free (Roche) and Illumina DNA PCR-free kits, respectively, and sequenced to 30× on the Illumina NovaSeq 6000 (2 × 150 bp). Reads were aligned to hg38 using DRAGMAP (v1.3.1) following DRAGEN-GATK best practices.

For scRNA-seq, PBMCs were isolated, captured using the single-cell 3’ gene expression high throughput (3’ GEX HT) kit v3.1 from 10X on a Chromium X machine, sequenced on the NovaSeq 6000, and aligned to hg38 (Gencode v44/Ensembl 110) using STAR^59^ within the 10X CellRanger software (v7.2.0). CellBender (v0.3.0) was used to identify residual ambient RNA in each sequencing library. Demultiplexing was performed with vireo^60^ (v2.1.0) via Demuxafy.

Individuals failing genotype or sample-level QC, exhibiting cryptic relatedness, or identified as ancestry outliers were removed, yielding a final set of 1,925 participants with paired WGS and scRNA-seq profiles.

This study complied with all relevant ethical regulations and was overseen by the Human Research Ethics Committees at the Royal Children’s Hospital (Melbourne, Australia), Northern Sydney Local Health District (Sydney, Australia), St. Vincent’s Hospital (Sydney, Australia), and University of Tasmania (Hobart, Australia). Informed consent was obtained from all participants.

### TR catalog curation

A catalog of 4,918,794 loci was generated by merging several existing catalogs: disease-associated loci (https://github.com/broadinstitute/str-analysis/blob/main/str_analysis/variant_catalogs); the hg38 reference catalogs from GangSTR v17 (1,340,266 loci)^19^, HipSTR (1,634,957 loci)^20^, and TRGT (171,146 loci)^61^; Illumina’s polymorphic catalog based on the 1000 Genomes Project (174,293 loci; https://github.com/Illumina/RepeatCatalogs); a polymorphic catalog derived from 51 haplotype-resolved assemblies from the Human Pangenome Reference Consortium (1,573,403 loci)^22,61^; and a catalog generated by Tandem Repeat Finder (TRF)^18^ (v4.09) consisting of hg38 TR loci ≥9 bp in length. The TRF catalog was generated using the parameters: match = 2, mismatch = 7, indel penalty = 1000000, PM = 80, PI = 10, minscore = 8, and maxperiod = 2000.

Merging of the TR catalogs is fully described in a separate manuscript by Weisburd and Dolzhenko *et al*.^23^, and the version of the catalog used in this study represents an earlier iteration of that framework, generated using tools and genome references available at the time of analysis. In brief, catalogs were parsed sequentially, and loci were added unless they overlapped an existing entry by ≥66% and shared the same repeat motif. To improve alignment performance and capture biologically coupled variation, locus boundaries were expanded to incorporate adjacent TRs when present, generating compound locus definitions that contain up to three TRs separated by ≤6 bp.

### Polymorphic TR catalog curation

To identify polymorphic TR loci, ExpansionHunter (v5)^62^ (threads = 16, analysis-mode = streaming) was run using the aforementioned genome-wide catalog on 200 randomly selected ‘XX’ individuals.

Using Hail Query v0.2.126 (https://github.com/hail-is/hail), we removed chrY, chrM, and monomorphic loci from the resulting VCF, producing a polymorphic catalog of 2,638,529 loci (**Data availability**).

### Long-read TR genotyping

Tandem repeats were genotyped in 25 PacBio HiFi sequencing samples using TRGT (v1.1.0) with a custom catalog of polymorphic tandem repeats. The resulting tandem repeat genotypes were then merged into a single VCF for downstream analysis. HiFi sequencing and alignment protocols are described in **Supplementary Methods**.

### TR genotyping and filtering

ExpansionHunter (v5)^62^ was used to genotype TRs from the polymorphic TR catalog, producing diploid repeat-length estimates per sample from realigned reads. VCFs were merged using mergeSTR (TRTools v5.0.2)^63^ and underwent locus-level quality control. We excluded loci with call rate < 90%, observed heterozygosity < 0.00995 (equivalent to a non-major allele frequency < 0.5%), or Hardy-Weinberg disequilibrium (binomial test, *P* < 1×10^-6^).

To mitigate genotype uncertainty, particularly at loci prone to expansion or contraction errors, we set genotypes to missing if either allele deviated by >30 repeat units below or >20 repeat units above the locus-specific modal allele. These thresholds were selected based on concordance with a publicly available truth set^22^, and reflect a conservative approach to exclude low-confidence calls, particularly at the tails of the allele-length distribution. All samples exhibited a per-sample TR call rate >99%.

### SNV genotyping and filtering

SNVs and indels were called using GATK4 HaplotypeCaller in DRAGEN mode following the same pipeline described in the accompanying manuscript by Cuomo *et al*. After joint genotyping, we removed monomorphic and rare variants (minor allele frequency < 1%). Multiallelic variants were split into biallelic variants. Sample- and variant-level QC followed the procedures detailed in Cuomo *et al.*. The resulting high-quality SNV set was used for downstream analyses alongside the TR genotype dataset.

### eTR and eSNV association analysis

To study the effect of TR length variation on gene expression across immune cell types, we mapped sc-eTRs using associaTR (TRTools v5.0.2)^63^, which performs length-based tests for multiallelic TRs against continuous phenotypes. An sc-eTR was defined as a TR-gene association detected in a given cell type. eQTL mapping was performed separately in the TOB and BioHEART cohorts, followed by meta-analysis.

scRNA-seq processing followed the framework described by Cuomo *et al.* Briefly, reads were aligned and quantified with cellranger (v7.2.0) (10X Genomics). Using scanpy^64^ (v1.8.2), we removed cells unassigned to individuals or flagged as doublets by ≥2 of scds^65^ (v.1.13.1), vireo^66^ or scDblFinder^67^ (v1.12.0), and excluded cells with >20% mitochondrial reads, <1,000 or >10,000 detected genes, or <800 reads detected. Cell types were assigned using a combined approach of scPred (v1.9.2)^57^, hierarchical progressive learning, scHPL^68^ (HierscPred v0.1.0), and trained on a published CITE-seq reference^69^. CellTypist^70,71^(v=1.6.2, model used ‘Immune_All_High’) was used to confirm cell typing of individual cells.

Single-cell counts were then processed into pseudobulk profiles. Using scanpy^64^ (v1.9.3), counts were normalized for sequencing depth, log-transformed, corrected for batch effects, and aggregated as means per individual and cell type (**Supplementary Fig. 9**), consistent with recommendations from Cuomo *et al.*^72^. For each cell type, genes expressed in <1% of cells were removed, and remaining expression values were quantile-normalized.

Expression values were adjusted for karyotypic sex, age, population structure, and technical effects. Population structure was captured by the top 12 SNV-derived PCs from QC-filtered European samples (**Supplementary Fig. 29-30**). Technical effects were accounted for using the top 6 PCs from PCA of pseudobulk expression profiles, selected after evaluating models with 1-10 PCs in CD14 monocytes, CD8_TEM_, and CD4_TCM_ (**Supplementary Fig. 31-33**).

For each TR within ±100 kb of a gene, we fitted an ordinary least squares model:

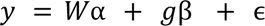

where *y* denotes normalized gene expression, *W* the covariate matrix, β the effect size the model is estimating, *g* the summed diploid TR repeat length, and ɛ a normally distributed error term. The same model was applied for SNVs and indels, with *g* representing the dosage of the non-reference allele.

A ±1 Mb scan on chromosome 1 in CD4_TCM_ cells showed that ∼81% of sc-eTRs lie within ±100 kb of the gene body (**Supplementary Fig. 34**), supporting the use of a ±100 kb cis window genome-wide.

### Meta-analysis

We performed fixed-effects meta-analysis of eQTL summary statistics from the TOB and BioHEART cohorts using the *meta* package in R (v4.0.0)^73^. Due to differences in library preparation protocols between cohorts, leading to distinct insert size and read depth profiles (**Supplementary Fig. 4-8**), we conducted association analyses separately and combined results by meta-analysis.

To assess potential inflation of Type 1 error, we performed a permutation of the sample identifiers in the CD4_TCM_ cell type and repeated the regression analysis and meta-analysis. Type 1 error was well controlled (**Fig. 2a**).

### Multiple testing correction

Gene-level p-values were computed by combining variant-level p-values of variants using the ACAT-V test^74^ based on the Cauchy combination. For each gene-cell type pair, we designated a set of top sc-eTRs as the variant(s) with the lowest nominal p-value. In each cell type, multiple testing correction was then performed across all gene-cell type tests using the Storey q-value procedure^75^, reporting results at FDR < 5%. Original (uncorrected) p-values are provided for illustrative examples in the main text, with the minimum p-value provided if the sc-eTR was observed in multiple cell types.

### Multivariate adaptive shrinkage (mash) analysis

To account for global effect size sharing and power differences across cell types, we ran mashR (v0.2.79)^26^, following the recommended workflow for eQTL analysis (https://stephenslab.github.io/mashr/articles/eQTL_outline.html). We assembled the effect sizes and standard errors for every sc-eQTL (FDR < 5%) that was tested in every cell type. These sc-eQTLs were used to compute the ‘data-driven’ covariance matrix using an extreme deconvolution method including the top five principal components. The ‘data-driven’ and ‘canonical’ covariance matrices were then used to fit the mash model to all chr2 variants tested in all cell types (‘random’ tests). The resulting mash model was used to compute posterior effect sizes for all sc-eQTLs across the 28 cell types. Analyses were performed separately for sc-eSNVs and sc-eTRs.

### Cell type specificity analysis

Following the framework used in the accompanying manuscript by Cuomo *et al.*, we quantified the cell type specificity of sc-eTRs by classifying each pairwise sc-eTR-gene relationship across 28 immune cell types into one of five scenarios. Scenario 1 captures cases where the gene was not expressed (<1% of cells) in the comparison cell type, while scenario 2 includes instances where the gene was expressed but no sc-eTR was detected. Scenario 3 encompasses independent signals, defined as distinct TRs associated with the same gene but in low linkage disequilibrium (LD; r² < 0.5). Scenario 4 represents shared or strongly linked sc-eTRs (r² ≥ 0.8) with concordant effects on gene expression; to conservatively account for haplotype tagging, we also included linked sc-eTRs with discordant effect directions when LD was high (r² ≥ 0.8). Scenario 5 captured cases involving the same sc-eTR or moderately linked sc-eTRs (0.5 ≤ r² < 0.8) with opposing effect directions. To adjust for differences in statistical power across cell types, we further refined scenario 2 assignments by performing pairwise meta-analyses: if the meta-analysis P-value was at least 10⁵-fold smaller than the original and the combined effect directions were concordant or opposing, we reclassified these pairs into scenario 4 or 5, respectively.

Unless otherwise specified, an sc-eTR was considered ‘shared’ if it had shared concordant effects (scenario 4) with at least one other cell type.

Because a given eGene can fall into different scenarios across different pairs of cell types, we summarized its overall specificity by taking the maximum scenario value (1-5) observed across any pairwise comparison for each eGene.

### caQTL association analysis

Single-cell chromatin accessibility data (scATAC-seq) from donors in the TOB cohort were processed as detailed in an accompanying manuscript by Xue *et al.* Briefly, PBMCs from 952 donors were pooled in 119 sets of eight, captured twice using the 10x Genomics scATAC-seq v2 platform, and sequenced on an Illumina NovaSeq 6000. Reads were aligned to GRCh38 with Cell Ranger ATAC (v2.2.0), and peaks were called using MACS2. Donor assignment and doublet detection were performed using vireo via Demuxafy2 (v2.1.0). After filtering for unassigned nuclei, doublets, and quality metrics (peak counts, fraction of reads in peaks, blacklist ratio, nucleosome signal, TSS enrichment), 3,472,552 high-quality nuclei from 922 donors were retained (mean ∼3,766 nuclei per donor).

Peak counts were aggregated per donor to create pseudobulk matrices for each repeat. Counts were normalized for GC-content using EDASeq (v.2.40.0), transformed to counts per million (CPM), and z-score standardized. Repeated pools were merged by averaging overlapping z-scores and re-standardizing each peak across donors. Principal components of chromatin accessibility were computed from these matrices. We conducted a series of sensitivity analyses to optimize the parameters used for association mapping, including peak calling methods, normalization methods, window size, and number of PCs, detailed in an accompanying manuscript by Xue *et al*.

For each sc-eTR within 10kb of a scATAC peak, we tested for TR-chromatin associations using associaTR (TRTools v5.0.2). Ordinary least squares regression was performed per cell type, with normalized accessibility as the outcome and summed TR repeat length as the predictor, adjusting for age, sex, 12 genotype PCs, and chromatin accessibility PCs.

A total of 52,873 TRs were tested in 26 cell types (excluding CD8 Proliferating and ILCs due to low donor numbers) across 165,149 ATAC peaks. Each TR was tested only in the cell types where it was identified as an sc-eTR. Multiple TRs near the same peak were tested independently. P-values were adjusted using the Benjamini-Hochberg procedure to control the false discovery rate at 5% within each cell type.

### Cell state inference and cell state-dependent eTR mapping

Cell function scores were computed using scDeepID, a multi-task transformer guided by biological pathway databases, further detailed in Cuomo *et al.*. We trained scDeepID on 539,820 canonical B cells (naive, intermediate, memory) from TenK10K Phase 1 scRNA-seq data using 2,351 highly variable genes (selected via Scanpy, min_mean=0.0123, max_mean=3, min_disp=0.5) and human Biological Process Gene Ontology (GOBP) pathways. We then extracted the attention-based latent space for the UMAP representation, and extracted the cell function scores followed by normalization of each cell function score to zero mean and unit variance.

Three cell functions with clear activity gradients across B cell subtypes were selected: protein localization to membrane^28^, activation of NK cells^29^, and response to multicellular organisms^30^. Cells were stratified into low (≤15th percentile), medium (15-85th percentile), and high (>85th percentile) activity bins within each B cell subtype. For ‘protein localization to membrane’, which showed a continuous gradient across B intermediate and B memory subtypes, we aggregated these subtypes prior to activity binning.

Pseudobulk profiles for cell type x activity bin combinations were generated, and sc-eTR mapping was performed as previously described. For each cell function, we focused on the B cell subtype(s) exhibiting the clearest gradient in activity scores: naive B cells for response to multicellular organisms, intermediate B cells for activation of NK cells, and a combined group of intermediate and memory B cells for protein localization to membrane.

Lead eTRs per gene were selected by the smallest nominal p-value, and gene-level p-values were computed using ACAT-V^74^. Multiple testing across genes was controlled within each bin using the Storey q-value procedure^75^ (FDR < 5%). Cell state-dependent sc-eTRs were identified by comparing high vs. low activity bins using a z-test, with significance defined at Benjamini-Hochberg correction FDR < 5% (**Supplementary Note 4**).

### Fine-mapping and conditional analysis

Prior to fine-mapping, we identified and removed indels that represent TR alleles, using a procedure similar to that described in Margoliash et al^11^. We excluded indels that were 1) found in TR intervals (as defined in our catalog) and 2) composed of whole copies of the TR motif, including its cyclical representations. Conservatively, we retained indels that were not perfect multiples of the TR motif and also retained indels where at least one base would represent a sequence impurity in the TR interval.

We fine-mapped each eGene passing multiple testing correction (FDR < 5%) using SuSiE (v0.12.35)^77^. The fine-mapping region for each gene was defined as the window extending 100 kilobases (kb) upstream of the transcription start site (TSS) and 100 kb downstream of the transcription end site (TES). Fine-mapping was performed using individual-level data from both cohorts jointly to obtain a unified set of credible sets per locus.

As input to SuSiE, we used residualized pseudobulk gene expression values and residualized summed genotype dosages. Residualization was performed by regressing out age, sex, and joint expression and genotype principal components computed across both cohorts. SuSiE was run under a multiple-causal variant assumption, allowing up to 10 credible sets. We used SuSIE’s posterior inclusion probability (PIP) with non-default parameter prune_by_cs set to TRUE, as per Margoliash et al^11^. We conservatively considered an sc-eTR to be candidate causal if it had a PIP of at least 0.7.

We then performed conditional analysis on every candidate causal sc-eTR by performing association testing as previously described, conditioned on the summed repeat dosages of the candidate causal sc-eTR. To account for selection, we controlled the average within-gene FDR using the Benjamini-Bogomolov procedure: for R selected genes out of m tested in discovery, we applied Benjamini-Hochberg within each gene at level α×R/m (α=0.05).

### Functional and genomic annotation of eTRs

We used publicly accessible resources to annotate sc-eTRs, including ENCODE (v3) *cis* candidate regulatory elements (cCREs) (https://hgdownload.soe.ucsc.edu/gbdb/hg38/encode3/ccre/encodeCcreCombined.bb), MANE (v1.0) (https://ftp.ncbi.nlm.nih.gov/refseq/MANE/MANE_human/),TR PhEWAS hits^13^ (Manigbas *et al.*,Table S3), activity by contact scores^78^, ChromHMM^79^, and Zoonomia^44^, and tracks from UCSC Genome Browser^80^.

We harmonized the TR locus definitions in the TR PheWAS catalog with those in our TR catalog by verifying, for each pair of intersecting locus definitions, whether one motif was the reverse complement and/or cyclical shift of the other. HOMER (v5.1) analysis^40^ was run using findMotifsGenome.pl hg38 -size 50

### Methylation analysis

Site-level CpG methylation probabilities were obtained using pb-CpG-tools (v2.3.2) for CpG sites genome-wide using PacBio HiFi sequencing data of 25 individuals in Phase 1 of TenK10K (**Supplementary Fig. 35-36**). CpG sites with methylation standard deviation >0.02 were retained, following a similar procedure described in Trujillo *et al*.^10^

For TRs within 5 kb of a CpG site, TR-methylation associations were tested using associaTR (TRTools v5.0.2)^63^ via ordinary least squares regression of summed TR repeat length on methylation probability, adjusting for sex, age, and the top three genotype PCs.

### Colocalization

We performed colocalization with two types of catalogs: (1) SNV-only GWAS catalogs for asthma^81^, autoimmune conditions (Crohn’s disease^82^, IgA nephropathy^83^, inflammatory bowel disease [IBD]^82^, rheumatoid arthritis^84^, systemic lupus erythematosus^85^, type 1 diabetes^86^, ulcerative colitis^82^), cancer (breast^87^, lung^88^, lymphoma^89^, prostate^90^), COVID-19^91^, and neuroinflammatory conditions (Alzheimer’s^92^, multiple sclerosis^93^, Parkinson’s disease^94^); and (2) joint SNV and imputed STR GWAS catalogs for 44 UKBB blood serum and full blood count traits^11^.

Colocalization was performed using coloc.abf (coloc v5.2.3)^49^. SNV-only colocalization was performed using SNV GWAS summary statistics and sc-eSNVs in our dataset with eQTL type set to ‘quant’ and sdY = 1. For TR and SNV joint colocalization, TR locus definitions were harmonized across datasets, accounting for reverse complements and cyclical motif shifts.

Linkage disequilibrium was calculated as the squared Pearson correlation of summed TR dosages or SNV non-reference allele dosages.

## Extended Data Figures

**Extended Data Figure 1:**
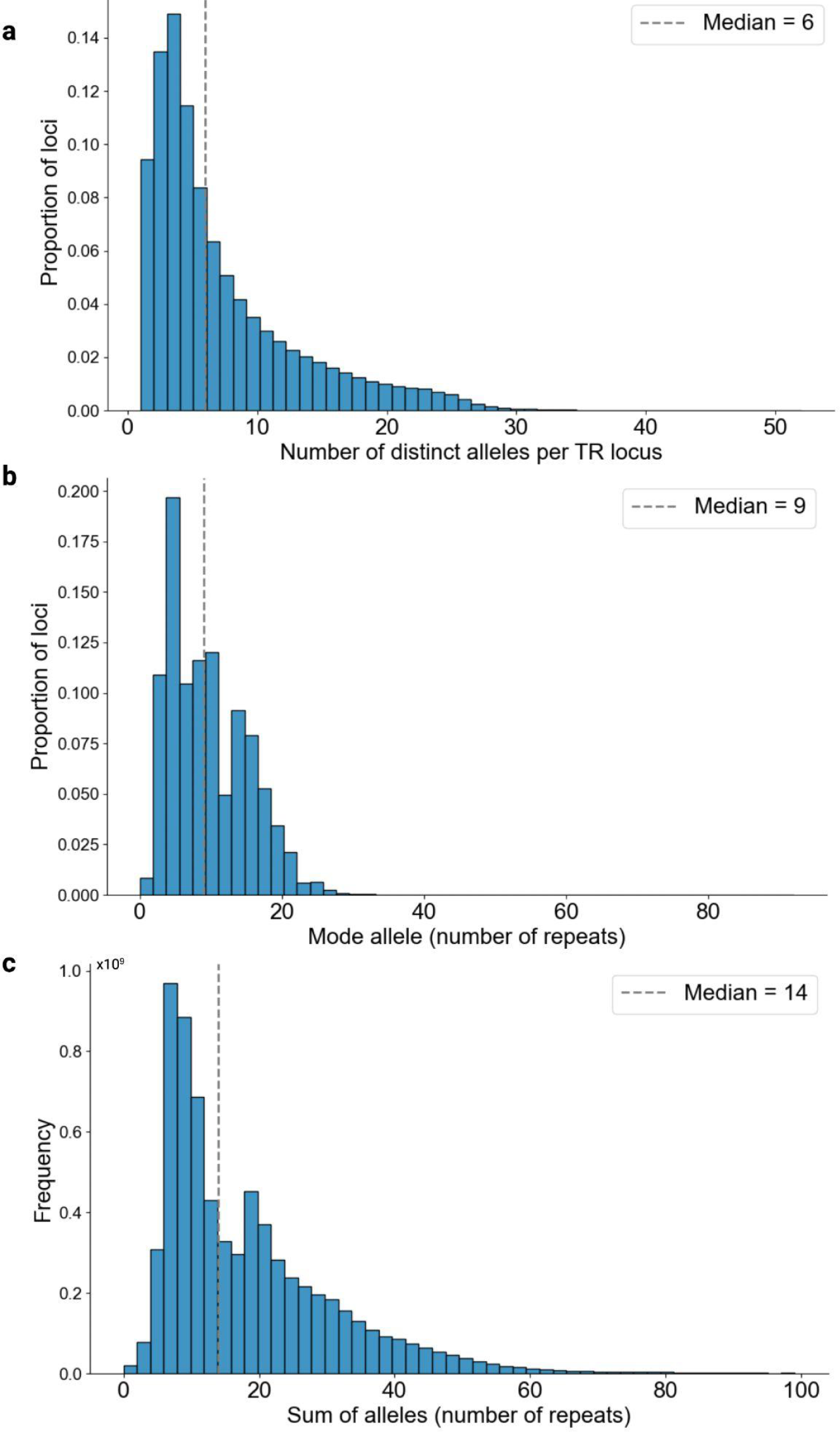
Distribution of tandem repeat genotype characteristics. **a**, Distribution of number of distinct alleles per locus. **b**, Distribution of mode allele per locus. **c**, Distribution of summed alleles (of a diploid genotype). X-axis truncated at 100 repeats. Overall, 13.7% of alleles represented expansions relative to the locus-specific mode, 9.3% were contractions, and the remainder matched the mode allele exactly. The median summed dosage across all loci was 14 repeat units. Following stringent filtering for call quality and locus-level metrics (**Methods**), the resulting high-confidence callset had a median of 6 distinct alleles per locus and a median mode allele length of 9 repeats.

**Extended Data Figure 2:**
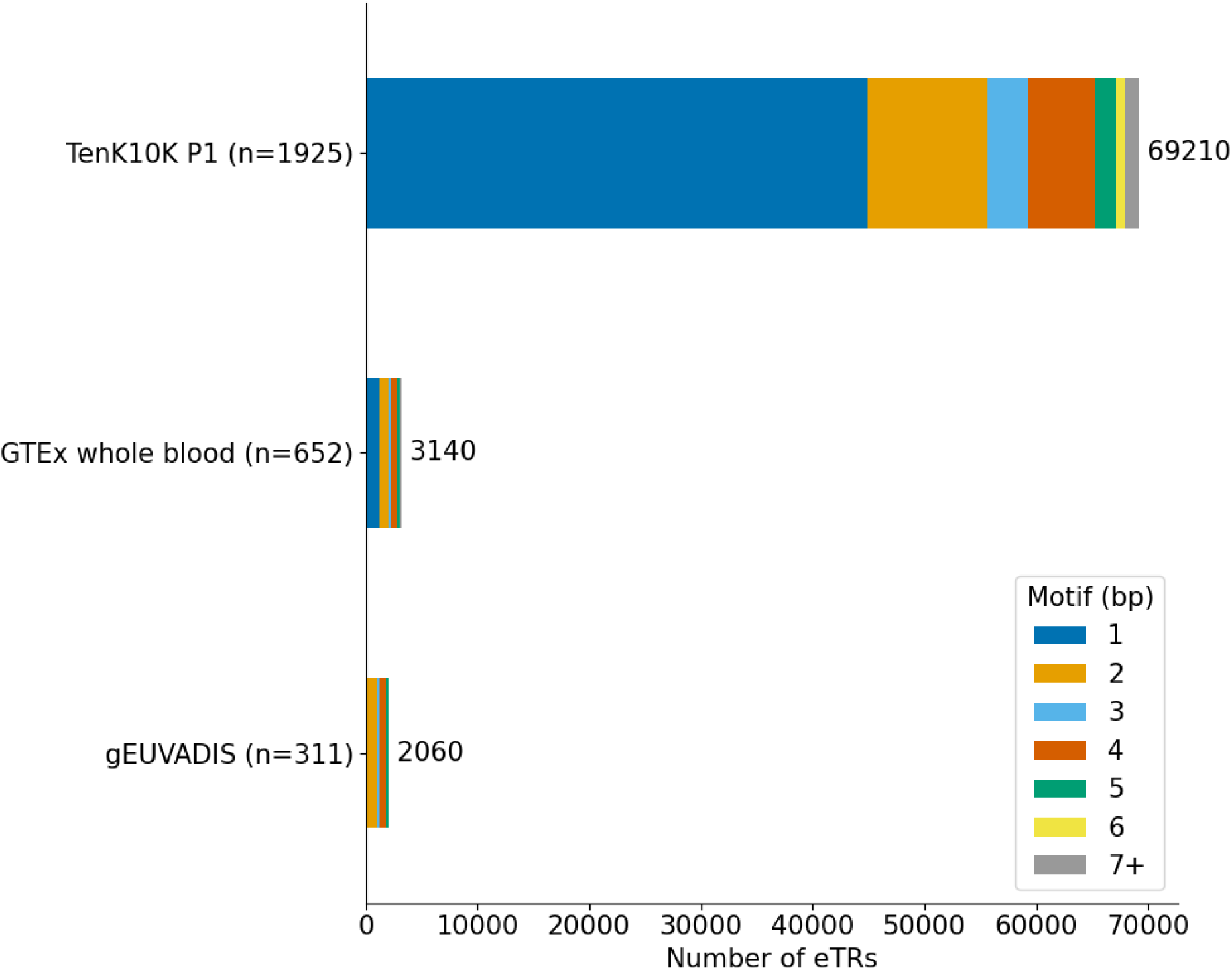
TR eQTL (eTR) discovery in TenK10K Phase 1 (FDR < 5%) compared to two publicly available bulk eTR catalogs. Bulk eTR catalogs were based on GTEx^9,34^ (bulk RNA-seq of whole blood; FDR < 10% for STRs and < 5% for VNTRs) and gEUVADIS^58^ (bulk RNA-seq of HapMap lymphoblastoid cell lines; FDR < 5%) datasets. Differences in cell type/tissue resolution, TR genotyping methods, and analysis pipelines limit interpretability of direct comparisons. In particular, cell type-specific signals discovered in TenK10K Phase 1 may be diluted in bulk tissue, and TR catalogs used in prior studies may not capture loci genotyped in TenK10K Phase 1.

**Extended Data Figure 3:**
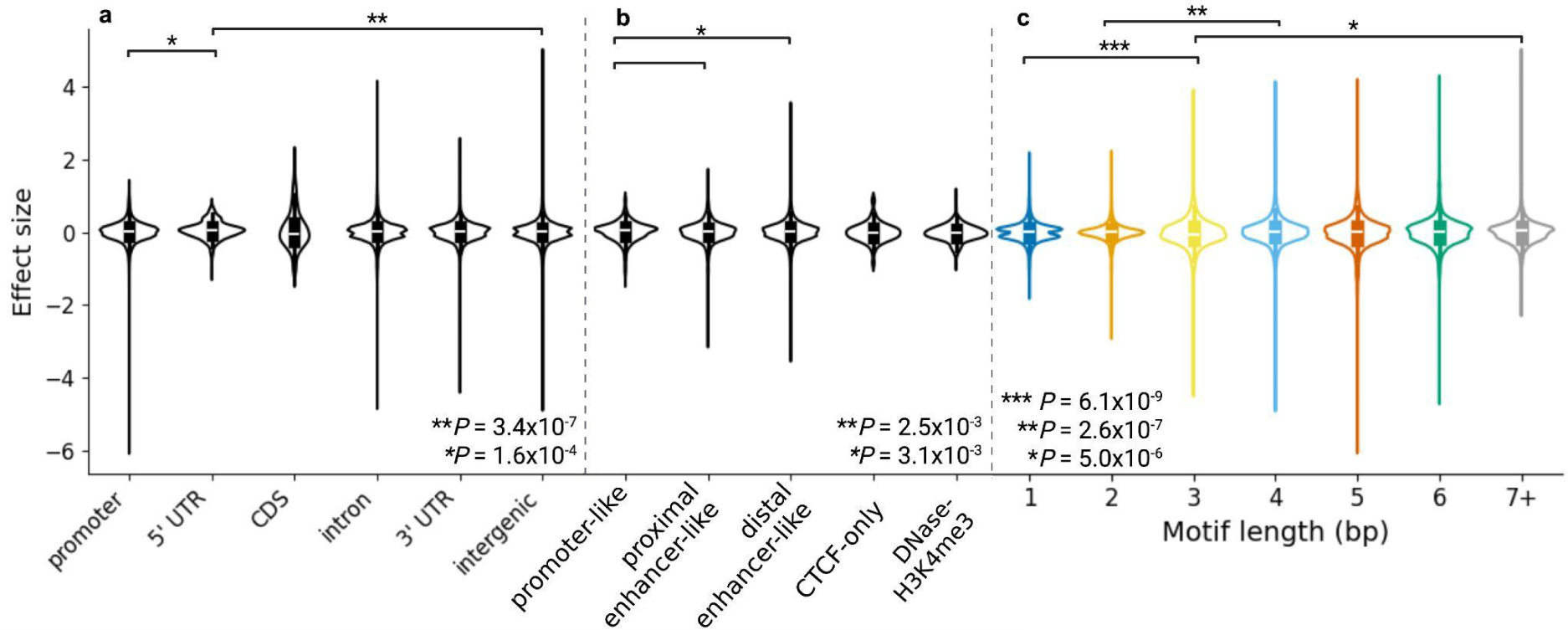
Distributions of sc-eTR effect sizes by genomic annotation. Distributions of sc-eTR effect sizes stratified by MANE gene annotation (a), ENCODE candidate *cis*-regulatory element (cCRE) annotation (b), and motif length (c). Violin plots show the distribution; centre lines indicate medians, boxes indicate interquartile ranges (IQRs), and whiskers extend to 1.5 x IQR. MANE annotations: 5′ untranslated region (5′ UTR), coding sequence (CDS), and 3′ untranslated region (3′ UTR). unadjusted P values were calculated using two-sided Mann–Whitney U tests (n = 112,100 sc-eTRs). In a, \*\**P* = 3.4 x 10^-7^ and \**P* = 1.6 x 10^-4^; in b, \*\**P* = 2.5 x 10^-3^ and \**P* = 3.1 x 10^-3^; in c, \*\*\**P* = 6.1 x 10^-9^, \*\**P* = 2.6 x 10^-7^, and \**P* = 5.0 x 10^-6^.

**Extended Data Figure 4:**
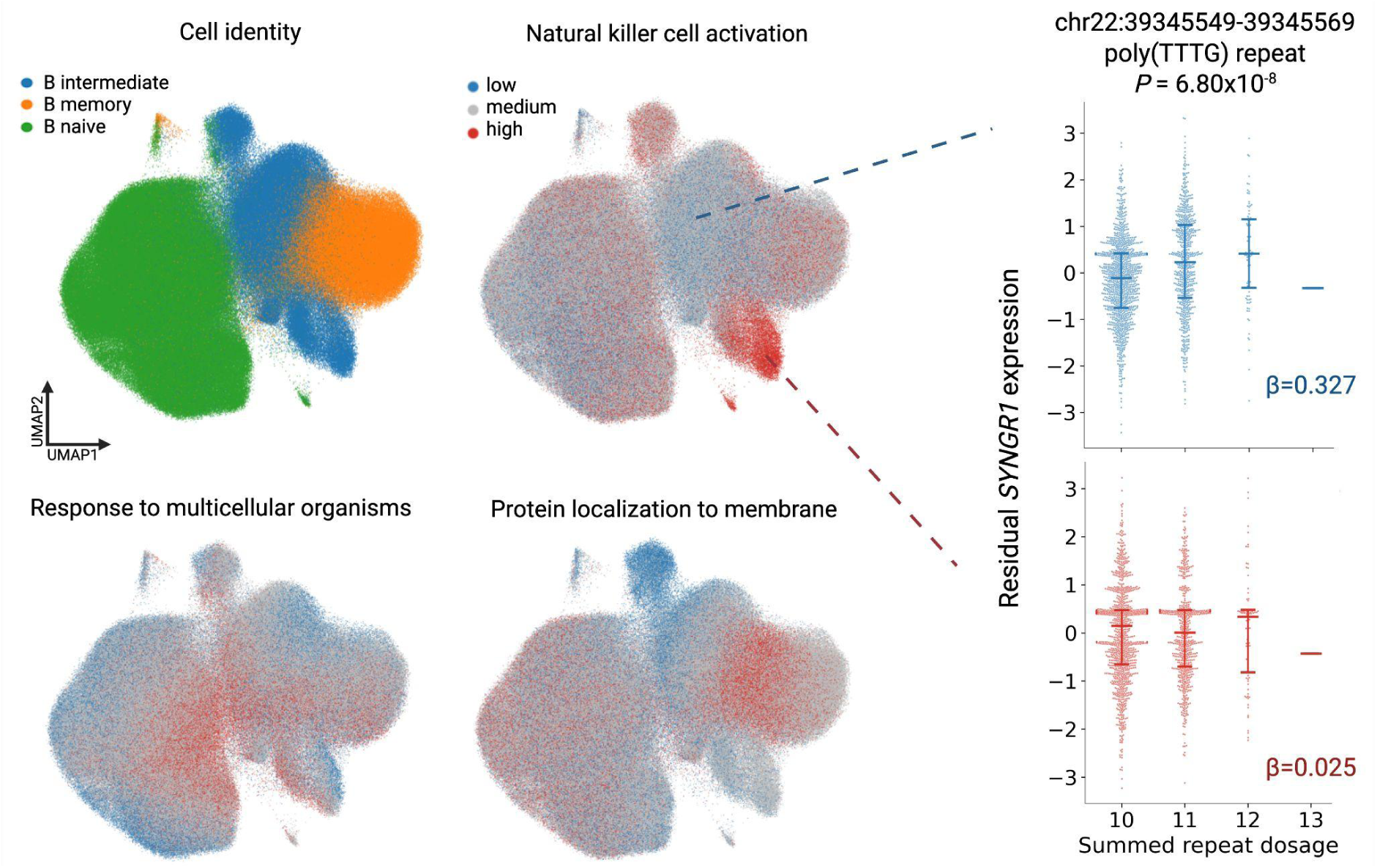
TR cell state inference. UMAP plots of B cells (n = 539,820 cells) first labeled by cell identity, then subsequently labeled by low (15th percentile), medium (between 15th and 85th percentile) and high (above 85th percentile) activity scores for cell functions of natural killer cell activation, response to multicellular organisms, and protein localization to membrane. Swarmplots demonstrate a cell state-specific poly(TTTG) sc-eTR associating with *SYNGR1* expression across the natural killer cell activation axis in n=1,925 individuals. Horizontal bars indicate the first quartile, median, and third quartile. P values were calculated using two-sided Wald tests of the regression coefficient (β) and are uncorrected for multiple testing.

**Extended Data Figure 5:**
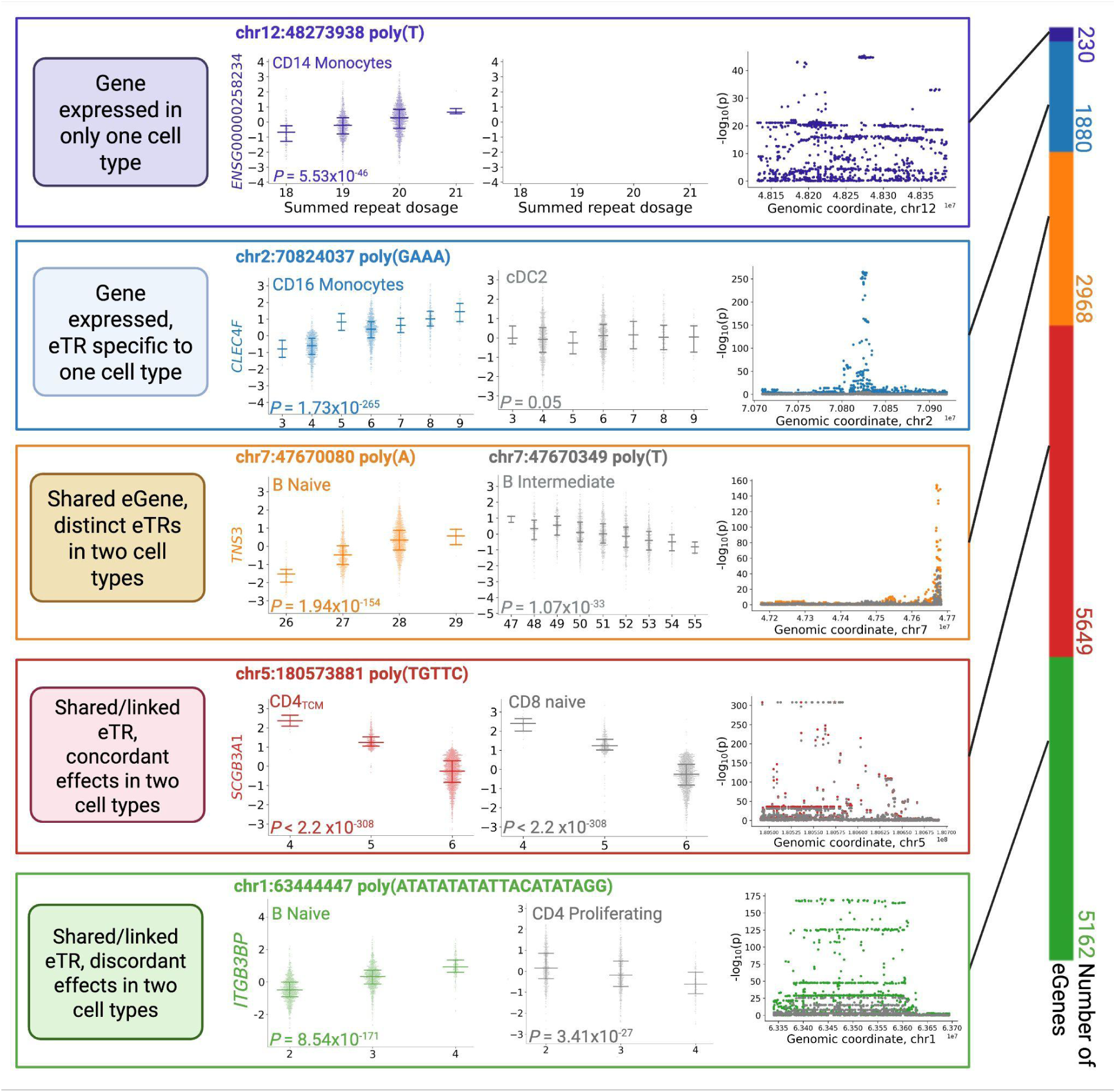
Examples of sc-eTRs illustrating the five scenarios of cell type specificity between two cell types. (cell type A in each scenario’s colour, cell type B in gray). For each scenario, from left to right: brief description of the scenario, association swarm plots for the lead TR in each of the two cell types (n = 1,925 individuals; horizontal bars indicate the first quartile, median, and third quartile), association plots of all tested variants around the gene, with signals for both cell types overlaid. P values were calculated using two-sided Wald tests of the regression coefficient (β) and are uncorrected for multiple testing. The bar chart at the end summarizes each scenario at the level of eGenes, using the maximum from all pairwise cell type comparisons.

**Extended Data Figure 6:**
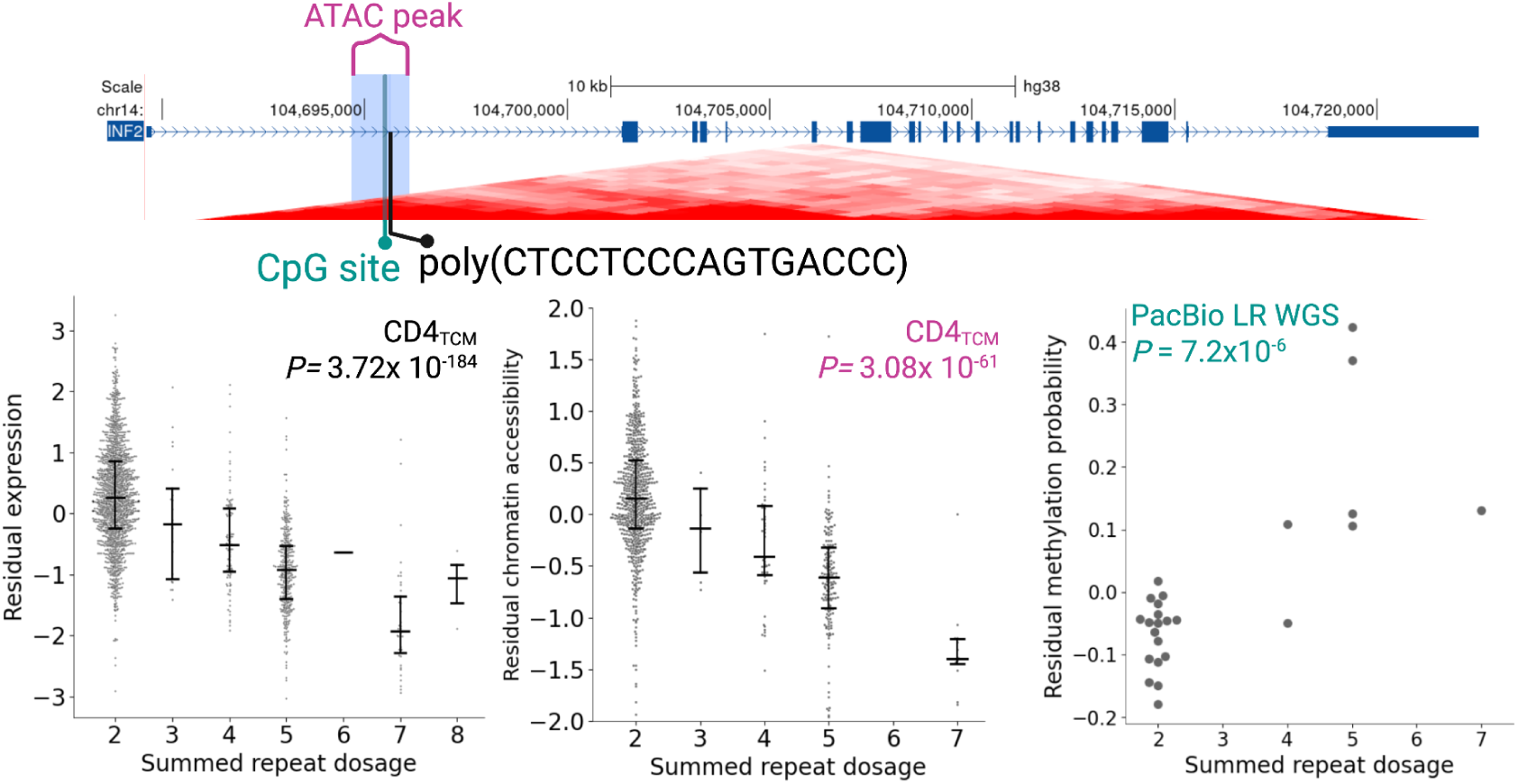
Locus zoom plot and accompanying swarmplots for an intronic poly(CTCCTCCCAGTGACCC) repeat in *INF2* in CD4_TCM_. Swarmplots use scRNA-seq data (n = 1,925 individuals), scATAC-seq data (n = 922 individuals), and PacBio long-read sequencing methylation data (n = 25 individuals). Horizontal bars indicate the first quartile, median, and third quartile. P values were calculated using two-sided Wald tests of the regression coefficient (β) and are uncorrected for multiple testing. Longer alleles at this TR locus are associated with decreased gene expression, decreased chromatin accessibility, and increased methylation.

**Extended Data Figure 7:**
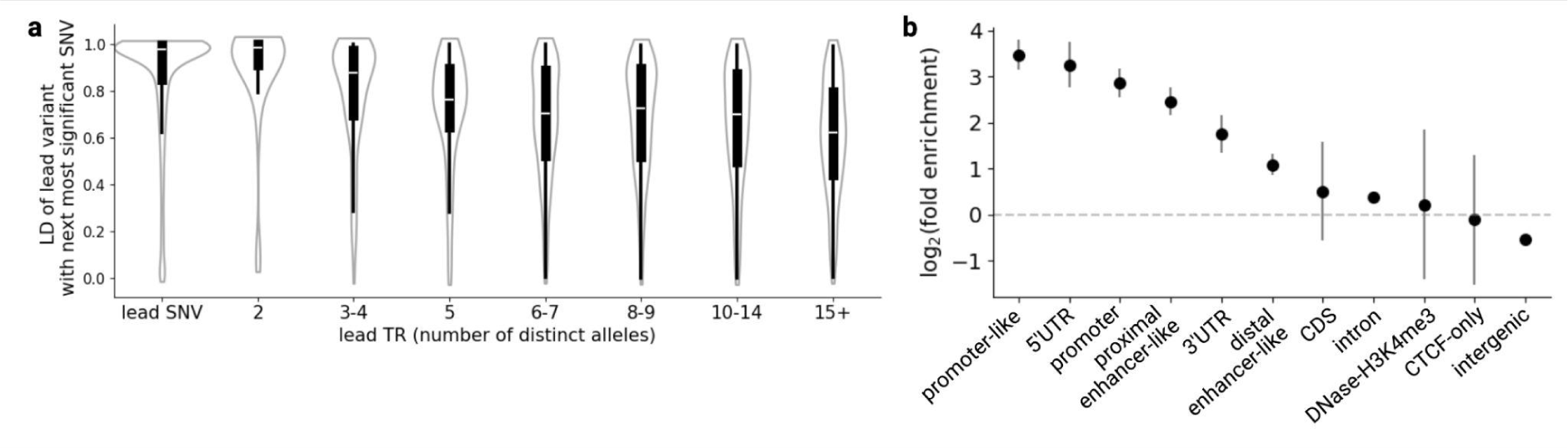
Characterization of lead sc-eTRs. **a**, Linkage disequilibrium (LD) between lead TRs or single nucleotide variants (SNVs) and their next most significant neighboring SNV within ±100 kb of the gene body, in the best-powered cell type (n = 13,093 genes). Violin plots depict the distribution of LD, highlighting median values (white horizontal bar), interquartile ranges (IQR; boxes indicate 25th to 75th percentiles), and whiskers extending to ±1.5×IQR. Lead TRs are grouped into equal-sized bins based on the number of distinct alleles genotyped. **b**, Functional enrichment of candidate causal sc-eTRs (n = 1,490 TRs) relative to all genotyped TRs. Observed log₂ fold enrichment (center point) is shown with 95% confidence intervals. CDS, coding sequence.

**Extended Data Figure 8:**
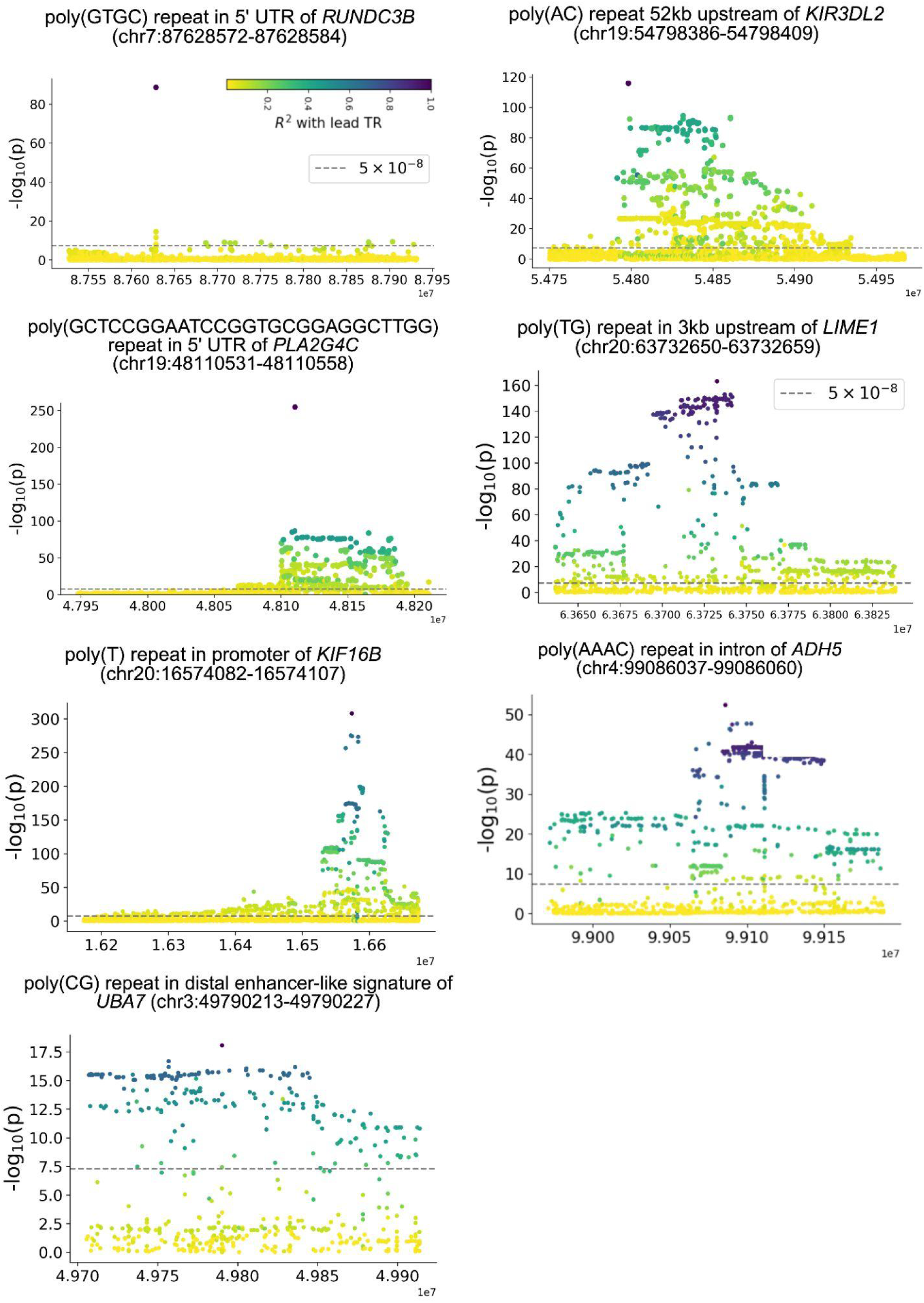
Regional linkage disequilibrium (LD) plots relative to the lead TR for case examples discussed in. Figure 4 **and Extended Data Figures 9-12.** P values were calculated using two-sided Wald tests of the regression coefficient (β) and are uncorrected for multiple testing. Coordinates of the lead TR are provided for each example.

**Extended Data Figure 9:**
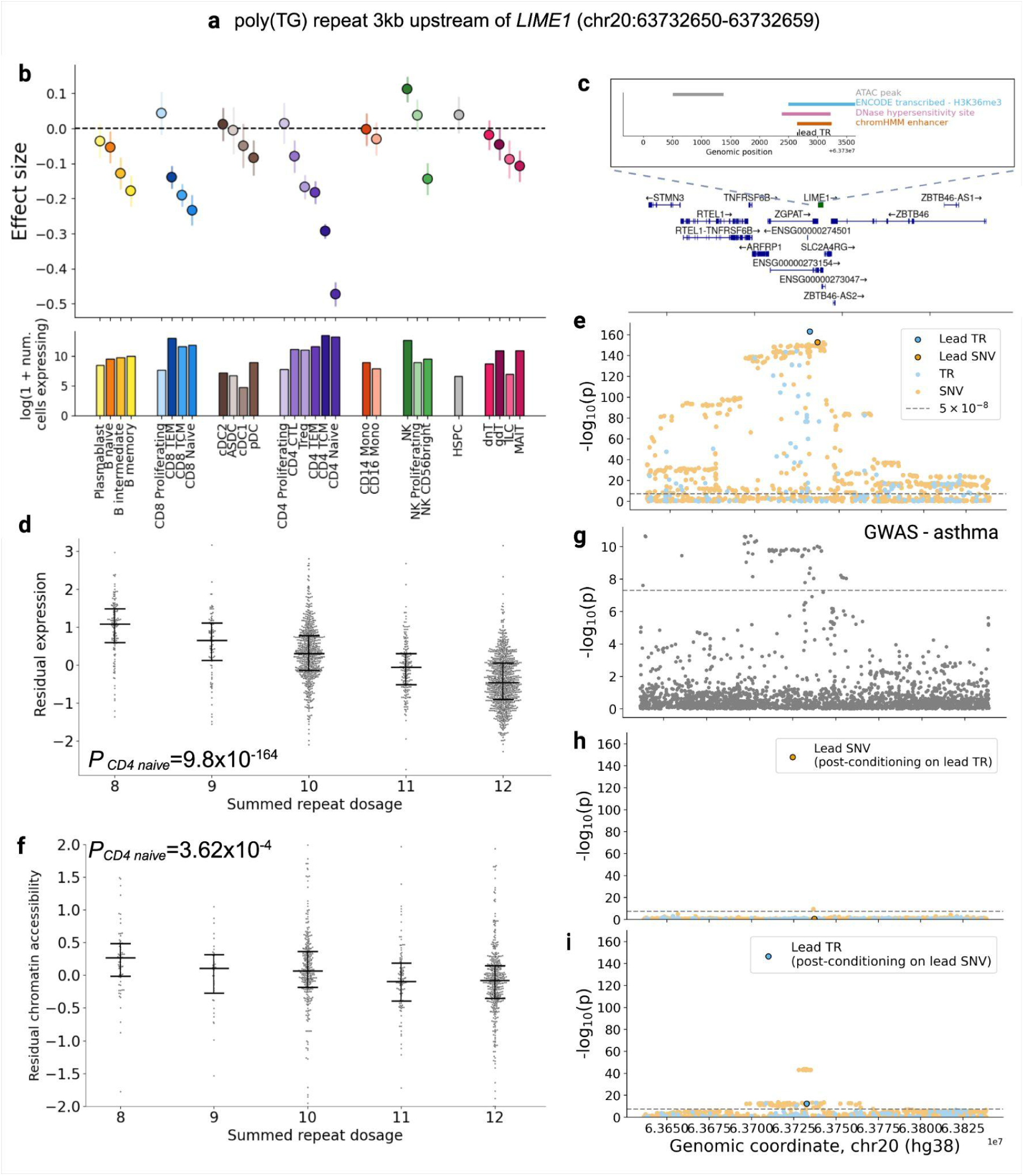
poly(TG) repeat 3kb upstream of LIME1 is a candidate causal variant at a locus colocalizing with asthma. a, Coordinates of the poly(TG) repeat. **b**, Distribution of observed effect sizes (center point) of the sc-eTR in immune cell types which had sufficient gene expression for association testing (top panel) (**Methods**). Error bars correspond to 95% confidence intervals. Bar plot of number of cells expressing *LIME1* by cell type (bottom panel). Cell counts are derived from a subset of a total dataset comprising *N* = 5,438,679 cells. **c**, Locus zoom plot of *LIME1* and overlapping annotation tracks (**Methods**). ATAC peak annotations reflect the ATAC peak mapped in **f**. **d,** Association between repeat length and residual gene expression in CD4 naive cells (n=1,925 individuals). Horizontal bars indicate the first quartile, median, and third quartile. **e**, sc-eQTL association signals for the candidate causal sc-eTR and other SNVs and TRs within +/-100kb of the gene body. **f,** Association between repeat length and residual chromatin accessibility (chr20:63730505-63731380) in CD4 naive cells (n=922 individuals). Horizontal bars indicate the first quartile, median, and third quartile. **g,** Asthma GWAS association signals^81^ for the equivalent window (+/- 100kB of the gene body of *LIME1*). **h,** sc-eQTL association signals for variants within +/- 100kb of the gene body, after conditioning on the genotype of the lead TR. **i,** sc-eQTL association signals for variants within +/- 100kb of the gene body, after conditioning on the genotype of the lead SNV. P values were calculated using two-sided Wald tests of the regression coefficient (β) and are uncorrected for multiple testing in **d-f** and **h-i**.

**Extended Data Figure 10:**
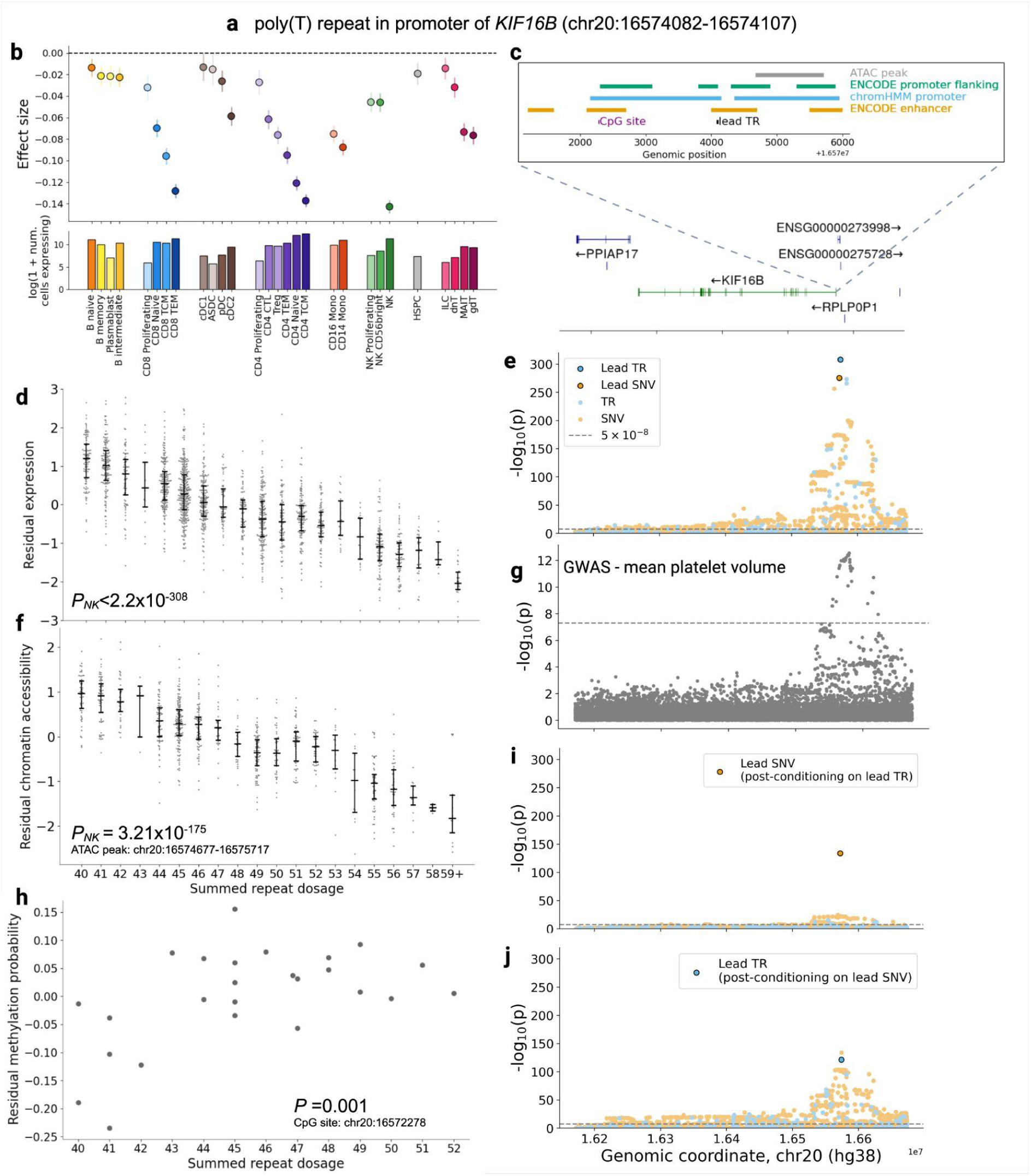
poly(T) repeat in the promoter of *KIF16B* is located within a locus that colocalizes with GWAS for mean platelet volume. a, Coordinates of the poly(T) repeat. **b**, Distribution of observed effect sizes (center point) for the sc-eTR across immune cell types with sufficient gene expression for association testing, with error bars indicating 95% confidence intervals (top panel). The bottom panel shows the number of cells expressing *KIF16B* per cell type. Cell counts are derived from a subset of a total dataset comprising *N* = 5,438,679 cells. **c**, Locus zoom plot of *KIF16B* and overlapping annotation tracks (**Methods**). ATAC-peak and CpG site annotations reflect coordinates in **f** and **h**. **d,** Association between repeat length and residual gene expression in NK cells (n=1,925 individuals). Horizontal bars indicate the first quartile, median, and third quartile. **e**, sc-eQTL association signals for the candidate causal sc-eTR and other SNVs and TRs within ±100kb of the gene body. **f,** Association between repeat length and residual chromatin accessibility in NK cells (n=922 individuals). Horizontal bars indicate the first quartile, median, and third quartile. **g,** Mean platelet volume GWAS association signals^11^ for the equivalent window (±100kB of the gene body of *KIF16B*). **h,** Association between repeat length and residual methylation probability derived from PacBio long-read sequencing (n=25). **i,** sc-eQTL association signals conditioned on the genotype of the candidate causal poly(T) repeat. **j,** sc-eQTL association signals conditioned on the genotype of the lead SNV. P values were calculated using two-sided Wald tests of the regression coefficient (β) and are uncorrected for multiple testing in **d-f** and **h-j**.

**Extended Data Figure 11:**
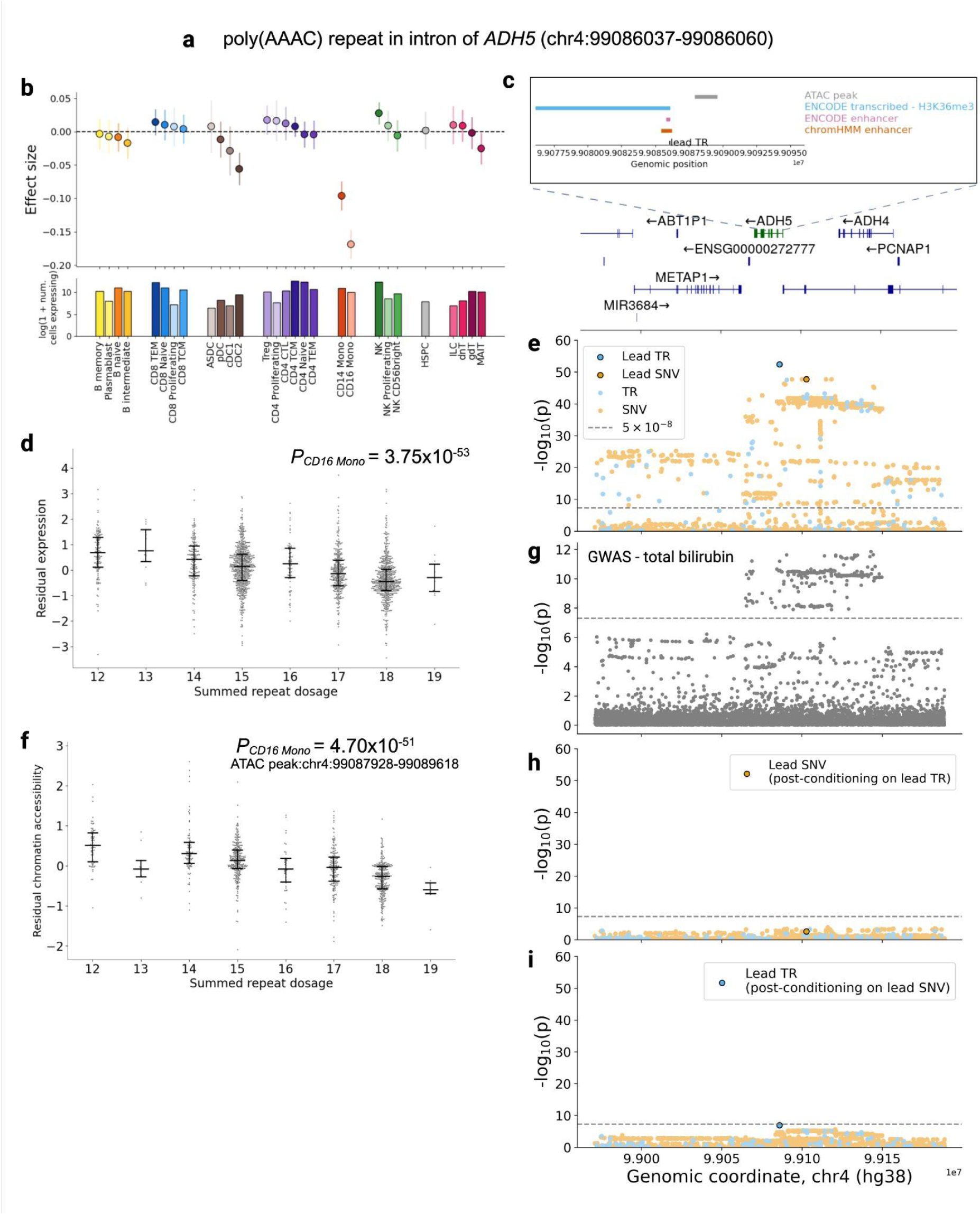
poly(AAAC) repeat in the intron of *ADH5* is located within a locus that colocalizes with GWAS for total bilirubin a, Coordinates of the poly(AAAC) repeat. **b**, Distribution of observed effect sizes (center point) of the sc-eTR in immune cell types which had sufficient gene expression for association testing (top panel) (**Methods**). Error bars correspond to 95% confidence intervals. Bar plot of number of cells expressing *ADH5* by cell type (bottom panel). Cell counts are derived from a subset of a total dataset comprising *N* = 5,438,679 cells. **c**, Locus zoom plot of *ADH5* and overlapping annotation tracks (**Methods**). ATAC-peak annotations reflect coordinates in **f**. **d,** Association between repeat length and residual gene expression in CD16 monocyte cells (n=1,925 individuals). Horizontal bars indicate the first quartile, median, and third quartile. P values were calculated using two-sided Wald tests of the regression coefficient (β) and are uncorrected for multiple testing. **e**, sc-eQTL association signals for the candidate causal sc-eTR and other SNVs and TRs within +/- 100kb of the gene body. **f,** Association between repeat length and residual chromatin accessibility in CD16 monocyte cells (n=922 individuals). Horizontal bars indicate the first quartile, median, and third quartile. **g,** Total bilirubin GWAS association signals^11^ for the equivalent window (+/- 100kB of the gene body of *ADH5*). **h,** sc-eQTL association signals for variants within +/- 100kb of the gene body, after conditioning on the genotype of the lead TR. **i,** sc-eQTL association signals for variants within +/- 100kb of the gene body, after conditioning on the genotype of the lead SNV. P values were calculated using two-sided Wald tests of the regression coefficient (β) and are uncorrected for multiple testing in **d-f** and **h-i**.

**Extended Dat Figure 12:**
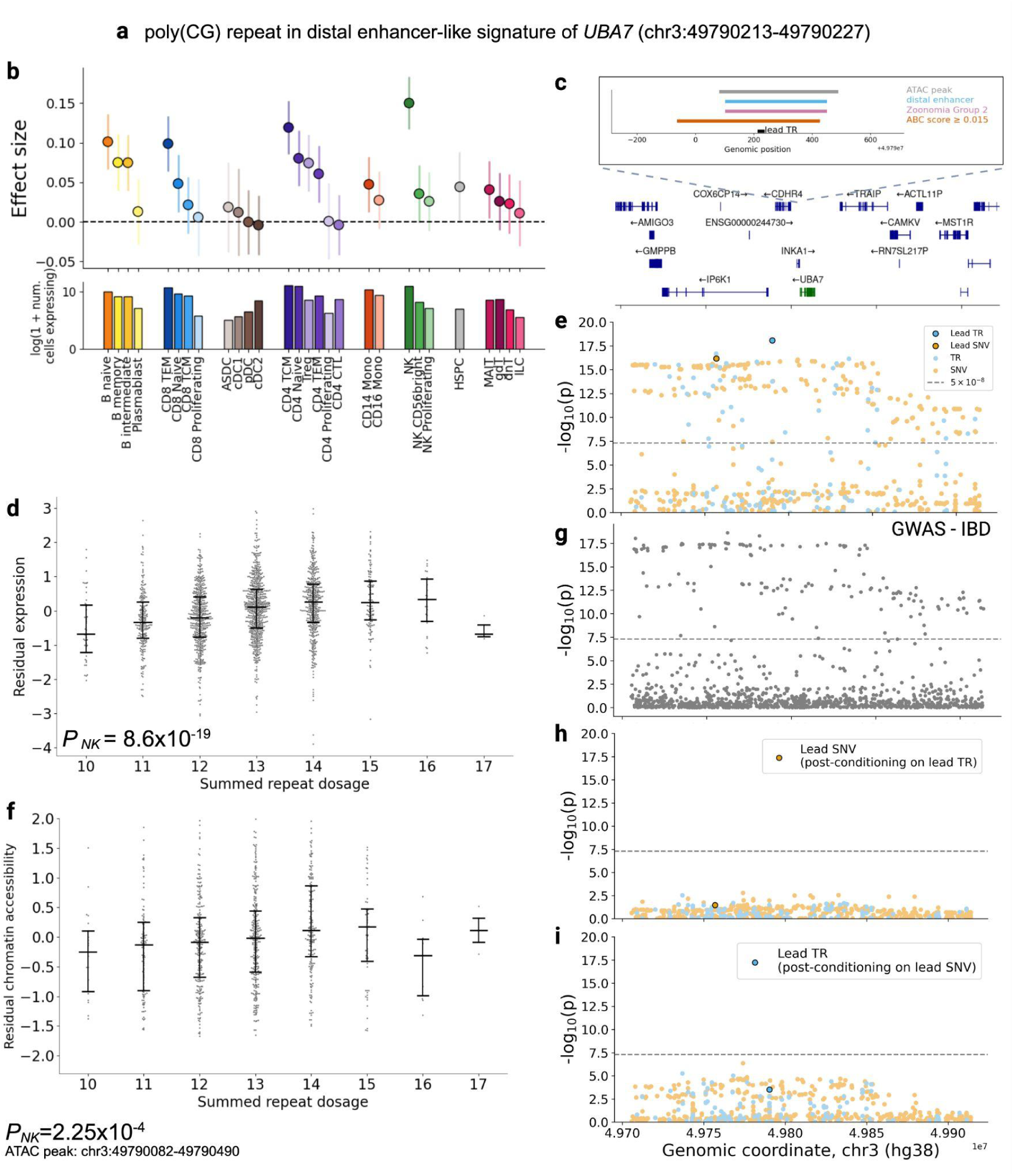
poly(CG) repeat associated with *UBA7* expression is located within a locus that colocalizes with GWAS for inflammatory bowel disease a, Coordinates of the poly(CG) repeat. **b**, Distribution of observed effect sizes (center point) of the sc-eTR in immune cell types which had sufficient gene expression for association testing (top panel) (**Methods**). Error bars correspond to 95% confidence intervals. Bar plot of number of cells expressing *UBA7* by cell type (bottom panel). Cell counts are derived from a subset of a total dataset comprising *N* = 5,438,679 cells. **c**, Locus zoom plot of *UBA7* and overlapping annotation tracks (**Methods**). ATAC-peak annotations reflect coordinates in **f**. **d,** Association between repeat length and residual gene expression in NK cells (n=1,925 individuals). Horizontal bars indicate the first quartile, median, and third quartile. **e**, sc-eQTL association signals for the candidate causal sc-eTR and other SNVs and TRs within +/- 100kb of the gene body. **f,** Association between repeat length and residual chromatin accessibility in NK cells (n=922 individuals). Horizontal bars indicate the first quartile, median, and third quartile. **g,** Inflammatory bowel disease GWAS association signals^82^ for the equivalent window (+/- 100kB of the gene body of *UBA7*). **h,** sc-eQTL association signals for variants within +/- 100kb of the gene body, after conditioning on the genotype of the lead TR. **i,** sc-eQTL association signals for variants within +/- 100kb of the gene body, after conditioning on the genotype of the lead SNV. P values were calculated using two-sided Wald tests of the regression coefficient (β) and are uncorrected for multiple testing in **d-f** and **h-i**.

## Acknowledgements

We acknowledge Vladislav Savelyev for contributions in prototyping the whole-genome sequencing production pipeline, and Garvan Genomics Platform for providing sequencing services and technical support. We acknowledge Leonard Goldstein, Michael Geaghan, and Matthew Hobbs from the Garvan Data Science Platform for support. The costs of WGS for most samples in this program were supported by the Centre for Population Genomics. Illumina provided discounted reagents for scRNA-seq, and 10X Genomics provided discounted reagents for single cell capture. Data generation and analysis was supported in part by the Centre for Population Genomics, with funding from Garvan Institute of Medical Research, Murdoch Children’s Research Institute, and a National Health and Medical Research Council (NHMRC) investigator grant (2009982). The contents of this manuscript are solely the responsibility of the authors and do not reflect the views of the NHMRC. H.D. is supported by NHGRI grant 4R00HG012796-03 and NHMRC Investigator grant GNT2026126. J.M.P. was supported by a Gates Cambridge fellowship (OPP1144). M.G. and J.Marg are supported by NHGRI grant R01HG010885. G.A.F. and the BioHEART Study have been supported by NHMRC Investigator Grant, NSW Health Office of Health and Medical Research, and the NSW Health Statewide Biobank scheme.

## Author contributions

H.A.T, A.S.E.C., and D.G.M. designed the study. H.A.T. designed and performed the tandem repeat eQTL and downstream analysis, and led the writing of the manuscript. B.W., H.A.T, N.E.K., and H.B. curated the genome-wide tandem repeat catalog and provided guidance on tandem repeat genotyping. M.W. and M.F. contributed to the tandem repeat eQTL pipeline. E.S. and C.B. designed and performed the single-cell RNA sequencing experiments. R. A. M developed the ethics protocol and contributed to the design of the single-cell RNA sequencing experiments. A.S.E.C and B.B. processed the single-cell RNA sequencing data. A.X., A.S., and P.A. contributed to the single-cell RNA analysis. A.X., J.F, and O.A.D. designed the single-cell ATAC sequencing experiments and processed the raw data. K.W., O.T., M.G., S.M.G., and T.N. assisted with participant recruitment and sample collection. A.M. and B.B. designed and led the submission of the publicly accessible datasets. A.L.M.R performed long-read tandem repeat genotyping. J.Marg. provided guidance on fine-mapping. N.E.K and H.B. provided guidance on optimizing genome-wide tandem repeat genotyping. J.M.P. provided guidance on colocalization and fine-mapping. A.S.L contributed to the functional interpretation of fine-mapped results. M.H., K.B., and M.S processed the whole genome sequencing data. J.Mars., V.B., and L.G. contributed to the cloud computing workflow. B.S.M. and C.U. developed the ethics protocol for the study and coordinated the generation of WGS data. H.D. and L.F. provided guidance on tandem repeat genotyping tools. A.M-T contributed to the methylation analysis. E.D. contributed to the analysis of the long-read tandem repeat genotypes. Z.Q., O.M.S., M.T., S.I.A, H.R.N, K.K-H.F., E.B-D., and L.C. contributed to the study design and discussion of analysis of results. J.P. and K.M.dL led the processing of whole genome sequencing data. C.S. provided guidance on interpreting whole genome sequencing data. C.W. provided guidance on fine-mapping, colocalization and study design. M.G. provided guidance on association analysis and study design. I.W.D contributed to long-read sequencing and tandem repeat genotyping. A.W.H designed the Tasmanian Ophthalmic Biobank study. G.A.F designed the BioHEART study.

All authors contributed to editing the manuscript. K.M.dL, J.E.P and D.G.M conceived the project and supervised all aspects of the work.

## Additional Information

Supplementary Information is available for this paper.

Correspondence and requests for materials should be addressed to Hope Tanudisastro or Daniel G. MacArthur.

## Data availability

Genomic coordinates and variant annotations were defined relative to the human reference genome GRCh38 (Genome Reference Consortium Human Build 38), obtained from the Genome Reference Consortium (https://www.ncbi.nlm.nih.gov/grc/human). Gene annotation was performed using GENCODE v44/Ensemble 110. sc-eTRs were annotated with ENCODE (v3) cis candidate regulatory elements (cCREs) (https://hgdownload.soe.ucsc.edu/gbdb/hg38/encode3/ccre/encodeCcreCombined.bb) and MANE (v1.0) (https://ftp.ncbi.nlm.nih.gov/refseq/MANE/MANE_human/).

Summary-level association statistics are provided in the Supplementary Tables. Raw summary association statistics and the polymorphic variant catalog are available at Zenodo (https://doi.org/10.5281/zenodo.15009519). Individual-level whole-genome sequencing (WGS) data and genotypes are available through the European Genome-Phenome Archive (EGA) under accession EGAS50000001654, under controlled access for non-commercial research use. Single-cell RNA sequencing (scRNA-seq) data are available under EGA accession EGAS50000001653, under controlled access for non-commercial research use. Summary-level tandem repeat (TR) genotypes and association statistics for sc-eTRs are accessible at trexplorer.broadinstitute.org.

## Code availability

Single-cell expression processing and downstream analyses is available at https://github.com/powellgenomicslab/tenk10k_phase1.

WGS data processing pipeline is available at https://github.com/populationgenomics/production-pipelines/tree/main/cpg_workflows/large_cohort.

TR genotyping, QC, and association analysis pipeline is available at https://github.com/populationgenomics/sv-workflows/tree/main/str.

TR merging and catalog curation is available at https://github.com/broadinstitute/str-analysis. scDeepID pipeline is available at https://github.com/powellgenomicslab/scDeepID_TenK10K_manuscripts.

## Main text References

1. Tanudisastro, H. A., Deveson, I. W., Dashnow, H. & MacArthur, D. G. Sequencing and characterizing short tandem repeats in the human genome. Nat. Rev. Genet. 25, 460–475 (2024).

2. Horton, C. A. et al. Short tandem repeats bind transcription factors to tune eukaryotic gene expression. Science 381, eadd1250 (2023).

3. Ziaei Jam, H., et al. A deep population reference panel of tandem repeat variation. Nat. Commun. 14, 6711 (2023).

4. Sun, J. X. et al. A direct characterization of human mutation based on microsatellites. Nat. Genet. 44, 1161–1165 (2012).

5. Gent, J. I. et al. Distinct influences of tandem repeats and retrotransposons on CENH3 nucleosome positioning. Epigenetics Chromatin 4, 3 (2011).

6. Vinces, M. D., Legendre, M., Caldara, M., Hagihara, M. & Verstrepen, K. J. Unstable tandem repeats in promoters confer transcriptional evolvability. Science 324, 1213–1216 (2009).

7. Sawaya, S. et al. Microsatellite tandem repeats are abundant in human promoters and are associated with regulatory elements. PLoS One 8, e54710 (2013).

8. Depienne, C. & Mandel, J.-L. 30 years of repeat expansion disorders: What have we learned and what are the remaining challenges? Am. J. Hum. Genet. 108, 764–785 (2021).

9. Fotsing, S. F. et al. The impact of short tandem repeat variation on gene expression. Nat. Genet. 51, 1652–1659 (2019).

10. Martin-Trujillo, A., Garg, P., Patel, N., Jadhav, B. & Sharp, A. J. Genome-wide evaluation of the effect of short tandem repeat variation on local DNA methylation. Genome Res. 33, 184–196 (2023).

11. Margoliash, J. et al. Polymorphic short tandem repeats make widespread contributions to blood and serum traits. Cell Genom. 3, 100458 (2023).

12. Erwin, G. S. et al. Recurrent repeat expansions in human cancer genomes. Nature 613, 96–102 (2023).

13. Manigbas, C. A. et al. A phenome-wide association study of tandem repeat variation in 168,554 individuals from the UK Biobank. Nat. Commun. 15, 10521 (2024).

14. Saini, S., Mitra, I., Mousavi, N., Fotsing, S. F. & Gymrek, M. A reference haplotype panel for genome-wide imputation of short tandem repeats. Nat. Commun. 9, 4397 (2018).

15. Cuomo, A. S. E., Nathan, A., Raychaudhuri, S., MacArthur, D. G. & Powell, J. E. Single-cell genomics meets human genetics. Nat. Rev. Genet. 24, 535–549 (2023).

16. Yazar, S. et al. Single-cell eQTL mapping identifies cell type-specific genetic control of autoimmune disease. Science 376, eabf3041 (2022).

17. Handsaker, R. E. et al. Long somatic DNA-repeat expansion drives neurodegeneration in Huntington’s disease. Cell 188, 623–639.e19 (2025).

18. Benson, G. Tandem repeats finder: a program to analyze DNA sequences. Nucleic Acids Res. 27, 573–580 (1999).

19. Mousavi, N., Shleizer-Burko, S., Yanicky, R. & Gymrek, M. Profiling the genome-wide landscape of tandem repeat expansions. Nucleic Acids Res. 47, e90 (2019).

20. Willems, T. et al. Genome-wide profiling of heritable and de novo STR variations. Nat. Methods 14, 590–592 (2017).

21. Halman, A., Dolzhenko, E. & Oshlack, A. STRipy: A graphical application for enhanced genotyping of pathogenic short tandem repeats in sequencing data. Hum. Mutat. 43, 859–868 (2022).

22. Weisburd, B., Tiao, G. & Rehm, H. L. Insights from a genome-wide truth set of tandem repeat variation. bioRxiv (2023) doi:10.1101/2023.05.05.539588.

23. Weisburd, B. et al. Defining a tandem repeat catalog and variation clusters for genome-wide analyses and population databases. bioRxiv (2024) doi:10.1101/2024.10.04.615514.

24. Steely, C. J., Watkins, W. S., Baird, L. & Jorde, L. B. The mutational dynamics of short tandem repeats in large, multigenerational families. Genome Biol. 23, 253 (2022).

25. Kimura, M. & Ohta, T. Stepwise mutation model and distribution of allelic frequencies in a finite population. Proc. Natl. Acad. Sci. U. S. A. 75, 2868–2872 (1978).

26. Urbut, S. M., Wang, G., Carbonetto, P. & Stephens, M. Flexible statistical methods for estimating and testing effects in genomic studies with multiple conditions. Nat. Genet. 51, 187–195 (2019).

27. Collin, M. & Bigley, V. Human dendritic cell subsets: an update. Immunology 154, 3–20 (2018).

28. Lou, Z., Casali, P. & Xu, Z. Regulation of B cell differentiation by intracellular membrane-associated proteins and microRNAs: role in the antibody response. Front. Immunol 6, 537 (2015).

29. Gyurova, I. E., Ali, A. & Waggoner, S. N. Natural killer cell regulation of B cell responses in the context of viral infection. Viral Immunol. 33, 334–341 (2020).

30. Cyster, J. G. & Allen, C. D. C. B cell responses: cell interaction dynamics and decisions. Cell 177, 524–540 (2019).

31. Thalayasingam, N. et al. CD4+ and B lymphocyte expression quantitative traits at rheumatoid arthritis risk loci in patients with untreated early arthritis: implications for causal gene identification. Arthritis Rheumatol. 70, 361–370 (2018).

32. Schmiedel, B. J. et al. Impact of genetic polymorphisms on human immune cell gene expression. Cell 175, 1701–1715.e16 (2018).

33. Nathan, A. et al. Single-cell eQTL models reveal dynamic T cell state dependence of disease loci. Nature 606, 120–128 (2022).

34. Bakhtiari, M. et al. Variable number tandem repeats mediate the expression of proximal genes. Nat. Commun. 12, 2075 (2021).

35. Damdinsuren, B., Dement-Brown, J., Li, H. & Tolnay, M. B cell receptor induced Fc receptor-like 5 expression is mediated by multiple signaling pathways converging on NF-κB and NFAT. Mol. Immunol. 73, 112–121 (2016).

36. Zhang, X. & Ren, L. Dynamic genetic regulation of CD4 T cells in obstructive sleep apnea: integrating context-specific eQTL, Mendelian randomization, single-cell sequencing, and experimental validation. Front. Immunol. 16, 1691347 (2025).

37. Smith, E. M. Neuropeptides as signal molecules in common with leukocytes and the hypothalamic-pituitary-adrenal axis. Brain Behav. Immun. 22, 3–14 (2008).

38. Andrés-Delgado, L. et al. INF2 promotes the formation of detyrosinated microtubules necessary for centrosome reorientation in T cells. J. Cell Biol. 198, 1025–1037 (2012).

39. Sun, J. H. et al. Disease-associated short tandem repeats co-localize with chromatin domain boundaries. Cell 175, 224–238.e15 (2018).

40. Heinz, S. et al. Simple combinations of lineage-determining transcription factors prime cis-regulatory elements required for macrophage and B cell identities. Mol. Cell 38, 576–589 (2010).

41. Hart, G. T., Hogquist, K. A. & Jameson, S. C. Krüppel-like factors in lymphocyte biology. J. Immunol. 188, 521–526 (2012).

42. Deaton, A. M. & Bird, A. CpG islands and the regulation of transcription. Genes Dev. 25, 1010–1022 (2011).

43. Kuderna, L. F. K. et al. Identification of constrained sequence elements across 239 primate genomes. Nature 625, 735–742 (2024).

44. Andrews, G. et al. Mammalian evolution of human cis-regulatory elements and transcription factor binding sites. Science 380, eabn7930 (2023).

45. Lek, M. et al. Analysis of protein-coding genetic variation in 60,706 humans. Nature 536, 285–291 (2016).

46. Taylor, K. H. et al. Large-scale CpG methylation analysis identifies novel candidate genes and reveals methylation hotspots in acute lymphoblastic leukemia. Cancer Res. 67, 2617–2625 (2007).

47. Shaw, J. & Kollnberger, S. New perspectives on the ligands and function of the killer cell immunoglobulin-like receptor KIR3DL2 in health and disease. Front. Immunol. 3, 339 (2012).

48. Xu, S. et al. Cytosolic phospholipase A2 gamma is involved in hepatitis C virus replication and assembly. J. Virol. 86, 13025–13037 (2012).

49. Giambartolomei, C. et al. Bayesian test for colocalisation between pairs of genetic association studies using summary statistics. PLoS Genet. 10, e1004383 (2014).

50. Martino, D. et al. Epigenetic dysregulation of naive CD4+ T-cell activation genes in childhood food allergy. Nat. Commun. 9, 3308 (2018).

51. Weaver, N. et al. KIF16B mediates anterograde transport and modulates lysosomal degradation of the HIV-1 envelope glycoprotein. J. Virol. 97, e0025523 (2023).

52. Ponomarev, E. D. Fresh evidence for platelets as neuronal and innate immune cells: their role in the activation, differentiation, and deactivation of Th1, Th17, and Tregs during tissue inflammation. Front. Immunol. 9, 406 (2018).

53. Luo, W.-W. et al. Transcription-independent regulation of STING activation and innate immune responses by IRF8 in monocytes. Nat. Commun. 13, 4822 (2022).

54. Li, X.-J.-Y., Qu, J.-R., Zhang, Y.-H. & Liu, R.-P. The dual function of cGAS-STING signaling axis in liver diseases. Acta Pharmacol. Sin. 45, 1115–1129 (2024).

55. Fan, J.-B. et al. Type I IFN induces protein ISGylation to enhance cytokine expression and augments colonic inflammation. Proc. Natl. Acad. Sci. U. S. A. 112, 14313–14318 (2015).

56. Sarafidou, T., Galliopoulou, E., Apostolopoulou, D., Fragkiadakis, G. A. & Moschonas, N. K. Reconstruction of a comprehensive interactome and experimental data analysis of FRA10AC1 may provide insights into its biological role in health and disease. Genes (Basel) 14, (2023).

57. Alquicira-Hernandez, J., Sathe, A., Ji, H. P., Nguyen, Q. & Powell, J. E. scPred: accurate supervised method for cell-type classification from single-cell RNA-seq data. Genome Biol. 20, 264 (2019).

58. Gymrek, M. et al. Abundant contribution of short tandem repeats to gene expression variation in humans. Nat. Genet. 48, 22–29 (2016).

## References for Methods

59. Dobin, A. et al. STAR: ultrafast universal RNA-seq aligner. Bioinformatics 29, 15–21 (2013).

60. Neavin, D. et al. Demuxafy: improvement in droplet assignment by integrating multiple single-cell demultiplexing and doublet detection methods. Genome Biol. 25, 94 (2024).

61. Dolzhenko, E., et al. Resolving the unsolved: Comprehensive assessment of tandem repeats at scale. bioRxiv (2023) doi:10.1101/2023.05.12.540470.

62. Dolzhenko, E. et al. ExpansionHunter: a sequence-graph-based tool to analyze variation in short tandem repeat regions. Bioinformatics 35, 4754–4756 (2019).

63. Mousavi, N. et al. TRTools: a toolkit for genome-wide analysis of tandem repeats. Bioinformatics 37, 731–733 (2021).

64. Wolf, F. A., Angerer, P. & Theis, F. J. SCANPY: large-scale single-cell gene expression data analysis. Genome Biol. 19, 15 (2018).

65. Bais, A. S. & Kostka, D. scds: computational annotation of doublets in single-cell RNA sequencing data. Bioinformatics 36, 1150–1158 (2020).

66. Huang, Y., McCarthy, D. J. & Stegle, O. Vireo: Bayesian demultiplexing of pooled single-cell RNA-seq data without genotype reference. Genome Biol. 20, 273 (2019).

67. Germain, P.-L., Lun, A., Garcia Meixide, C., Macnair, W. & Robinson, M. D. Doublet identification in single-cell sequencing data using. F1000Res. 10, 979 (2021).

68. Michielsen, L., Reinders, M. J. T. & Mahfouz, A. Hierarchical progressive learning of cell identities in single-cell data. Nat. Commun. 12, 2799 (2021).

69. Hao, Y. et al. Integrated analysis of multimodal single-cell data. Cell 184, 3573–3587.e29 (2021).

70. Xu, C. et al. Automatic cell-type harmonization and integration across Human Cell Atlas datasets. Cell 186, 5876–5891.e20 (2023).

71. Domínguez Conde, C., et al. Cross-tissue immune cell analysis reveals tissue-specific features in humans. Science 376, eabl5197 (2022).

72. Cuomo, A. S. E. et al. Optimizing expression quantitative trait locus mapping workflows for single-cell studies. Genome Biol. 22, 188 (2021).

73. Balduzzi, S., Rücker, G. & Schwarzer, G. How to perform a meta-analysis with R: a practical tutorial. Evid. Based Ment. Health 22, 153–160 (2019).

74. Liu, Y. et al. ACAT: A fast and powerful p value combination method for rare-variant analysis in sequencing studies. Am. J. Hum. Genet. 104, 410–421 (2019).

75. Storey, J. D. The positive false discovery rate: a Bayesian interpretation and the q-value. Ann. Stat. 31, 2013–2035 (2003).

76. Franco, L. M. et al. Immune regulation by glucocorticoids can be linked to cell type-dependent transcriptional responses. J. Exp. Med. 216, 384–406 (2019).

77. Zou, Y., Carbonetto, P., Wang, G. & Stephens, M. Fine-mapping from summary data with the ‘Sum of Single Effects’ model. PLoS Genet. 18, e1010299 (2022).

78. Nasser, J. et al. Genome-wide enhancer maps link risk variants to disease genes. Nature 593, 238–243 (2021).

79. Ernst, J. & Kellis, M. Chromatin-state discovery and genome annotation with ChromHMM. Nat. Protoc. 12, 2478–2492 (2017).

80. Casper, J. et al. The UCSC Genome Browser database: 2026 update. Nucleic Acids Res. 54, D1331–D1335 (2026).

81. Tsuo, K. et al. Multi-ancestry meta-analysis of asthma identifies novel associations and highlights the value of increased power and diversity. Cell Genom. 2, 100212 (2022).

82. Liu, Z. et al. Genetic architecture of the inflammatory bowel diseases across East Asian and European ancestries. Nat. Genet. 55, 796–806 (2023).

83. Kiryluk, K. et al. Genome-wide association analyses define pathogenic signaling pathways and prioritize drug targets for IgA nephropathy. Nat. Genet. 55, 1091–1105 (2023).

84. Ishigaki, K. et al. Multi-ancestry genome-wide association analyses identify novel genetic mechanisms in rheumatoid arthritis. Nat. Genet. 54, 1640–1651 (2022).

85. Bentham, J. et al. Genetic association analyses implicate aberrant regulation of innate and adaptive immunity genes in the pathogenesis of systemic lupus erythematosus. Nat. Genet. 47, 1457–1464 (2015).

86. Chiou, J. et al. Interpreting type 1 diabetes risk with genetics and single-cell epigenomics. Nature 594, 398–402 (2021).

87. Michailidou, K. et al. Association analysis identifies 65 new breast cancer risk loci. Nature 551, 92–94 (2017).

88. McKay, J. D. et al. Large-scale association analysis identifies new lung cancer susceptibility loci and heterogeneity in genetic susceptibility across histological subtypes. Nat. Genet. 49, 1126–1132 (2017).

89. Sakaue, S. et al. A cross-population atlas of genetic associations for 220 human phenotypes. Nat. Genet. 53, 1415–1424 (2021).

90. Wang, A. et al. Characterizing prostate cancer risk through multi-ancestry genome-wide discovery of 187 novel risk variants. Nat. Genet. 55, 2065–2074 (2023).

91. COVID-19 Host Genetics Initiative. The COVID-19 Host Genetics Initiative, a global initiative to elucidate the role of host genetic factors in susceptibility and severity of the SARS-CoV-2 virus pandemic. Eur. J. Hum. Genet. 28, 715–718 (2020).

92. Bellenguez, C. et al. New insights into the genetic etiology of Alzheimer’s disease and related dementias. Nat. Genet. 54, 412–436 (2022).

93. International Multiple Sclerosis Genetics Consortium (IMSGC) et al. Analysis of immune-related loci identifies 48 new susceptibility variants for multiple sclerosis. Nat. Genet. 45, 1353–1360 (2013).

94. Nalls, M. A. et al. Identification of novel risk loci, causal insights, and heritable risk for Parkinson’s disease: a meta-analysis of genome-wide association studies. Lancet Neurol. 18, 1091–1102 (2019).

