## Supplementary Information Guide for "Tandem repeat variation shapes immune cell type-specific gene expression"

**Table of Contents**

**Supplementary Methods** ……………………………………………………………..………………1

**Benchmarking against Simons Genome Diversity Project (SGDP) samples**
**PacBio HiFi sequencing, alignment, and TR genotyping**

**Supplementary Notes**………………………………………………………………..………………3

**Supplementary Note 1**: Benchmarking ExpansionHunter performance with capillary electrophoresis and long read genotypes
**Supplementary Note 2:** Median pairwise sharing across all cell types
**Supplementary Note 3**: Cell type-specific poly(GCA) repeat is associated with expression of *GLS* and lnc-NEMP2-1
**Supplementary Note 4**: Benchmarking of cell-state interaction models

**Supplementary Figures**…………………………………………………………………………………5

**Supplementary Figure 1**: Principal component analysis (PCA) of TenK10K Phase 1 cohort samples overlaid with global reference populations
**Supplementary Figure 2**: Accuracy of ExpansionHunter, HipSTR, and GangSTR as a function of sensitivity
**Supplementary Figure 3**: Concordance between long-read TRGT calls and short-read ExpansionHunter calls
**Supplementary Figure 4**: Distribution of mean insert size (bp) of samples prepared with the KAPA Hyper PCR-free kit (Roche) and those prepared with the DNA PCR-free kit (Illumina)
**Supplementary Figure 5**: Distribution of percent of target bases with at least 30x coverage in samples prepared with the KAPA Hyper PCR-free kit (Roche) and those prepared with the DNA PCR-free kit (Illumina)
**Supplementary Figure 6** Distribution of median coverage (X) of samples prepared with the KAPA Hyper PCR-free kit (Roche) and those prepared with the DNA PCR-free kit (Illumina)
**Supplementary Figure 7**: Distribution of reads mapped in the BAM file (millions) of samples prepared with the KAPA Hyper PCR-free kit (Roche) and those prepared with the DNA PCR-free kit (Illumina)
**Supplementary Figure 8**: Principal component analysis (PCA) plot of TR genotypes. Points are colored by sequencing library preparation kit
**Supplementary Figure 9**: Quality control plots of scRNA-seq data
**Supplementary Figure 10**: Power analysis
**Supplementary Figure 11**: Heatmap of pairwise correlations of *cis* sc-eSNV (FDR < 5%) posterior effect sizes across cell types, adjusted for global effect size sharing and power differences with multivariate adaptive shrinkage
**Supplementary Figure 12**: Proportion of sc-eTRs (n = 112,100 eTRs) and common sc-eSNVs (from Cuomo *et al.;* n = 154,932 eSNVs) intersecting each ENCODE candidate cis-regulatory element (cCRE) class
**Supplementary Figure 13**: Fold enrichment of tandem repeat motif lengths among sc-eTRs (n = 112,100 sc-eTRs) across genic regions
**Supplementary Figure 14**: Scatterplot of starting dosage (summed repeats) and effect size of sc-eTRs (n = 112,100 sc-eTRs)
**Supplementary Figure 15**: $Log_2$(fold enrichment) of cell state-dependent sc-eTRs (FDR < 5%; n = 579 sc-eTRs) relative to all TRs genotyped, by ENCODE cCRE annotation
**Supplementary Figure 16**: $Log_2$(fold enrichment) of cell type-specific sc-eTRs relative to shared sc-eTRs, by MANE and ENCODE cCRE annotation (n = 112,100 sc-eTRs)

**Supplementary Figure 17**: Quantile-quantile (QQ) plot comparing observed two-sided *P* values for each TR x ATAC peak test (colored by cell type) against the expected uniform distribution
**Supplementary Figure 18:** Dot plot showing the enrichment ($\log_2$ odds ratio) of caQTL$^+$ sc-eTRs relative to caQTL- sc-eTRs (n= 112,100 sc-eTRs) across detailed ENCODE cCRE annotations
**Supplementary Figure 19**: Linkage disequilibrium (LD, calculated as $R^2$) of lead variant with next most significant neighboring SNV (within +/- 100 kb of the gene body) in the $CD4_{TCM}$ cell type, grouped by loci where the lead variant is a SNV or a TR (n = 13,093 genes)
**Supplementary Figure 20**: Distribution of the ratio of p-values of the lead variant relative to that of the next significant neighboring variant (within +/- 100 kb of the gene body) in the $CD4_{TCM}$ cell type, grouped by loci where the lead variant is a SNV or a TR (n = 13,093 genes)
**Supplementary Figure 21**: Enrichment of sc-eTRs and candidate causal sc-eTRs relative to all TRs genotyped
**Supplementary Figure 22**: Enrichment of candidate causal sc-eTRs relative to all TRs genotyped in primate-specific and Zoonomia annotations
**Supplementary Figure 23**: Number of eGenes (associated with fine-mapped sc-eTRs [PIP ≥ 0.7]) by loss-of-function observed/expected upper bound fraction (LOEUF) score decile
**Supplementary Figure 24**: Number of eGenes (associated with sc-eSNVs [FDR <5%]) by loss-of-function observed/expected upper bound fraction (LOEUF) score decil**e**
**Supplementary Figure 25**: Distribution of linkage disequilibrium (LD) with the lead SNV of the respective *cis* window for all sc-eTRs and for candidate causal sc-eTRs
**Supplementary Figure 26**: Cell type-specific associations of poly (GCA) repeat and expression of *GLS* and lnc-NEMP2-1
**Supplementary Figure 27**: Barplot of the number of candidate causal sc-eTRs intersecting significant TR UKBB PheWAS hits, colored by gene annotation
**Supplementary Figure 28**: Barplot of the proportion of colocalized genes (PPH4 ≥0.8) containing a candidate causal sc-eTR, per GWAS trait tested
**Supplementary Figure 29:** Principal components (PCs) for BioHEART cohort (n=975 individuals)
**Supplementary Figure 30:** Principal components (PCs) for TOB cohort (n=950 individuals)
**Supplementary Figure 31**: Changes in the number of *cis*-eGenes detected with the addition of expression principal components
**Supplementary Figure 32**: Variance ratio plots for the TOB cohort across 28 immune cell types
**Supplementary Figure 33**: Variance ratio plots for the BioHEART cohort across 28 immune cell types
**Supplementary Figure 34**: sc-eTR discovery in $CD4_{TCM}$ using +/- 1Mb *cis* window around the gene body
**Supplementary Figure 35**: Distribution of methylation probability (per CpG site) across chromosomes and n=25 samples
**Supplementary Figure 36**: Principal component analysis of methylation probability values across 26,191,720 CpG sites in chr1-22

**Supplementary Tables**……………………………………………………………………….34

*All supplementary tables, except for Supplementary Table 8, are provided as supplementary files.*

**Supplementary Table 1**: List of single-cell expression quantitative trait TR loci (sc-eTRs) (FDR < 5%)
**Supplementary Table 2**: Cell state sc-eTRs identified in B cells (FDR < 5%)
**Supplementary Table 3**: List of single-cell eTRs (nominal P-value < 0.05) that overlapped with bulk eTRs previously identified in Fotsing et al. (DOI: 10.1038/s41588-019-0521-9) and Bakhtiari et al. (DOI: 10.1038/s41467-021-22206-z)
**Supplementary Table 4**: List of single-cell expression quantitative trait TR loci (sc-eTRs) (FDR < 5%) that associate with a nearby sc-ATAC peak (within 10kb of the TR) (FDR < 5%)
**Supplementary Table 5**: HOMER results of transcription factor binding sites enriched in sc-eTRs that associate with scATAC-seq (caQTL+ sc-eTRs) (FDR < 5%)

**Supplementary Table 6**: List of candidate causal sc-eTRs
**Supplementary Table 7**: List of candidate causal sc-eTRs post-conditioning on the lead SNV (FDR < 5%)
**Supplementary Table 8:** Candidate causal sc-eTRs intersecting known disease-associated loci
**Supplementary Table 9**: List of eGenes associated with candidate causal sc-eTRs intersecting with list of genes curated in PanelApp Australia (downloaded on October 15, 2024)
**Supplementary Table 10**: List of candidate causal sc-eTRs intersecting UKBB TR PheWAS hits (Manigbas et al. 2024 [DOI: 10.1038/s41467-024-54678-0])
**Supplementary Table 11:** List of colocalized GWAS loci [PP H4 ≥ 0.8] containing at least one sc-eTR that is candidate causal for gene expression
