## Supplementary Information for "Tandem repeat variation shapes immune cell type-specific gene expression"

Supplementary Methods

***Benchmarking against Simons Genome Diversity Project (SGDP) samples***

The performance of ExpansionHunter (v5)[1], GangSTR (v2.5)[2], and HipSTR (v0.6.2)[3] were benchmarked using capillary electrophoresis results of hypervariable microsatellite loci. Specifically, WGS data and capillary electrophoresis results of 120 samples from the Simons Genome Diversity Project (SGDP) were downloaded from publicly accessible websites[4]. FASTQ files were downloaded from ENA Archive (accession no. ERP010710) and aligned to hg38 reference genome with DRAGMAP (https://github.com/Illumina/DRAGMAP), according to National Approach to Genomic Information Management (NAGIM) guidelines (https://queenslandgenomics.org/qldgenomics-updated/wp-content/uploads/2021/02/NAGIM-Blueprint-v20201010-Final-v1.2.2.pdf). Capillary electrophoresis results (in bp) corresponding to 627 STR loci were downloaded from (https://web.stanford.edu/group/rosenberglab/data/pembertonEtAl2009/combinedmicrosats_627loci_1048indivs_numRpts.stru). Primer sequences and the repeat motif of each target PCR amplicon were downloaded from (https://web.stanford.edu/group/rosenberglab/data/pembertonEtAl2009/Pemberton_AdditionalFile1_11242009.txt). We filtered for motif structures that were simple (consisting of one motif) and pure (without any sequence interruptions), resulting in a subset of 482 loci. Primer sequences of the selected loci were then mapped to hg38 reference genome using *in silico* PCR[5]. 94 loci were removed from the dataset as primer sequences could not be mapped to hg38. This produced a PCR validation dataset of 388 STR loci.

To confirm the TR motif, the number of repeat copies found in the reference genome, and the coordinates of each TR locus, TandemRepeatFinder[6] was run on each target PCR amplicon using the settings match = 2, mismatch = 30, delta = 30, PM = 80, PI = 10, minscore = 16, maxperiod = 2000. TR loci initially defined on the antisense strand were further processed by taking the reverse complement of the sequence and re-performing TandemRepeatFinder to standardize all TR locus definitions by the sense strand.

To calculate the repeat copy allele from the PCR base pair result, the following formula was used at each TR locus: $c = r + (w - l)/s$, where *c* is the repeat copy, *r* is the number of repeats in hg38, *w* is the PCR amplified fragment (bp), *l* is the length of the PCR amplicon (bp) in hg38, and $s$ is the motif length (bp) of the target TR[7]. Repeat copy alleles were rounded down to the nearest whole number. PCR genotypes were masked for a sample at a particular locus if an indel was identified within the PCR target region but outside of the TR locus. Indels were identified using publicly accessible FermiKit calls (https://github.com/lh3/sgdp-fermi).

Using the coordinates of each TR locus, hg38 variant catalogs were created for ExpansionHunter, GangSTR, and HipSTR. All three callers were used to genotype 120 WGS samples. ExpansionHunter was run with --analysis-mode = streaming and --threads = 16 options. GangSTR was run using default parameters. HipSTR was run using the joint calling option.

In a subset of loci, non-zero offsets were applied to correct the PCR-derived repeat copy alleles. We followed a previously published protocol where, at each locus, every discordant call made by a TR caller relative to the PCR-derived genotype was recorded and the magnitude of the discordance was scored[8]. The offset was scored as a 1 if applying the offset to the genotype produced by the TR caller recovered both PCR-derived alleles. A score of 0.5 was given if the genotype produced by the TR caller was homozygous but applying the offset recovered only one PCR-derived allele. Lastly, a score of 0.25 was given if the genotype produced by the TR caller was heterozygous but the offset recovered only one PCR-derived allele. At each locus, offset calculations were calculated separately for each of the three TR callers. Only loci with at least 20 calls were considered. An offset was applied to the PCR-derived genotype only if it was the highest-ranking offset in calls made by all three TR callers at a particular locus.

***PacBio HiFi sequencing, alignment, and TR genotyping***

HiFi sequencing methods

Cells frozen in 90% FBS:10% DMSO were defrosted at room temperature. The cells were pelleted at 4°C, 500g for 5 minutes, washed with 100 µL of 1x PBS, and then pelleted again under the same conditions. The supernatant was discarded, and the pellet was processed immediately using the Nanobind CBB DNA extraction kit (102-301-900, PacBio). Cell counts ranged from 1.6 to 4.22 x $10^6$ cells, and extracted DNA was eluted in 150 µL of EB buffer.

DNA samples were homogenized using a Diagenode Megaruptor with the 3 DNAFluid+ Kit (E07020001, Diagenode) under the following conditions: volume 150 µL, speed 40, and concentration 50 ng/µL. Subsequently, 3 µg of material was diluted in low TE to a final volume of 130 µL. Shearing was performed with the Megaruptor shearing kit (E07010003, Diagenode) at speed 30 or 31, aiming for average fragment lengths of 15–24 kb. Clean-up and concentration of the sheared material were conducted using SMRTbell clean-up beads (102-158-300, PacBio), and the DNA was eluted in 47 µL. Average fragment lengths were determined using the Femto Pulse system with the Genomic DNA 165kb Analysis Kit (FP-1002-0275, Agilent).

SMRTbell libraries were prepared using the SMRTbell® Prep Kit 3.0 (102-141-700, PacBio) following standard procedures, including unique barcoding of each sample with the SMRTbell Adapter Index Plate 96A (102-009-200, PacBio). Size selection was performed with AMPure PB beads (102-182-500, PacBio) at a 2.9x ratio. Final library sizes were confirmed with the Femto Pulse, and libraries were diluted to below 60 ng/µL before the ABC loading procedure. SMRT library fragment lengths ranged between 12.895 and 26.001 kb. Sequencing was performed using the following PacBio products: Revio sequencing plate (02-587-400), Revio Polymerase kit (102-739-100), and Revio SMRT cell tray (102-202-200). On-plate loading concentration was set to 250 pM, with 30-hour movie times. SMRT libraries that did not yield 90 Gb using one SMRT cell were pooled for additional sequencing in "top-up" runs.

Alignment & TR genotyping

The resulting uBAM files were aligned to the hg38 reference genome using minimap2 (v2.28-r1209) with the following parameters: -y --secondary=no --MD -a -x map-hifi. Tandem repeats were genotyped in individual samples using TRGT (v1.1.0) with a custom catalog of polymorphic tandem repeats. The resulting tandem repeat genotypes were then merged into a single VCF for downstream analysis, also using TRGT.

### Supplementary Notes

**Supplementary Note 1: Benchmarking ExpansionHunter performance with capillary electrophoresis and long read genotypes**
Because TR genotyping is prone to stutter and remapping errors, we benchmarked the accuracy of three short-read TR genotyping tools - ExpansionHunter[1,9], HipSTR[3], and GangSTR[2] - by comparing their WGS-derived genotypes with gold standard capillary electrophoresis-derived genotypes of 388 STR loci in 120 samples from the Simons Genome Diversity Project[4] (**Supplementary Fig. 2**, **Supplementary Methods**). ExpansionHunter had the best performance with a validation rate of 95.0%, followed by HipSTR and GangSTR (94.9%, 87.4% respectively). The performance of ExpansionHunter and HipSTR are comparable to prior validation studies[3,9].

Using approximately 1.7 million filtered loci, we further validated the accuracy of ExpansionHunter with PacBio HiFi long-read TR genotypes (n=25 individuals), reporting a strict genotype concordance rate of 80.9%, which increased to 86.5% when allowing for +/- 1 repeat unit difference **(Fig. 1f, Methods)**. As expected, validation rate tended to decrease as TR reference length increased **(Fig. 1g)**. Notably, this PacBio genome-wide approach included TRs in low complexity regions, in contrast to the capillary electrophoresis validation dataset described above where TR loci were flanked by non-repetitive sequences and thus had improved read mappability (**Supplementary Fig. 3**), thus providing a more unbiased genome-wide estimate of error rates. While these error rates are (as expected) substantially higher than those seen for more simple variant classes such as SNVs, such errors are expected to reduce our sensitivity in eTR mapping, rather than to create false positive associations.

Comparing long-read and short-read TR genotypes, we found that poorly concordant calls were enriched in repeat loci with multiple adjacent motifs specified (odds ratio (OR) = 26.74; Fisher's two sided $P < 2.2\times10^{-16}$), reflecting the known limitation of short-read TR genotyping in low complexity regions with multiple adjacent, and possibly overlapping, repeat motifs[10] (**Supplementary Fig. 3**).

**Supplementary Note 2: Median pairwise sharing across all cell types**
Across all cell types, the median pairwise sharing of sc-eTRs was 57.7% (range: 5.1% to 98.7%). Notably, the B cell lineage showed the highest interlineage sharing (median = 93.0%). In T cells, however, sharing appears more closely aligned with functional state than with classical CD4/CD8 lineage markers: CD4 and CD8 central memory T (TCM) subtypes exhibited a median pairwise sharing of 65.6%, while effector memory (TEM) subtypes showed even higher sharing at 88.5%, compared to 41.8% and 49.0% for CD4 and CD8 lineages respectively. This likely reflects that memory T-cell subsets, irrespective of CD4/CD8 background, share highly similar transcriptional programs, leading to greater similarity in their underlying regulatory architecture.

**Supplementary Note 3**: **Cell type-specific poly(GCA) repeat is associated with expression of *GLS* and lnc-NEMP2-1**
While rare expansions of the poly(GCA) repeat in the 5' UTR of *GLS* is known to cause reduced *GLS* transcription[11], we found common variation of the repeat to be candidate causal for expression of two genes - *GLS* and lnc-NEMP2-1 (**Supplementary Fig. 26**, **Supplementary Table 8**). *GLS* encodes glutaminase which regulates glutamine, a key amino acid involved in immune proliferation and cytokine production[12]. Increasing TR length was found to associate with decreasing expression of *GLS* in $CD4_{TCM,}$ CD4 naive, $CD4_{TEM}$, NK, B naive, and $CD8_{TCM}$ (minimum $P = 3.63\times10^{-56}$). Meanwhile, increasing TR length was found to associate with increasing expression of lnc-NEMP2-1 in B naive, NK, CD4 naive, $CD4_{TCM}$, B memory, B intermediate, CD8 naive, $CD8_{TCM}$, NK CD56bright, MAIT (minimum $P = 8.27\times10^{-62}$).

**Supplementary Note 4: Benchmarking of cell-state interaction models**
We benchmarked several statistical frameworks to model cell state-dependent TR-gene interactions. In principle, a linear mixed-effects model with an interaction term between genotype and activity score would provide a unified analysis of repeated measurements per donor, as performed in the accompanying manuscript by Cuomo et *al*. However, in our pseudobulk setting each sample contributes up to three activity-stratified expression values, with the dataset including ~1,000 donors per cohort. Incorporating a random intercept for sample ID led to very slow model fitting and convergence issues when applied genome-wide. Using fixed effects for sample ID (i.e., adding ~1,000 indicator variables per model) slightly improved runtime but was not computationally feasible for tens of thousands of TR-gene pairs.

Given these constraints, we evaluated an alternative approach based on fitting separate pseudobulk eQTL models within each activity bin and comparing the resulting effect sizes using Wald z-tests. In benchmarking on a representative subset of loci, this z-test framework produced results directionally consistent with the mixed-effects interaction model while reducing computation time by several orders of magnitude. For this reason, we adopted the stratified effect size approach for the analysis.

Supplementary Figures

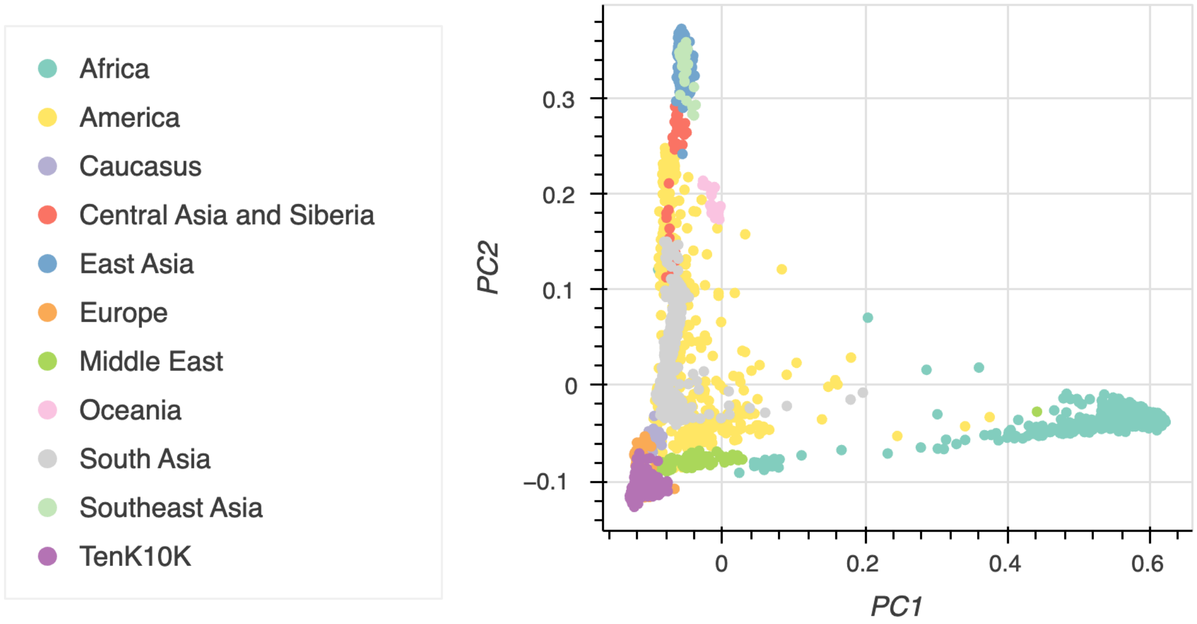


**Supplementary Figure 1: Principal component analysis (PCA) of TenK10K Phase 1 cohort samples overlaid with global reference populations.** Scatter plot of the first two principal components derived from SNVs, showing genetic variation among study participants (purple; n = 1,925 individuals) in the context of reference samples from the Human Genome Diversity Project (HGDP) and 1000 Genomes Project (1KGP). Reference samples are labeled by major continental ancestry groups. Study cohort samples cluster in genetic principal component space primarily with participants self-described as of European ancestry.

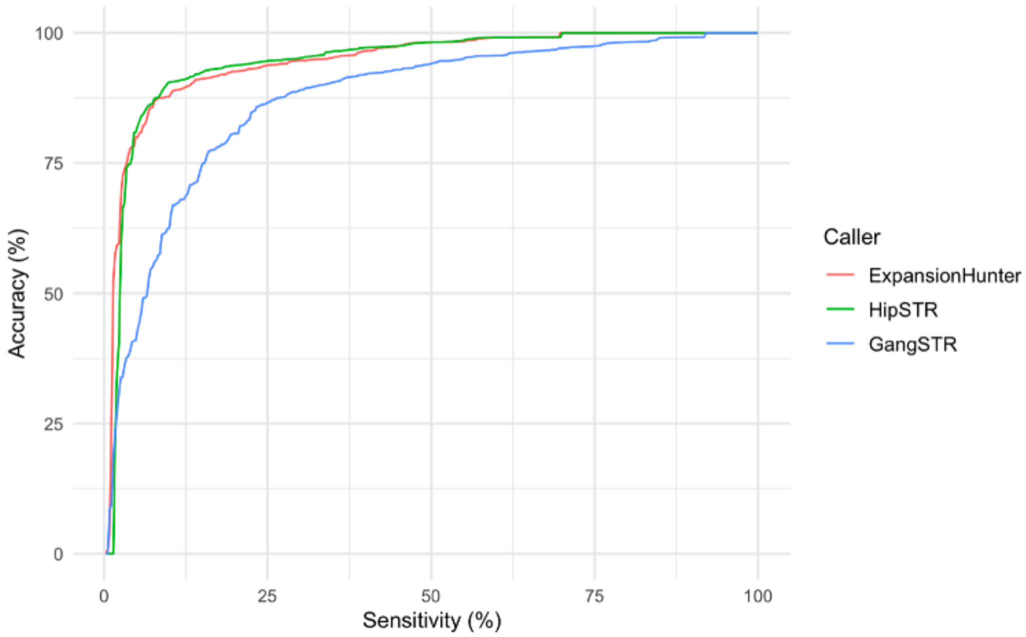


**Supplementary Figure 2: Accuracy of ExpansionHunter, HipSTR, and GangSTR as a function of sensitivity.** Accuracy was defined as the proportion of summed diploid genotypes that exactly match those in the PCR dataset (n = 388 STR loci), taking into account offsets (**Methods**). Sensitivity was defined as the proportion of STR loci in the catalog that was genotyped.

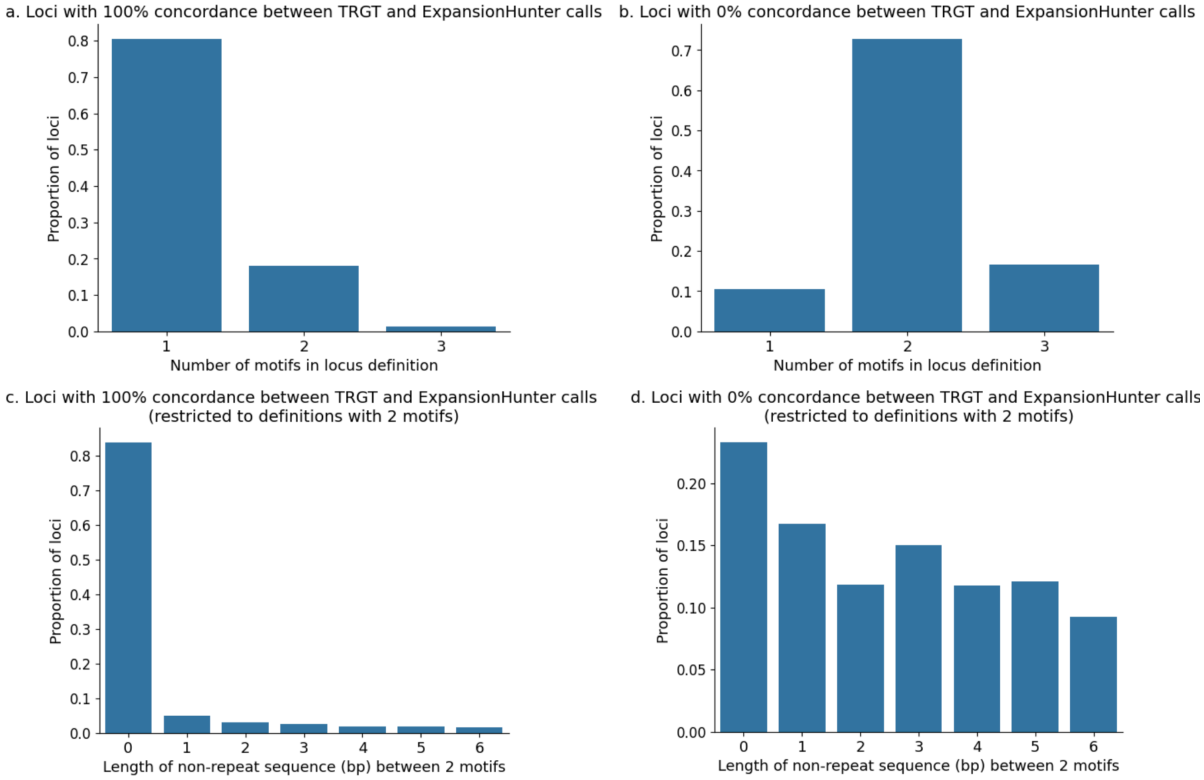


**Supplementary Figure 3**: **Concordance between long-read TRGT calls and short-read ExpansionHunter calls. a**, Distribution of compound motif definitions (n=25 samples) in loci with 100% concordance between long-read TRGT calls and short-read ExpansionHunter calls and **b**, those with 0% concordance. **c,** Distribution of interrupting (non-repeat) sequence length in loci defined by 2 motifs (n=25 samples) in loci with 100% concordance between long-read TRGT calls and short-read ExpansionHunter calls and **d**, 0% concordance.

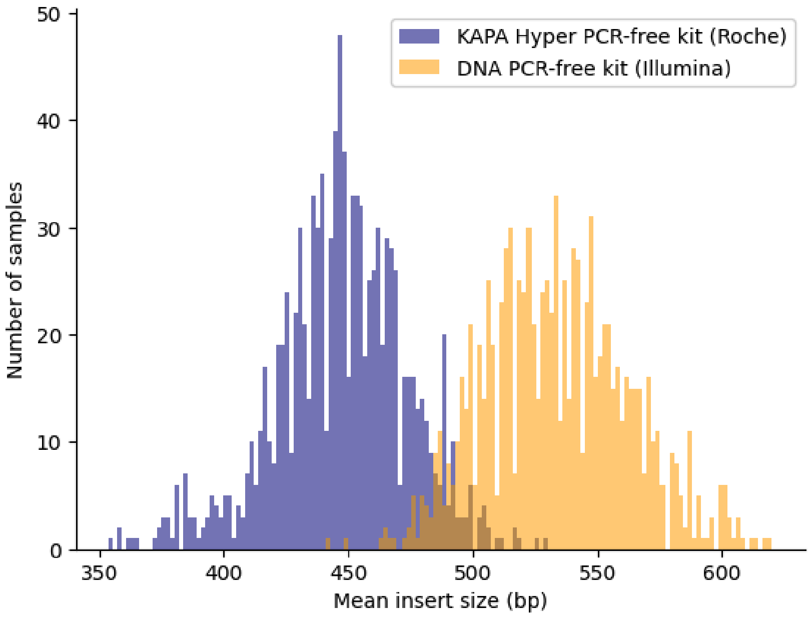


**Supplementary Figure 4: Distribution of mean insert size (bp) of samples prepared with the KAPA Hyper PCR-free kit (Roche) and those prepared with the DNA PCR-free kit (Illumina).**

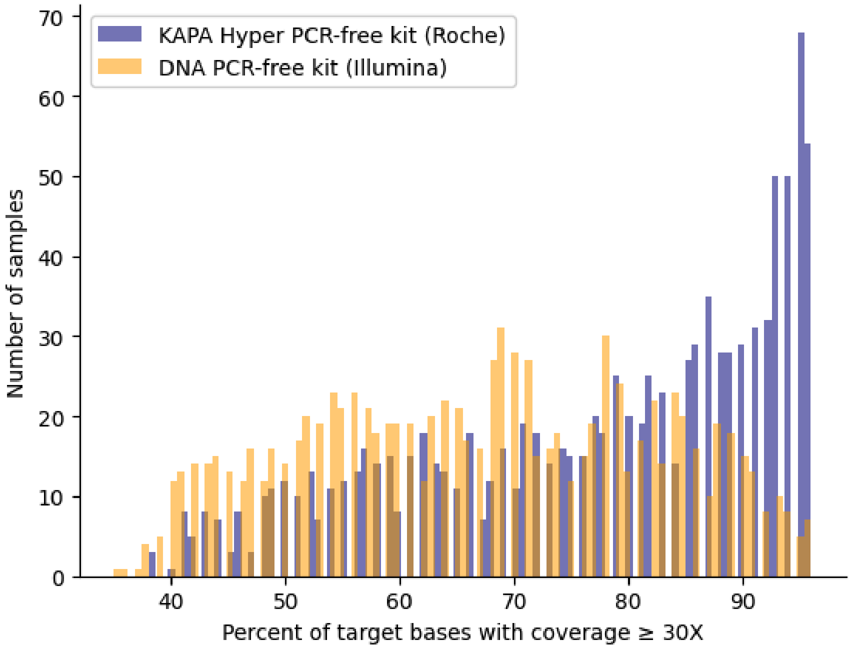


**Supplementary Figure 5: Distribution of percent of target bases with at least 30x coverage in samples prepared with the KAPA Hyper PCR-free kit (Roche) and those prepared with the DNA PCR-free kit (Illumina).**

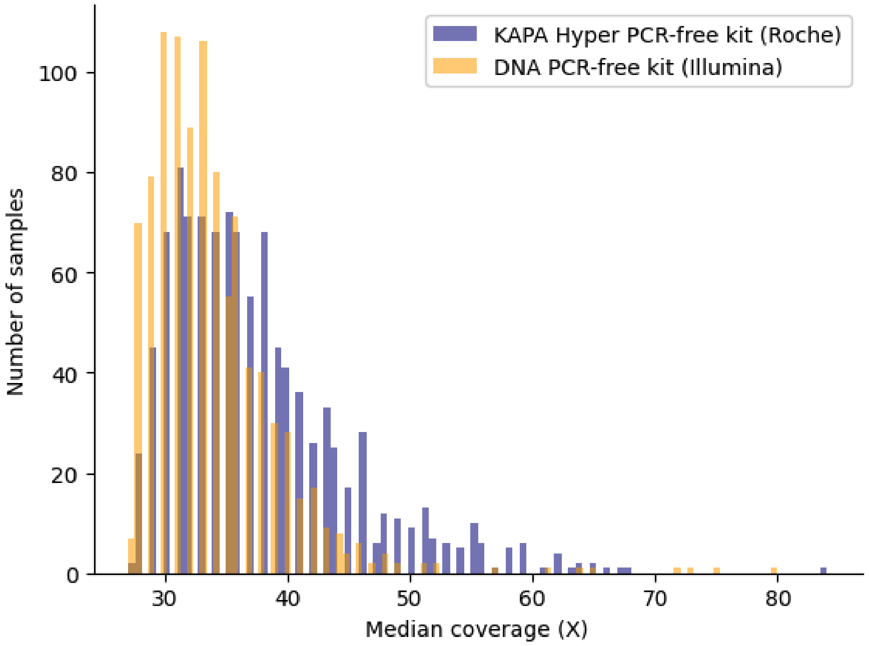


**Supplementary Figure 6: Distribution of median coverage (X) of samples prepared with the KAPA Hyper PCR-free kit (Roche) and those prepared with the DNA PCR-free kit (Illumina).**

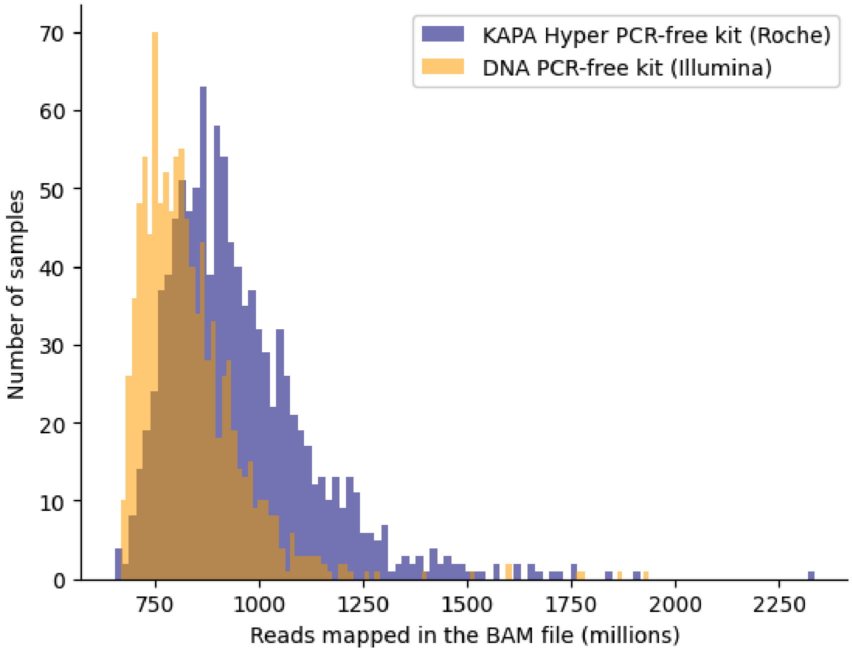


**Supplementary Figure 7: Distribution of reads mapped in the BAM file (millions) of samples prepared with the KAPA Hyper PCR-free kit (Roche) and those prepared with the DNA PCR-free kit (Illumina).**

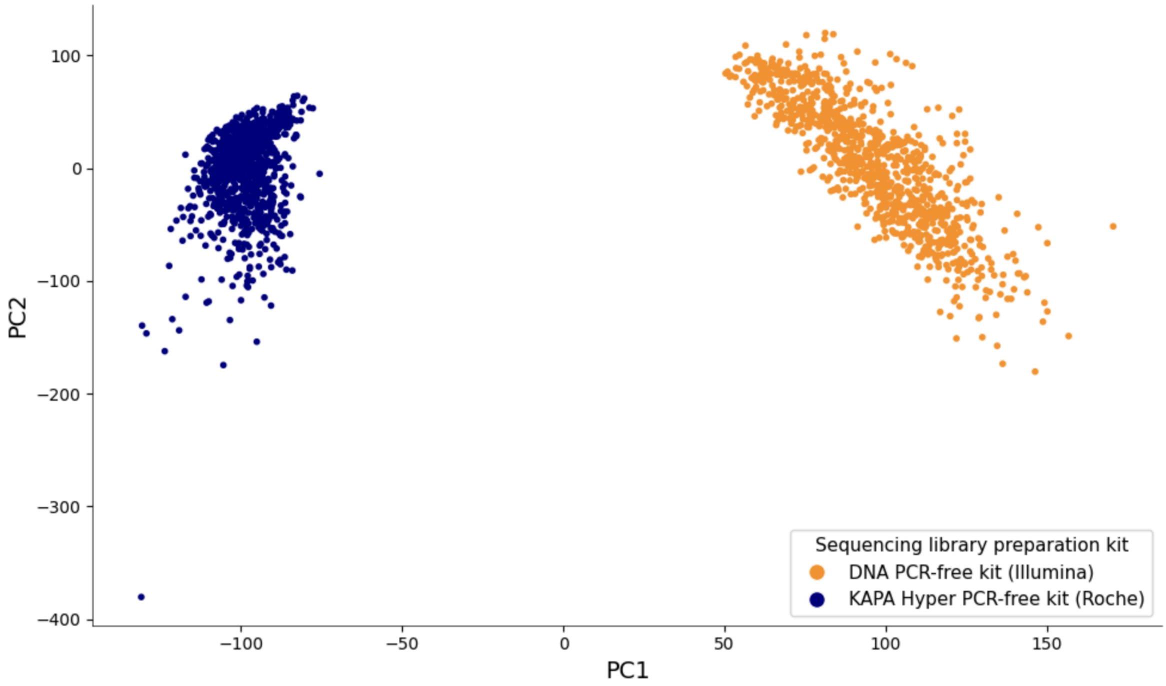


**Supplementary Figure 8: Principal component analysis (PCA) plot of TR genotypes. Points are colored by sequencing library preparation kit.** Sample sizes: Illumina DNA PCR-free kit (n=975 individuals) and Roche KAPA Hyper PCR-free kit (n=950 individuals).

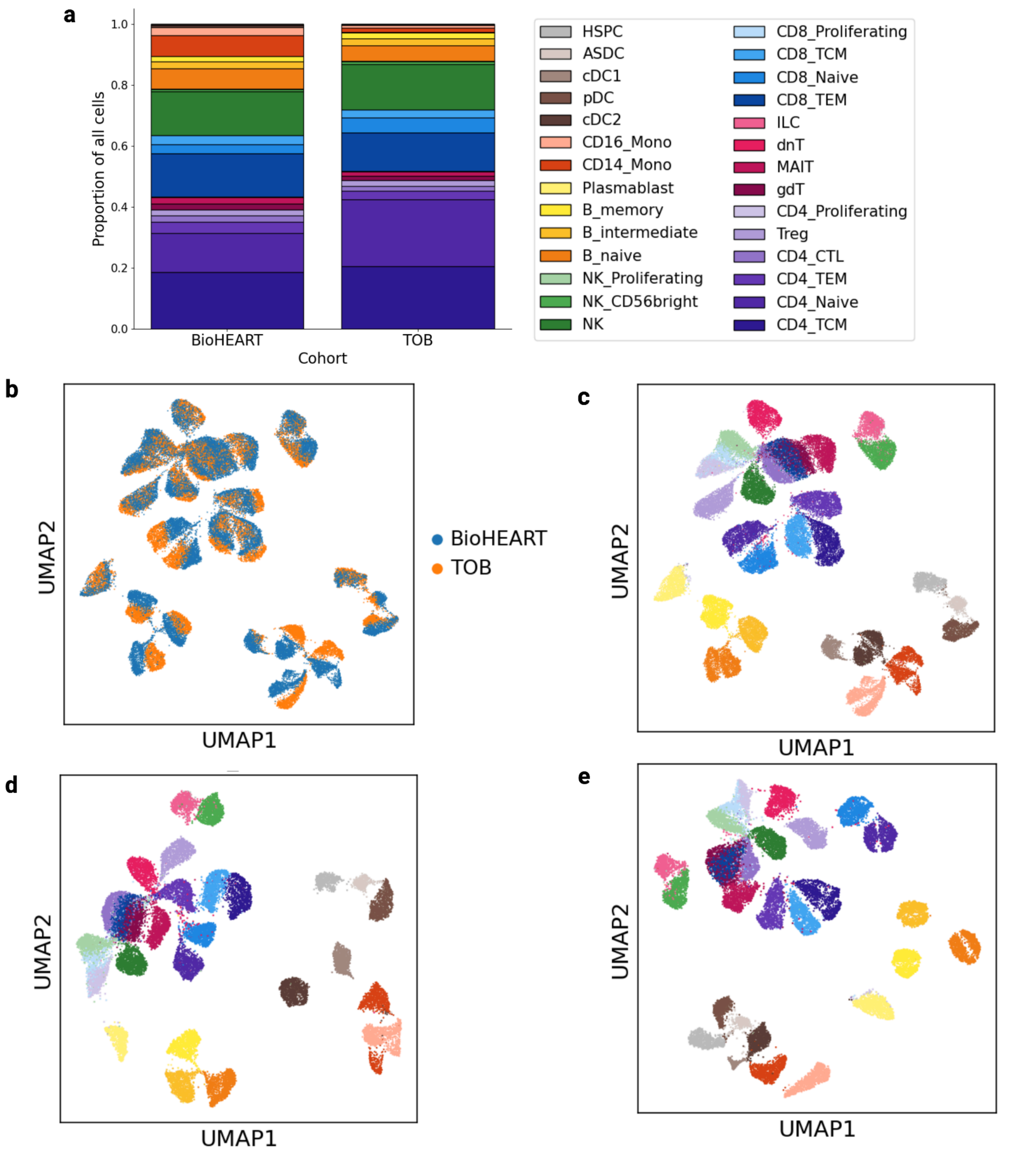


**Supplementary Figure 9: Quality control plots of scRNA-seq data. a,** Distribution of the proportion of cell types sequenced in each cohort. Legend is also applicable for panels **c**,**d**, and **e**. **b,** UMAP of pseudobulk scRNA-seq values, colored by cohort**. c,** UMAP of pseudobulk scRNA-seq values, colored by cell type. **d,** UMAP of pseudobulk scRNA-seq values in BioHEART cohort (n = 975 individuals), colored by cell type. **e,** UMAP of pseudobulk scRNA-seq values in TOB cohort (n = 950 individuals), colored by cell type. Sequencing library was regressed out of scRNA-seq data prior to pseudobulking in **b-e.**

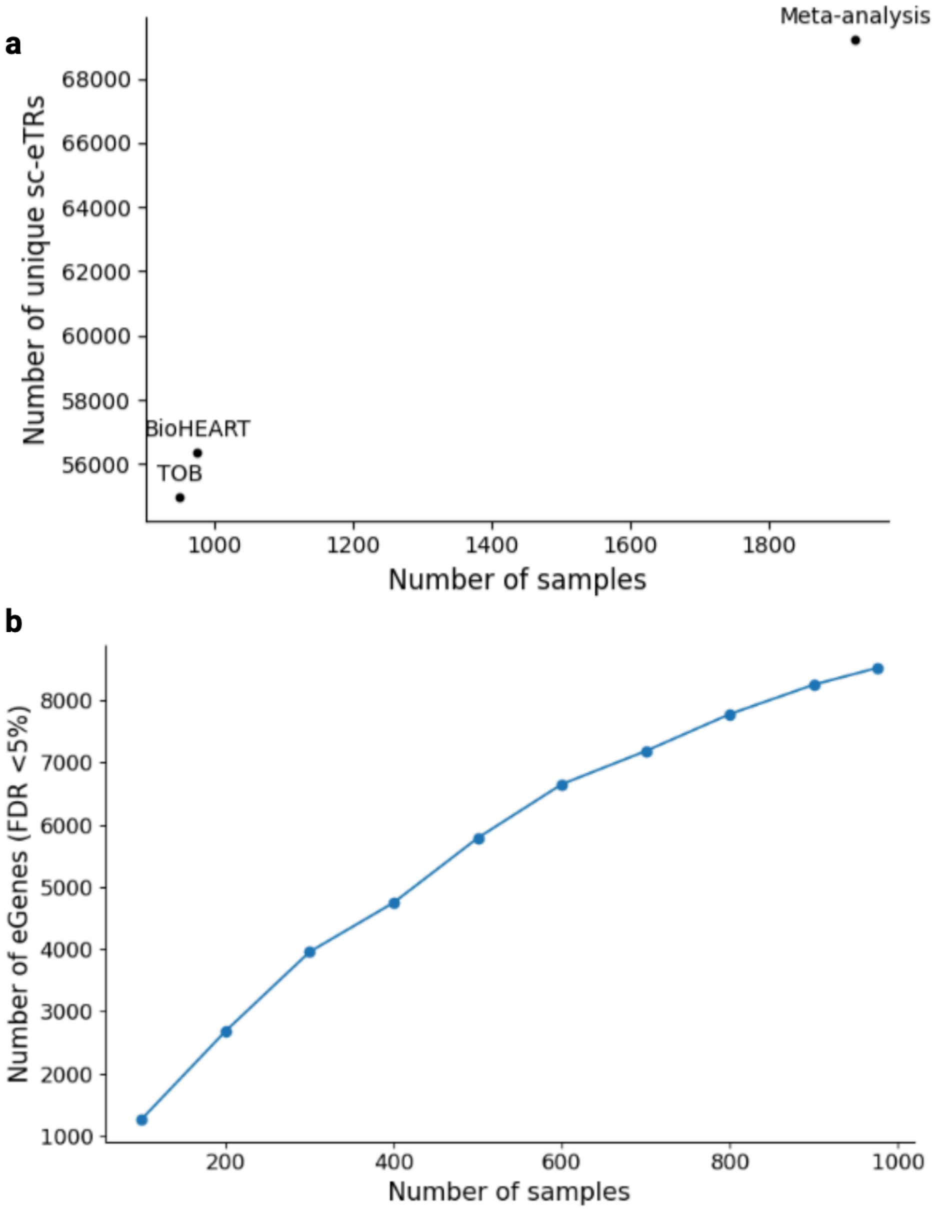


**Supplementary Figure 10: Power analysis a,** single-cell eTR meta-analysis discovers more unique sc-eTRs (FDR <5%) than individual cohorts (TOB [n=950 individuals]; BioHEART [n=975 individuals])**.** **b,** Line plot showing the number of eGenes discovered (FDR < 5%) as a function of sample size (n = 100 bins from 100 to 975 individuals, using samples from the BioHEART cohort).

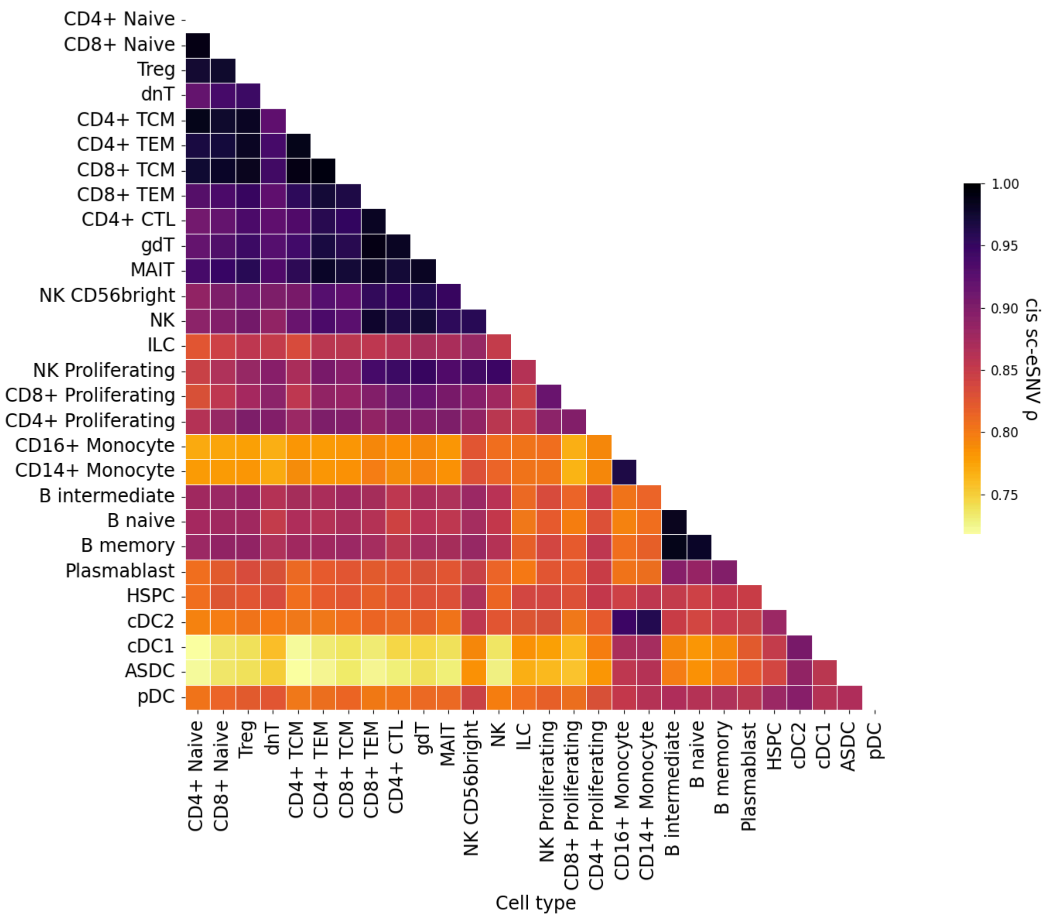


**Supplementary Figure 11: Heatmap of pairwise correlations of *cis* sc-eSNV (FDR < 5%) posterior effect sizes across cell types, adjusted for global effect size sharing and power differences with multivariate adaptive shrinkage**[16]. Order of rows and columns reflect scRNA-seq cell type classification by scPred[17] (**Methods**).

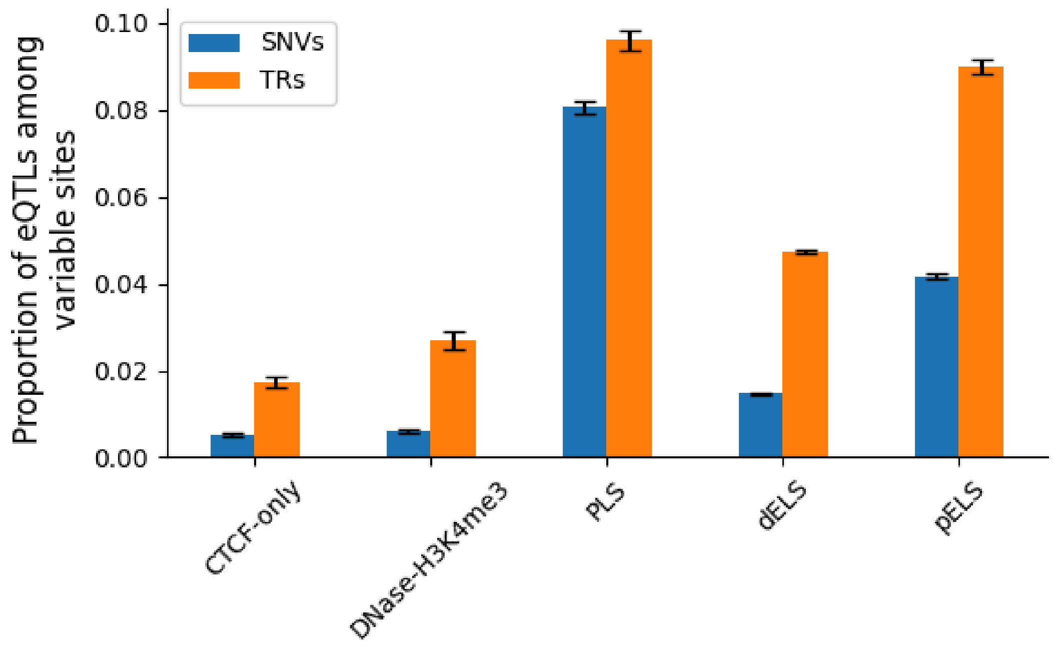


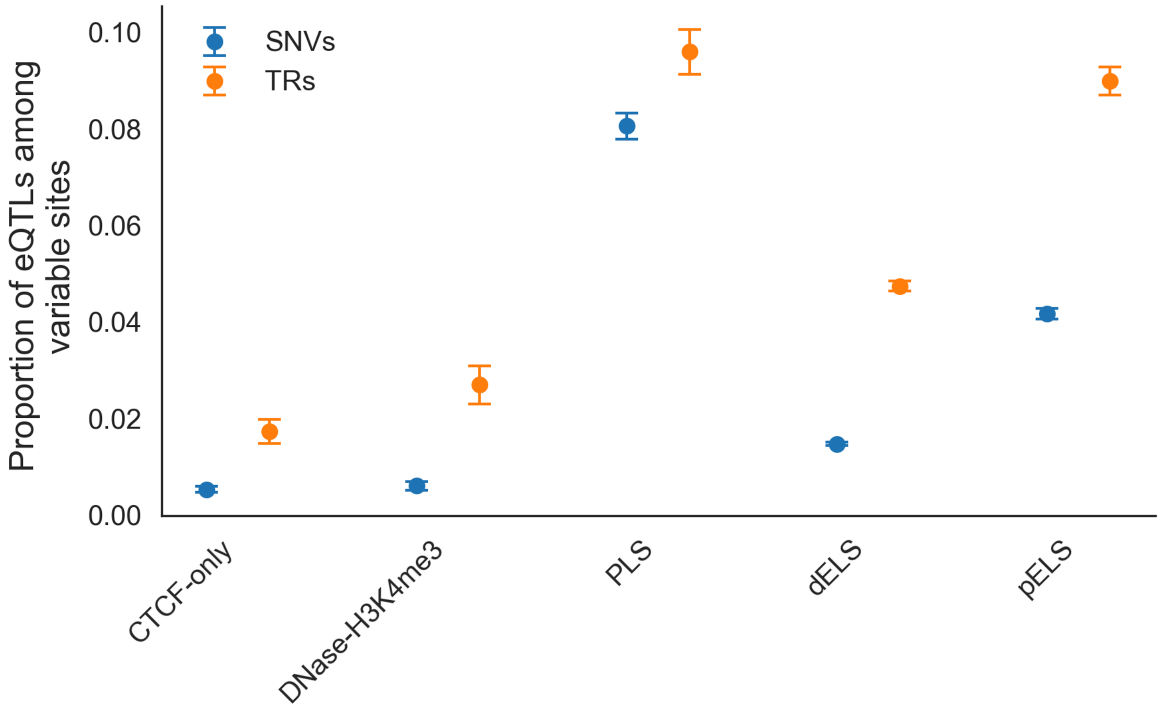


**Supplementary Figure 12: Proportion of sc-eTRs (n = 112,100 eTRs) and common sc-eSNVs (from Cuomo *et al.;* n = 154,932 eSNVs) intersecting each ENCODE candidate cis-regulatory element (cCRE) class.** Proportions (centre point) are normalised by the number of variable loci in each annotation (sc-eTRs ÷ variable TRs; sc-eSNVs ÷ common SNVs). Error bars correspond to 95% confidence intervals. ENCODE candidate *cis* regulatory element (cCRE) annotations are abbreviated as follows: promoter-like signature [PLS], proximal enhancer-like signature [pELS], distal enhancer-like signature [dELS], CTCF binding site failing to intersecting another cCRE annotation

[CTCF-only], DNase-H3K4me3 site failing to intersect promoter and enhancer signatures [DNase-H3K4me3].

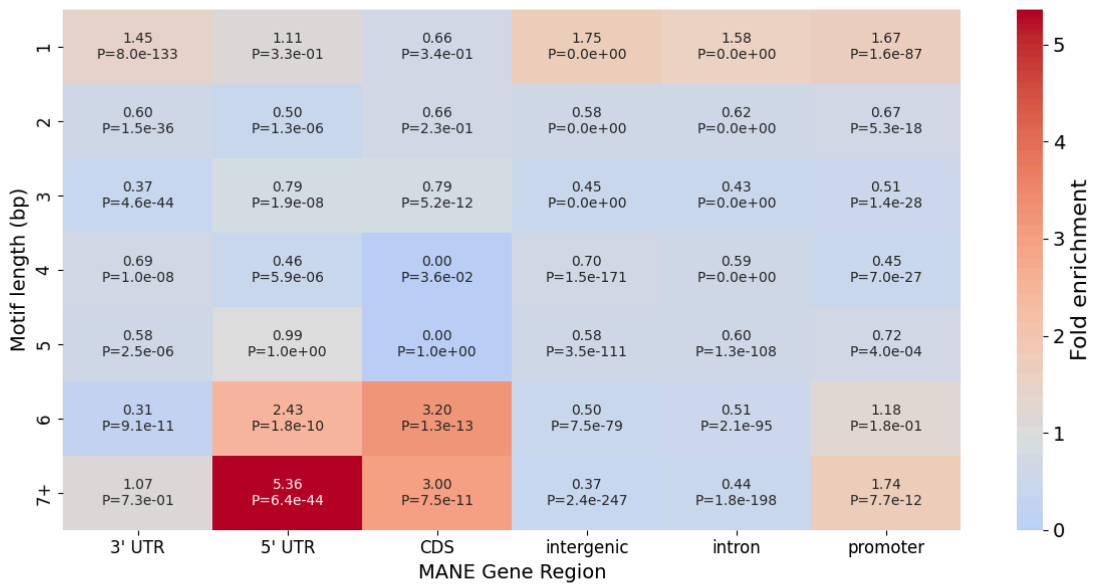


**Supplementary Figure 13: Fold enrichment of tandem repeat motif lengths among sc-eTRs (n = 112,100 sc-eTRs) across genic regions.** Heatmap showing the fold enrichment of TRs of varying motif lengths (1-7+ bp) among single-cell expression quantitative trait TRs (sc-eTRs) relative to the background polymorphic TR catalog, stratified by genomic region annotation (MANE). Each cell displays the fold enrichment and corresponding two-sided P-value from Fisher's exact test. CDS, coding sequence.

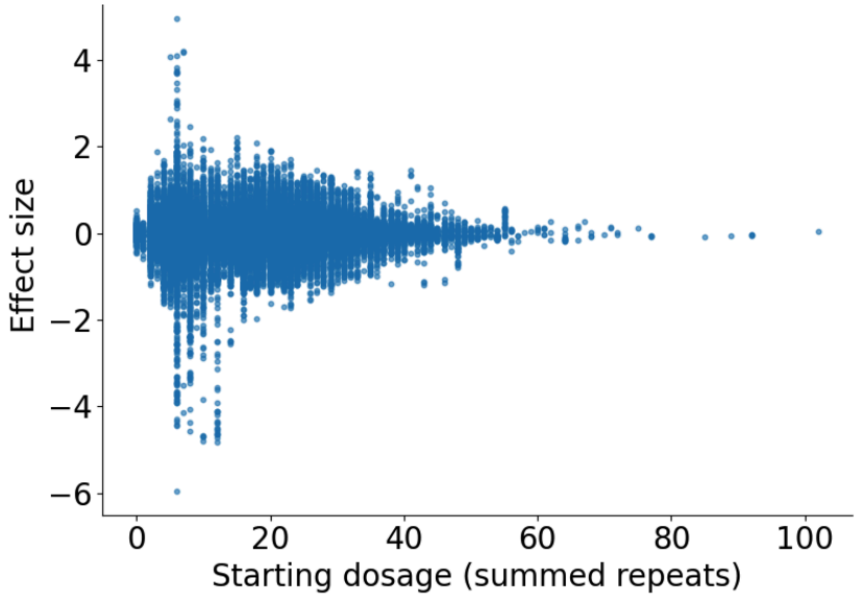


**Supplementary Figure 14: Scatterplot of starting dosage (summed repeats) and effect size of sc-eTRs (n = 112,100 sc-eTRs).** Starting dosage is the minimum dosage in the study cohort reported for a particular TR locus. The cluster of outlier effect sizes at starting dosages of ~5-15 repeats likely reflects the behaviour of short, highly polymorphic TRs, often dinucleotide or GC-rich motifs, where small length changes can cross regulatory thresholds (e.g., TF spacing or nucleosome positioning).

Repeats in this size range are also genotyped with high accuracy using short-read WGS, increasing power to detect strong sc-eTR effects.

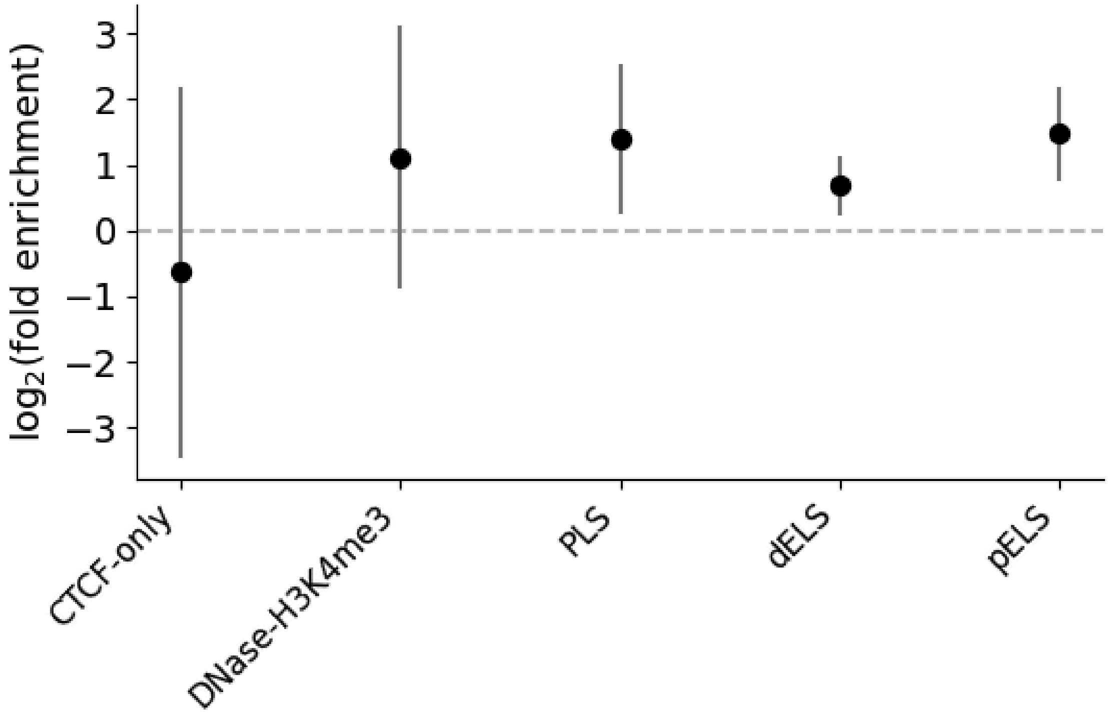


**Supplementary Figure 15: Log$_2$(fold enrichment) of cell state-dependent sc-eTRs (FDR < 5%; n = 579 sc-eTRs) relative to all TRs genotyped, by ENCODE cCRE annotation.** Error bars correspond to 95% confidence intervals. ENCODE candidate *cis* regulatory element (cCRE) annotations are abbreviated as follows: promoter-like signature [PLS], proximal enhancer-like signature [pELS], distal enhancer-like signature [dELS], and CTCF binding site failing to intersecting another cCRE annotation [CTCF-only].

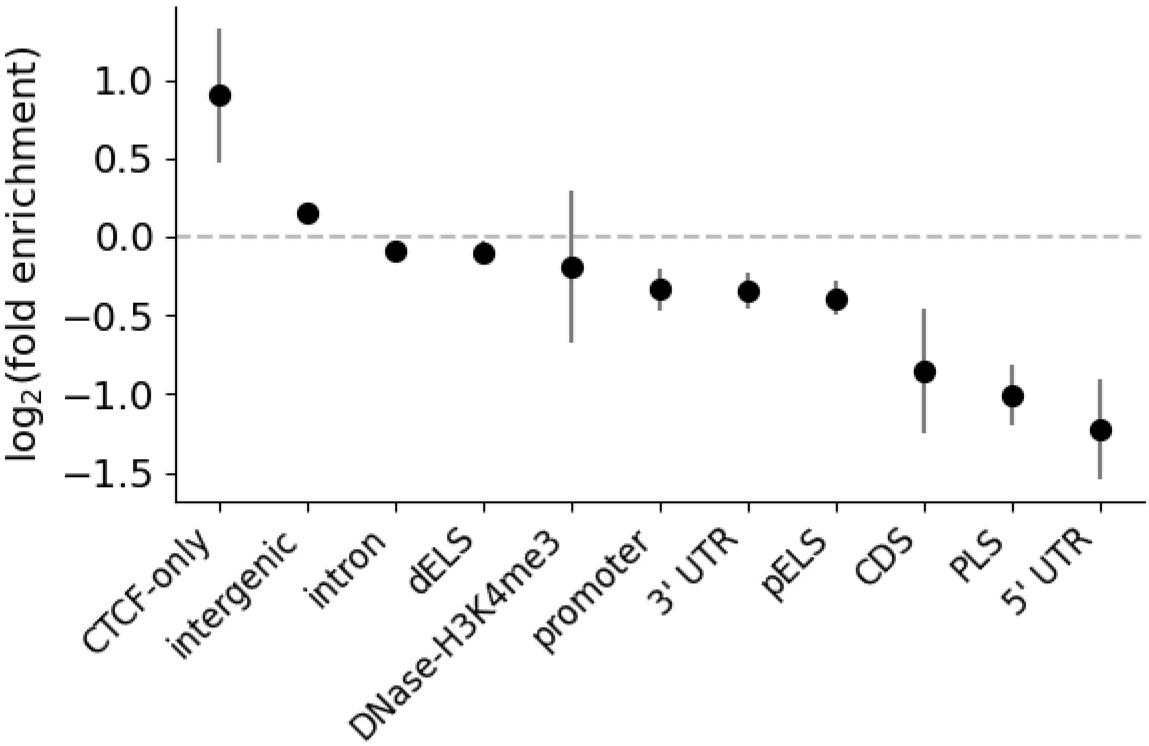


**Supplementary Figure 16**: **$Log_2$(fold enrichment) of cell type-specific sc-eTRs relative to shared sc-eTRs, by MANE and ENCODE cCRE annotation (n = 112,100 sc-eTRs)**. Error bars correspond to 95% confidence intervals. ENCODE candidate *cis* regulatory element (cCRE) annotations are abbreviated as follows: promoter-like signature [PLS], proximal enhancer-like signature [pELS], distal enhancer-like signature [dELS], CTCF binding site failing to intersecting another cCRE annotation [CTCF-only], DNase-H3K4me3 site failing to intersect promoter and enhancer signatures [DNase-H3K4me3]. MANE annotations are abbreviated as follows: 5' untranslated region [5' UTR], 3' untranslated region [3' UTR], coding sequence [CDS].

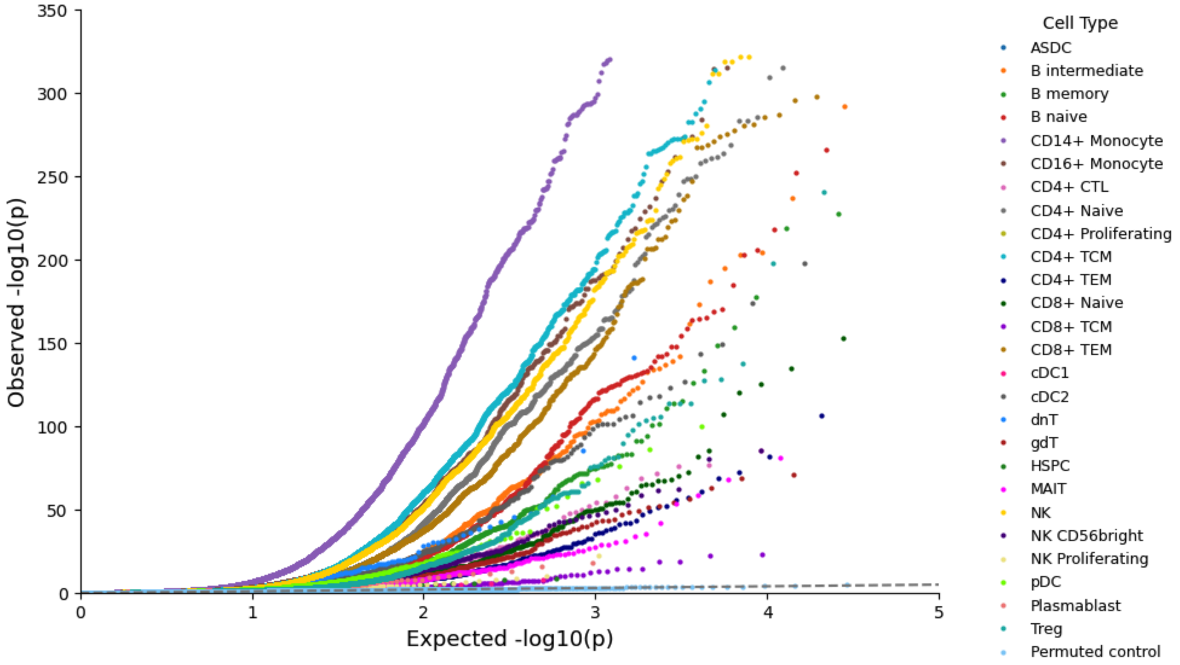


**Supplementary Figure 17: Quantile-quantile (QQ) plot comparing observed two-sided *P* values for each TR x ATAC peak test (colored by cell type) against the expected uniform distribution (dashed gray line).** Permuted control is highlighted in sky blue (n=922 individuals).

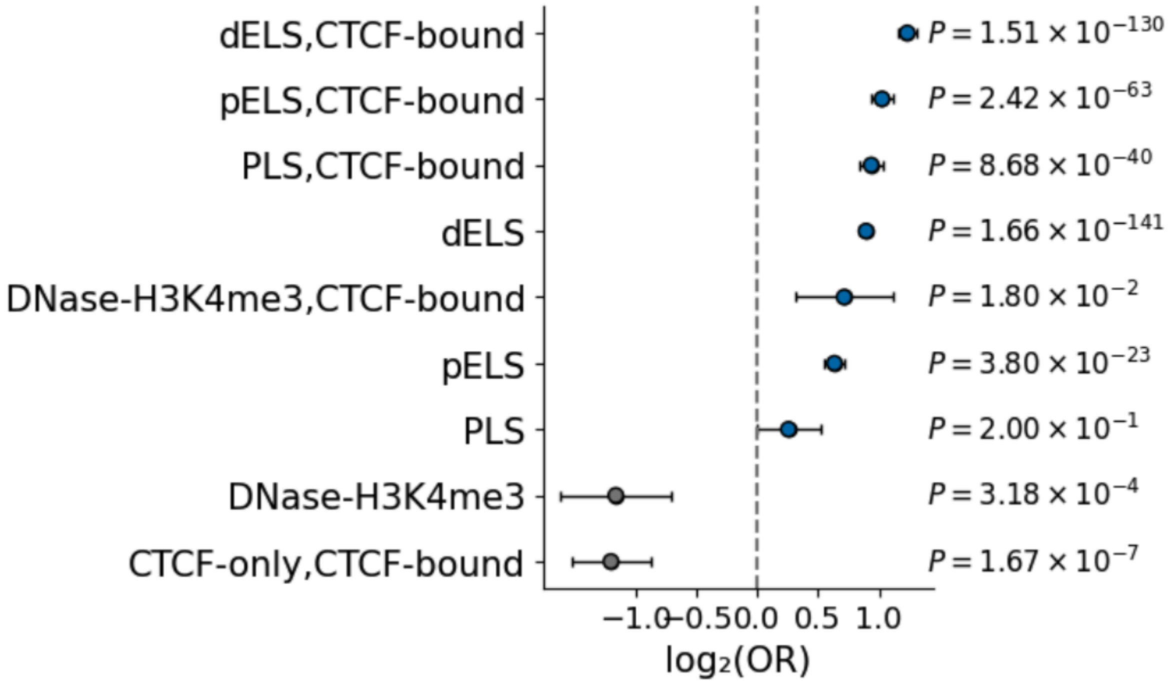


**Supplementary Figure 18: Dot plot showing the enrichment ($\log_2$ odds ratio) of caQTL$^+$ sc-eTRs relative to caQTL- sc-eTRs (n= 112,100 sc-eTRs) across detailed ENCODE cCRE annotations.** Error bars correspond to 95% confidence intervals. ENCODE candidate *cis* regulatory element (cCRE) annotations are abbreviated as follows: promoter-like signature [PLS], proximal enhancer-like signature [pELS], distal enhancer-like signature [dELS], CTCF binding site failing to intersecting another cCRE annotation [CTCF-only]. Two-sided Fisher's exact P values (uncorrected) are displayed.

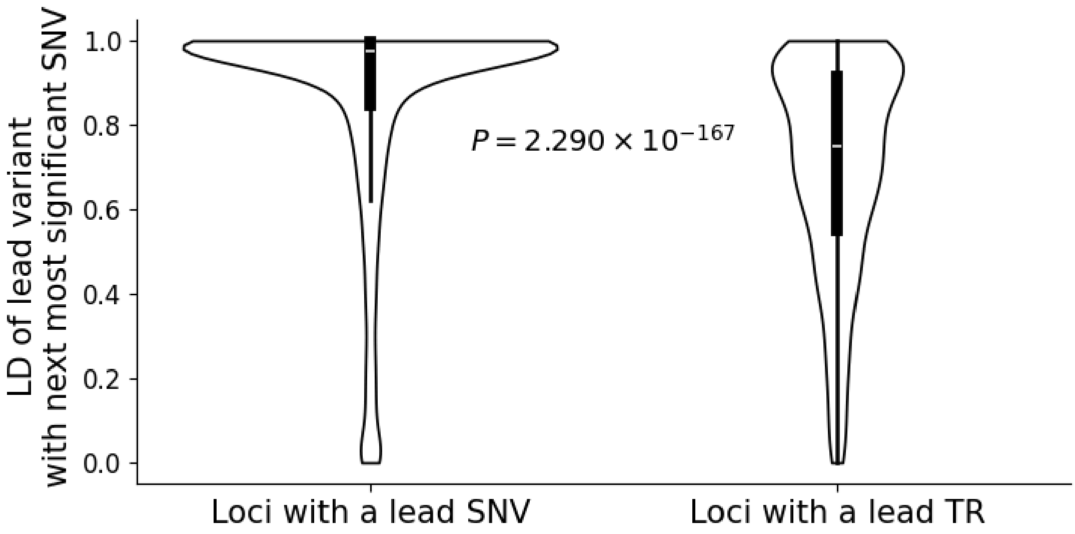


**Supplementary Figure 19: Linkage disequilibrium (LD, calculated as $R^2$) of lead variant with next most significant neighboring SNV (within +/- 100 kb of the gene body) in the $CD4_{TCM}$ cell type, grouped by loci where the lead variant is a SNV or a TR (n = 13,093 genes).** Violin plots summarize the distribution of effect sizes with horizontal lines showing median values and boxes spanning from the 25th (Q1) to the 75th percentile (Q3). Whiskers extend to Q1 – 1.5 x IQR and Q3 + 1.5 x IQR, where IQR is the interquartile range. Lead variants have nominal $P < 5\times10^{-8}$. P value derived from a two-sided Mann Whitney U test.

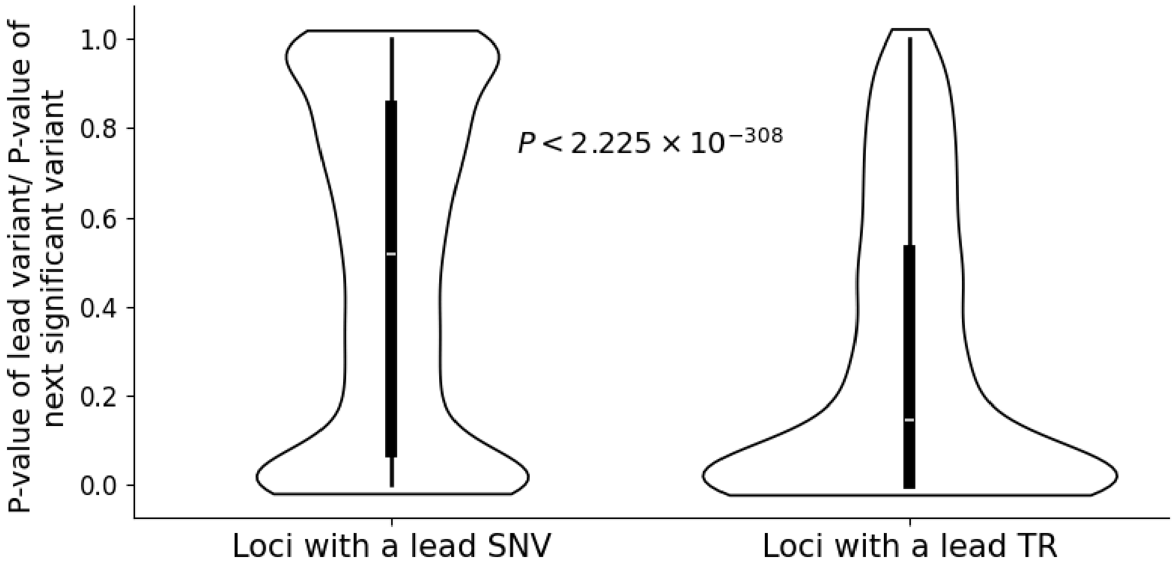


**Supplementary Figure 20: Distribution of the ratio of p-values of the lead variant relative to that of the next significant neighboring variant (within +/- 100 kb of the gene body) in the $CD4_{TCM}$ cell type, grouped by loci where the lead variant is a SNV or a TR (n = 13,093 genes).** Violin plots summarize the distribution of effect sizes with horizontal lines showing median values and boxes spanning from the 25th (Q1) to the 75th percentile (Q3). Whiskers extend to Q1 – 1.5 x IQR and Q3 + 1.5 x IQR, where IQR is the interquartile range. Lead variants have nominal $P < 5\times10^{-8}$. P value derived from a two-sided Mann Whitney U test.

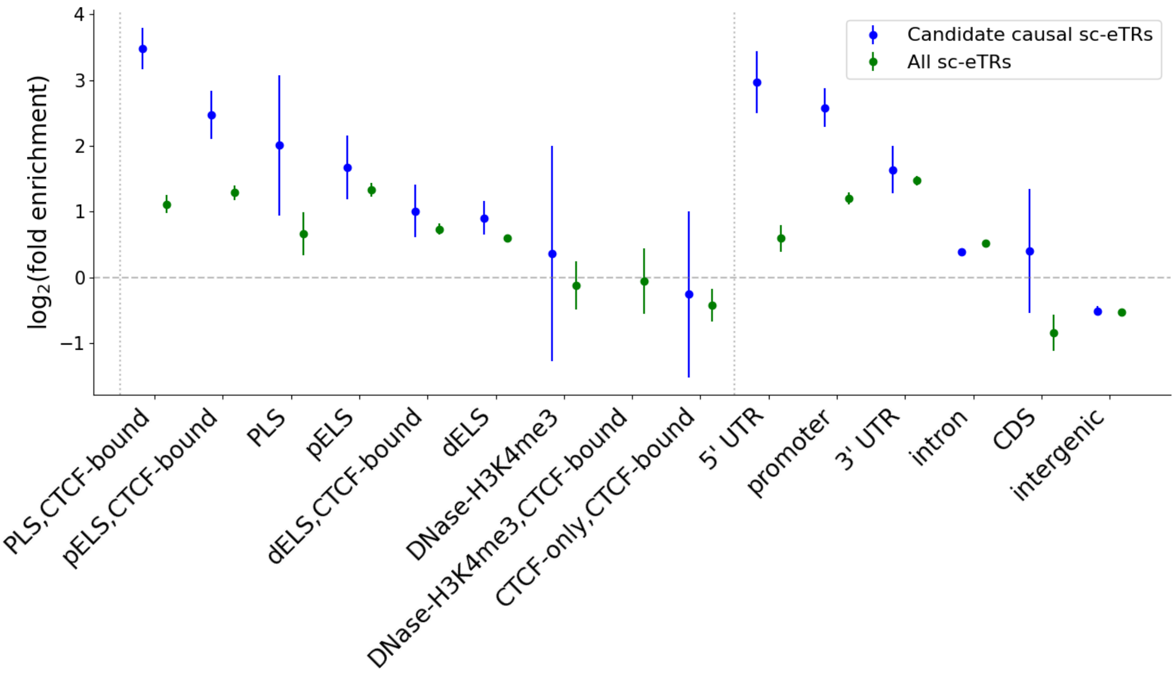


**Supplementary Figure 21: Enrichment of sc-eTRs and candidate causal sc-eTRs relative to all TRs genotyped.** The y axis denotes $\log_2$fold enrichment, comparing sc-eTRs (FDR <5%; n=112,100 sc-eTRs) and candidate causal sc-eTRs (n=1,490 TRs) to all TRs genotyped. Error bars correspond to 95% confidence intervals. ENCODE candidate *cis* regulatory element (cCRE) annotations are abbreviated as follows: promoter-like signature [PLS], proximal enhancer-like signature [pELS], distal enhancer-like signature [dELS], CTCF binding site failing to intersect another cCRE annotation [CTCF-only]. No candidate causal sc-eTRs intersected the 'DNase-H3K4me3,CTCF-bound' annotation.

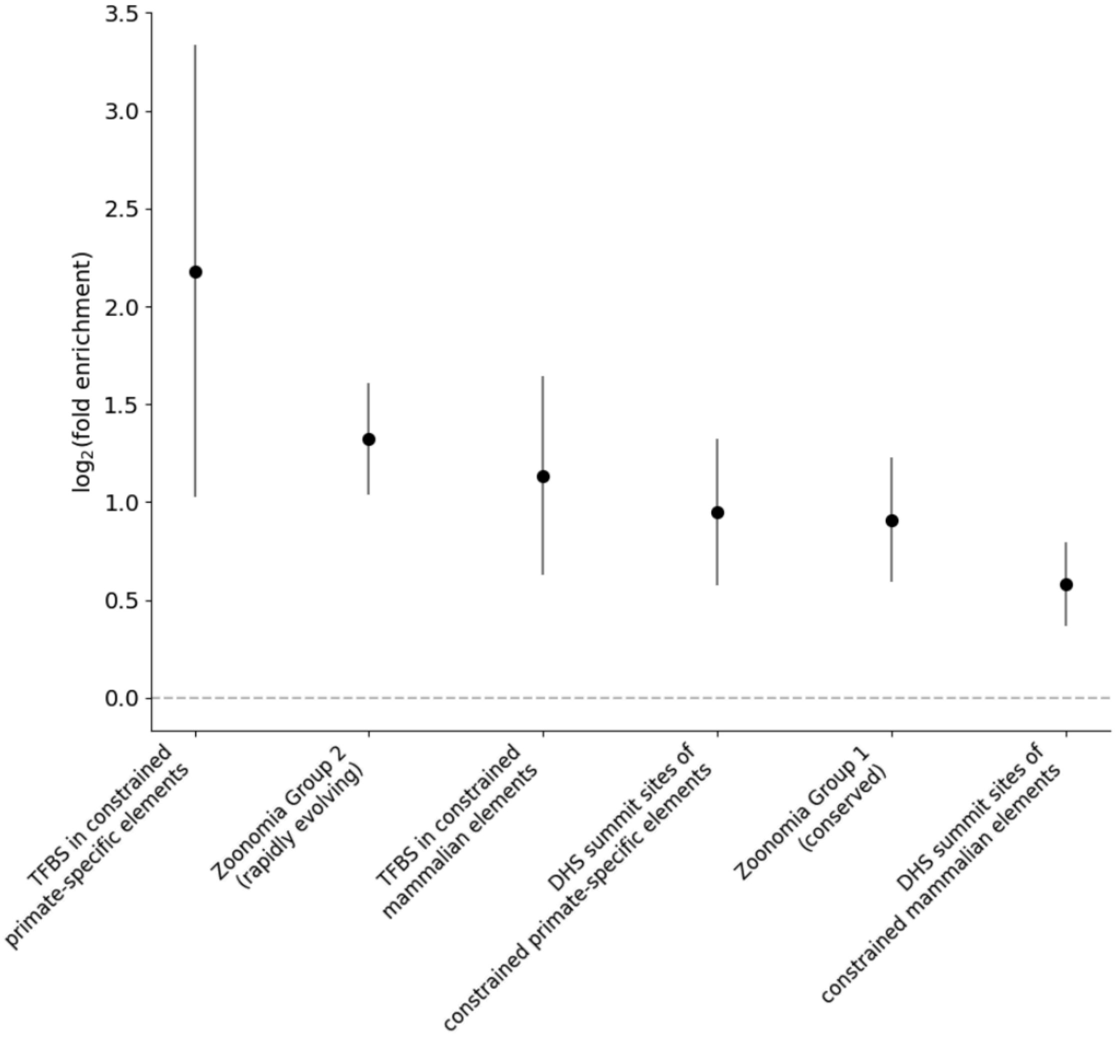


**Supplementary Figure 22: Enrichment of candidate causal sc-eTRs relative to all TRs genotyped in primate-specific[18,19] and Zoonomia[20] annotations.** The y axis denotes $\log_2$fold enrichment. Error bars correspond to 95% confidence intervals. Abbreviations are as follows: transcription factor binding site [TFBS]; DNase hypersensitivity site +/- 250 bp [DHS].

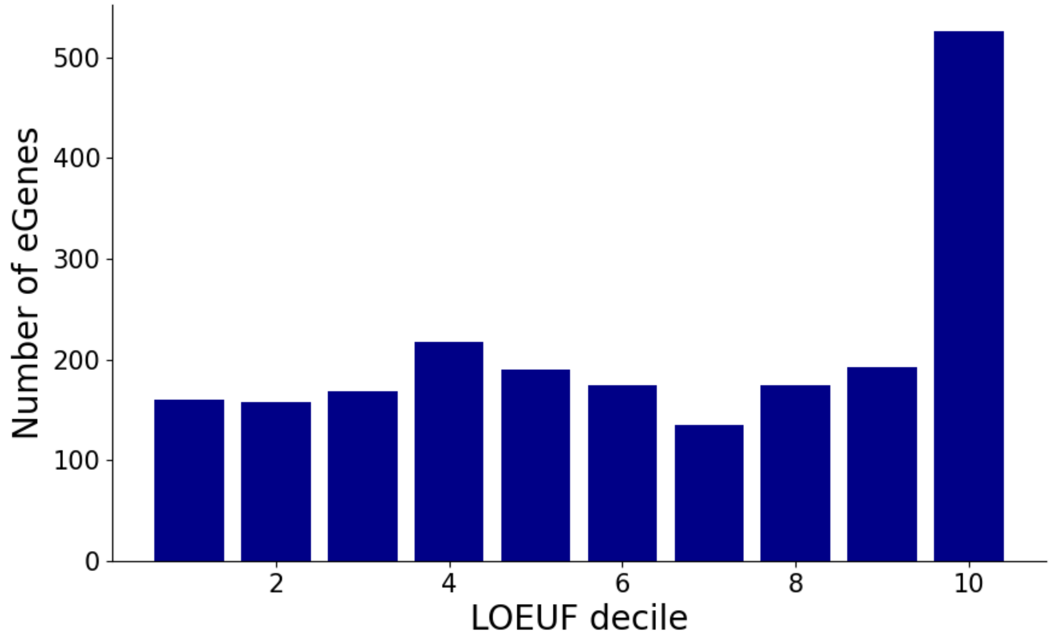


**Supplementary Figure 23**: **Number of eGenes (associated with fine-mapped sc-eTRs [PIP ≥ 0.7]) by loss-of-function observed/expected upper bound fraction (LOEUF) score decile.** Lower LOEUF deciles correspond to genes (based on MANE select transcripts) with greater constraint.

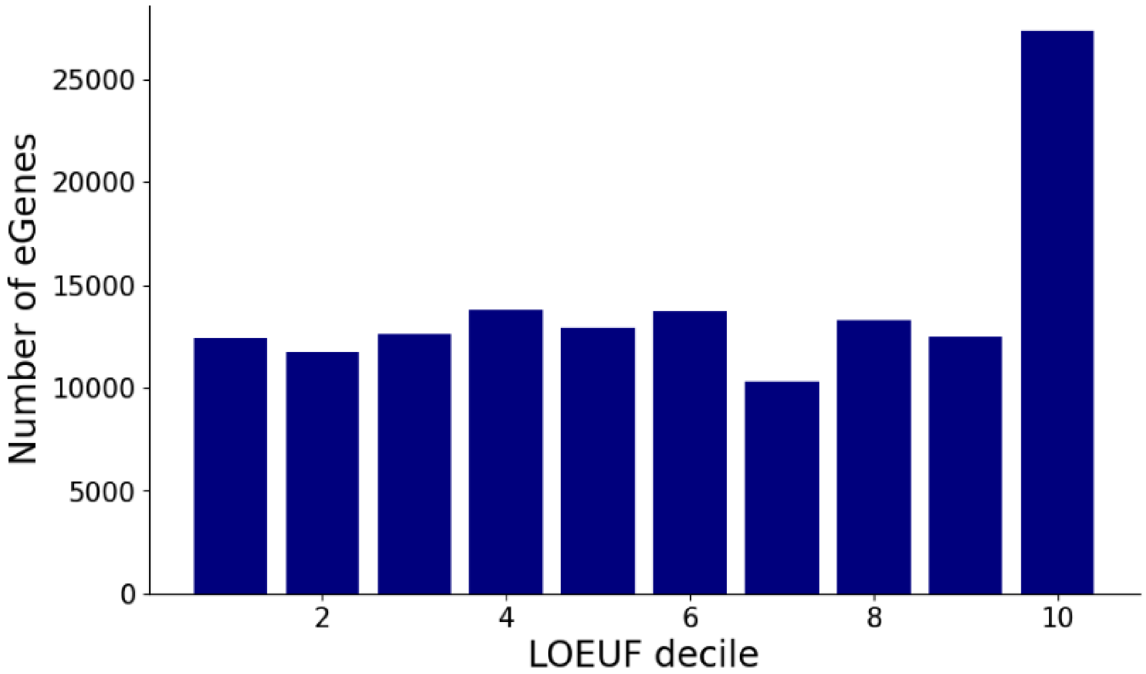


**Supplementary Figure 24**: **Number of eGenes (associated with sc-eSNVs [FDR <5%]) by loss-of-function observed/expected upper bound fraction (LOEUF) score decile.** Lower LOEUF deciles correspond to genes (based on MANE select transcripts) with greater constraint. The number of eGenes associated with sc-eSNVs was enriched in the least constrained genes, defined by the lowest loss-of-function observed/expected upper bound fraction (LOEUF) decile (binomial test $P = 4.1 \times 10^{-293}$).

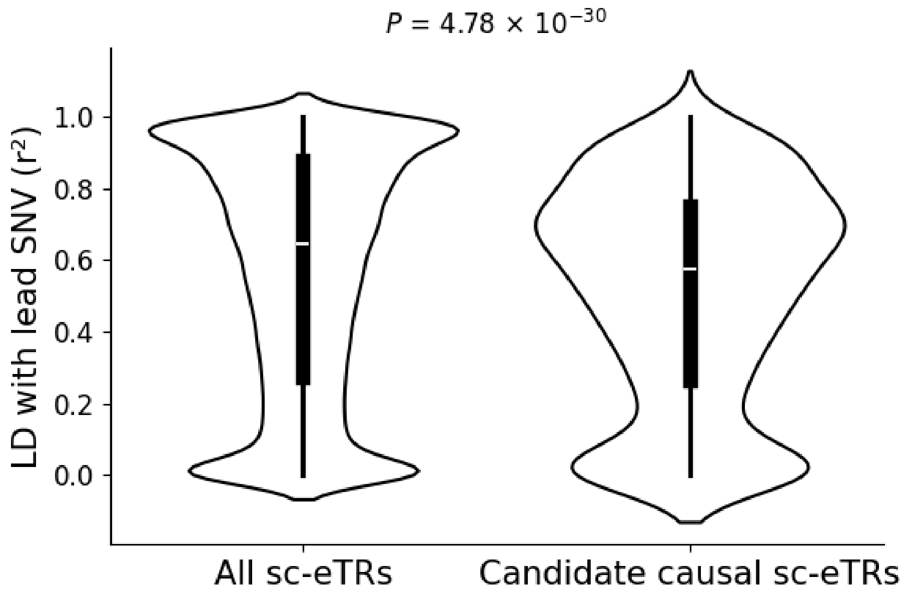


**Supplementary Figure 25: Distribution of linkage disequilibrium (LD) with the lead SNV of the respective *cis* window for all sc-eTRs and for candidate causal sc-eTRs.** Violin plots summarize the distribution of effect sizes with horizontal lines showing median values and boxes spanning from the 25th (Q1) to the 75th percentile (Q3). Whiskers extend to Q1 – 1.5 x IQR and Q3 + 1.5 x IQR, where IQR is the interquartile range. *Cis* window is the area within +/- 100 kb of the gene body. P value derived from a two-sided Mann Whitney U test.

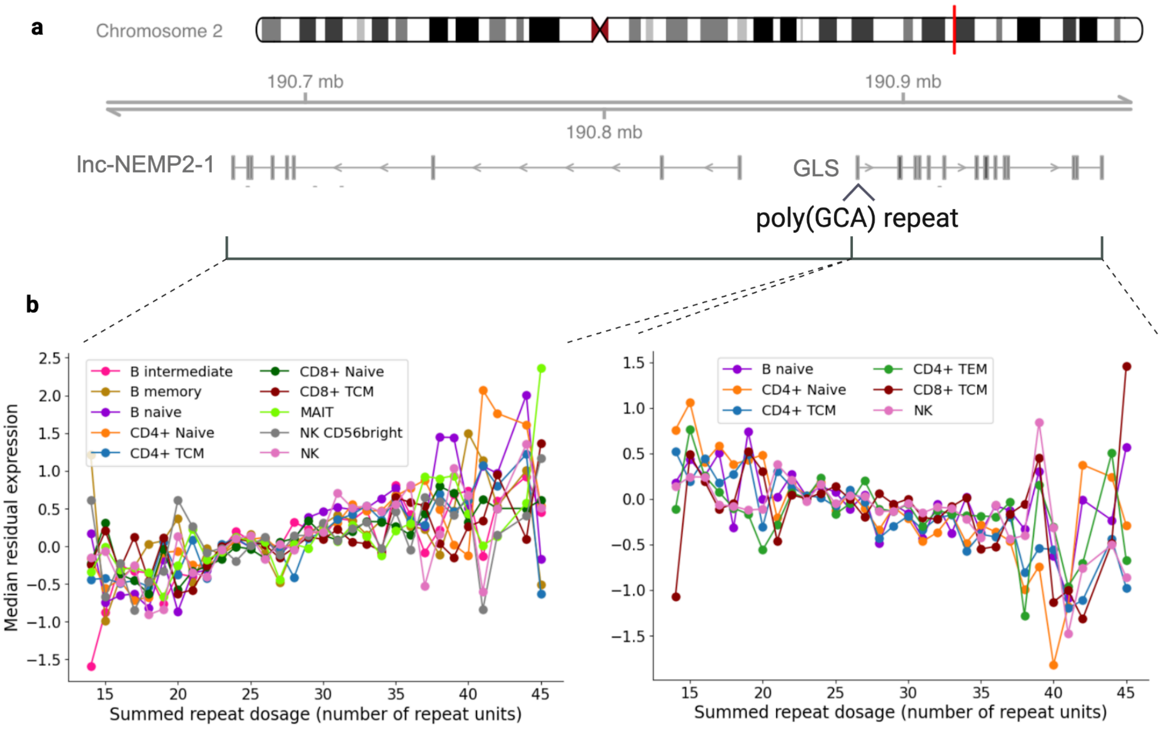


**Supplementary Figure 26: Cell type-specific associations of poly (GCA) repeat and expression of *GLS* and lnc-NEMP2-1.** **a**, Locus zoom plot of *GLS* and lnc-NEMP2-1 using Gviz[21], loaded with

GENCODE v44 track. **b**, Association between repeat length and median residual expression of lnc-NEMP2-1 (left) and *GLS* (right) in cell types where the sc-eTR was considered candidate causal for driving expression of the respective gene (n=1,925 individuals).

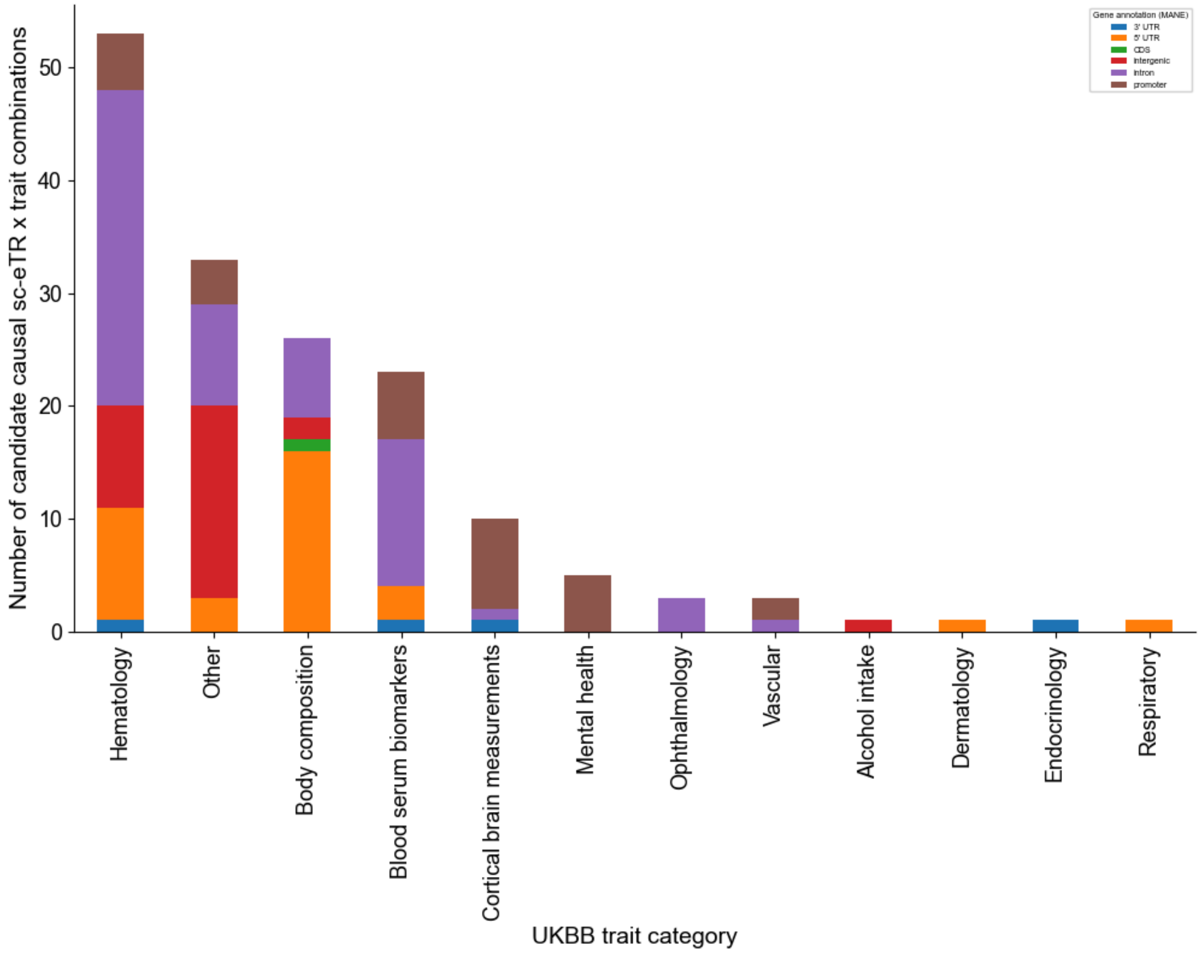


**Supplementary Figure 27: Barplot of the number of candidate causal sc-eTRs intersecting significant TR UKBB PheWAS hits, colored by gene annotation.**

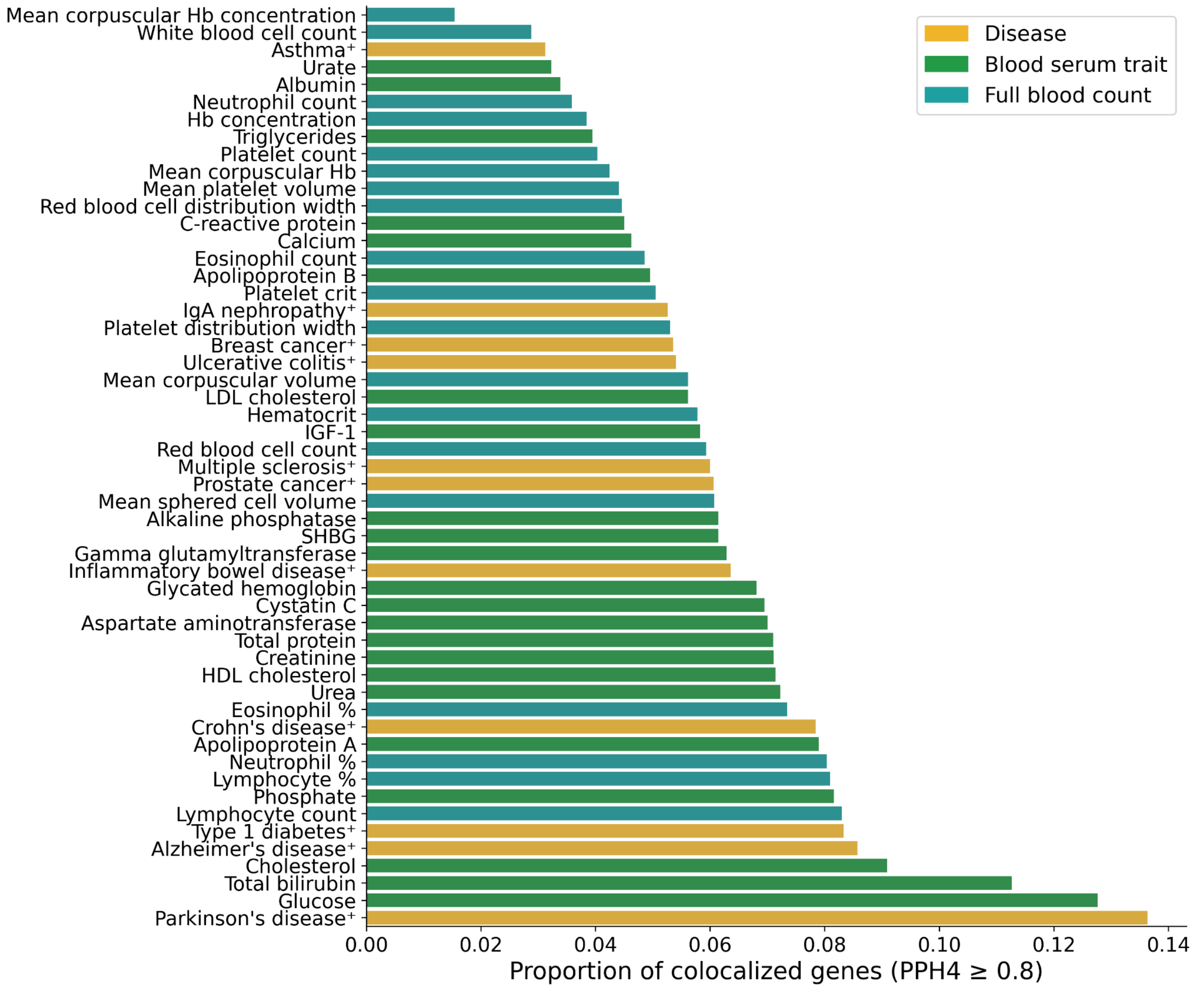


**Supplementary Figure 28: Barplot of the proportion of colocalized genes (PPH4 ≥0.8) containing a candidate causal sc-eTR, per GWAS trait tested.** Phenotypes marked with † were colocalized using only SNVs while unmarked phenotypes were colocalized using SNVs and imputed TRs.

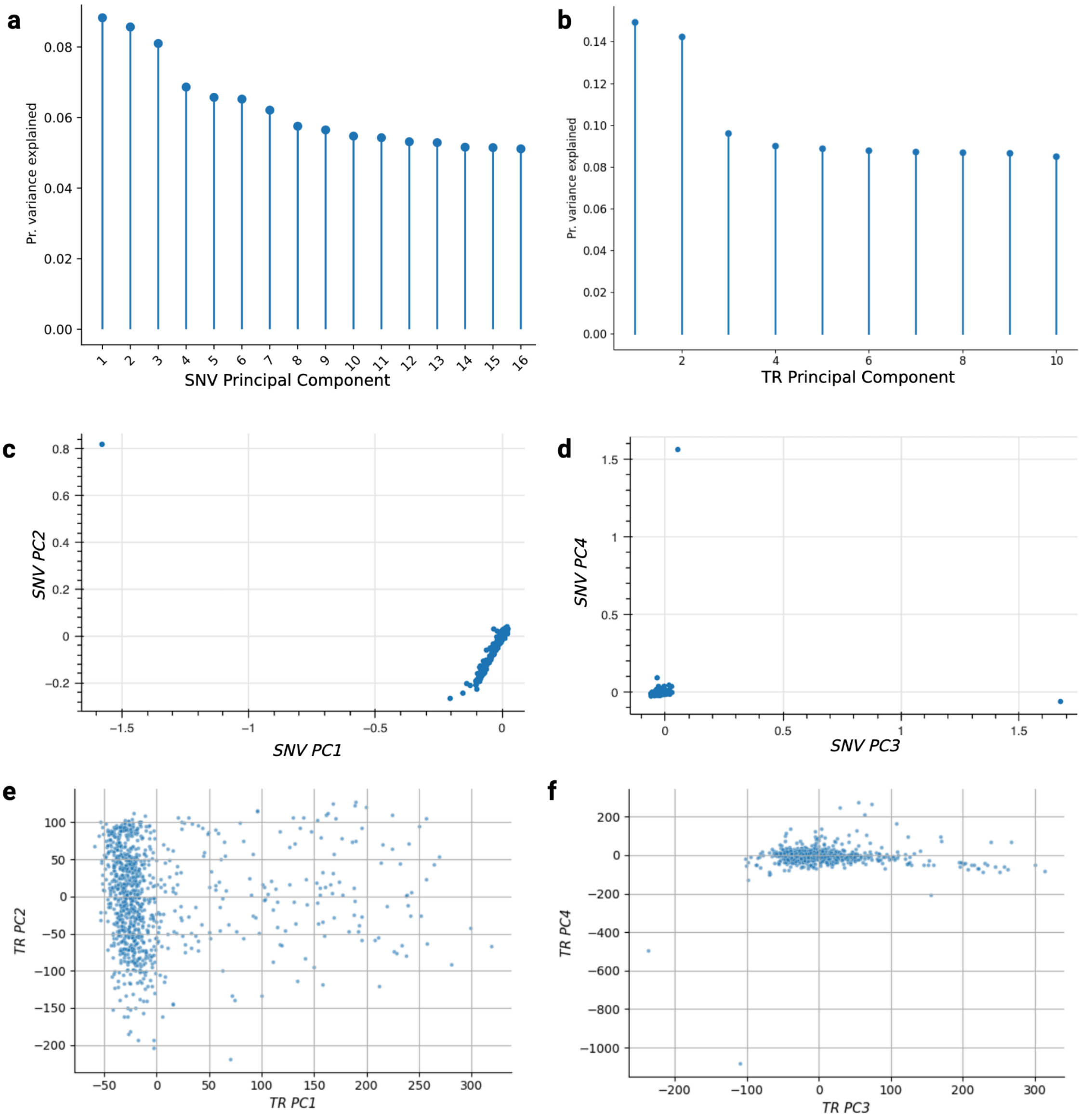


**Supplementary Figure 29: Principal components (PCs) for BioHEART cohort (n=975 individuals). a,** Scree plot showing the proportion of variance explained by the top 16 SNV-derived principal components for BioHEART cohort. The low variance explained and the absence of clustering across the top PCs are expected given the QC-filtered, European-ancestry cohort used for downstream eQTL analyses. **b**, Scree plot showing the proportion of variance explained by the top 10 TR-derived principal components for the BioHEART cohort. **c**, Scatterplot of SNV PC1 vs PC2. **c**, Scatterplot of SNV PC3 vs PC4. **d**, Scatterplot of TR PC1 vs PC2. High-loading TRs (top 5%) along PC2 axis exhibited significantly longer motifs (mean 4.07 bp vs. 2.23 bp; Mann–Whitney $P < 1\times10^{-300}$), higher GC content (0.33 vs. 0.15; $P < 1\times10^{-300}$), and longer lengths in the reference genome (22.55 bp vs. 17.11 bp; $P = 1.4\times10^{-167}$) than low-loading TRs (bottom 5%). **e**, Scatterplot of TR PC3 vs PC4.

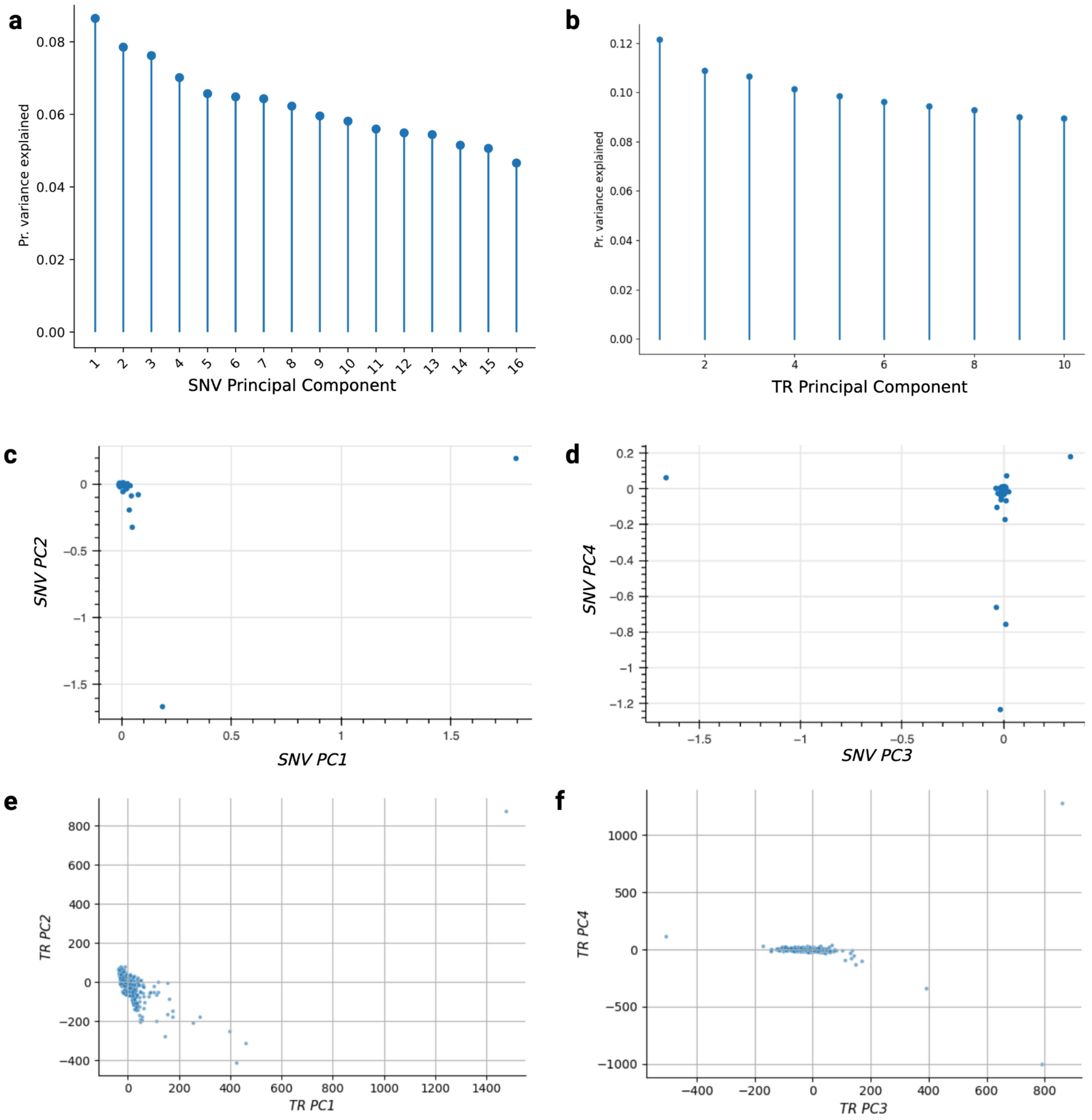


**Supplementary Figure 30: Principal components (PCs) for TOB cohort (n=950 individuals). a,** Scree plot showing the proportion of variance explained by the top 16 SNV-derived principal components for the TOB cohort. The low variance explained and the absence of clustering across the top PCs are expected given the QC-filtered, European-ancestry cohort used for downstream eQTL analyses. **b**, Scree plot showing the proportion of variance explained by the top 10 TR-derived principal components for the TOB cohort **c**, Scatterplot of SNV PC1 vs PC2. **c**, Scatterplot of SNV PC3 vs PC4. **d**, Scatterplot of TR PC1 vs PC2. TRs within the top 5% of PC1 contributors had slightly longer motifs (mean 3.08 bp vs. 2.25 bp; $P < 1\times10^{-300}$) and longer lengths in the reference genome (19.10 bp vs. 16.84 bp; $P = 1.8\times10^{-76}$) compared with those in the bottom 5%. **e**, Scatterplot of TR PC3 vs PC4.

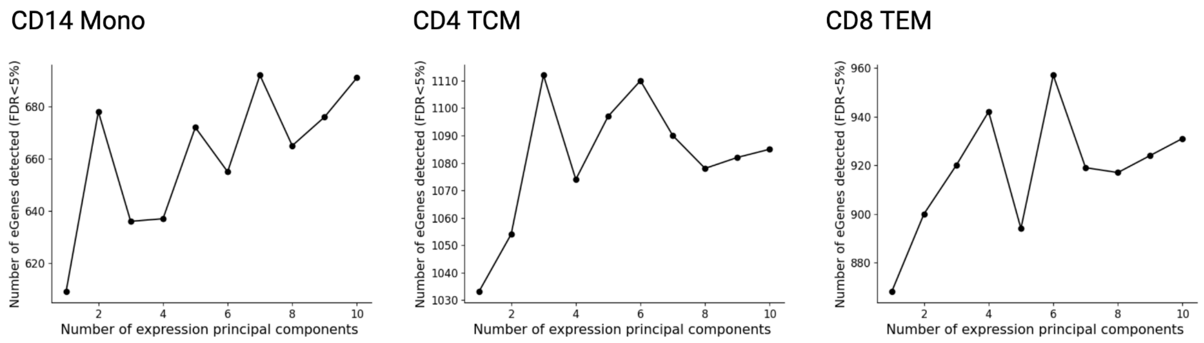


**Supplementary Figure 31: Changes in the number of *cis*-eGenes detected with the addition of expression principal components.** eGenes (gene-level FDR < 5%) in chromosome 1 were computed for CD14 monocytes (CD14 Mono), $CD4_{TCM}$, and $CD8_{TEM}$ cells after adjusting for a number of expression principal components ranging from 1 to 10. The x-axis shows the number of expression principal components accounted for by the model. The y-axis shows the number of eGenes detected at gene-level FDR < 5% using Storey q values[22] on ACAT[23]-corrected p-values. The final model used in the study adjusted for 6 expression principal components.

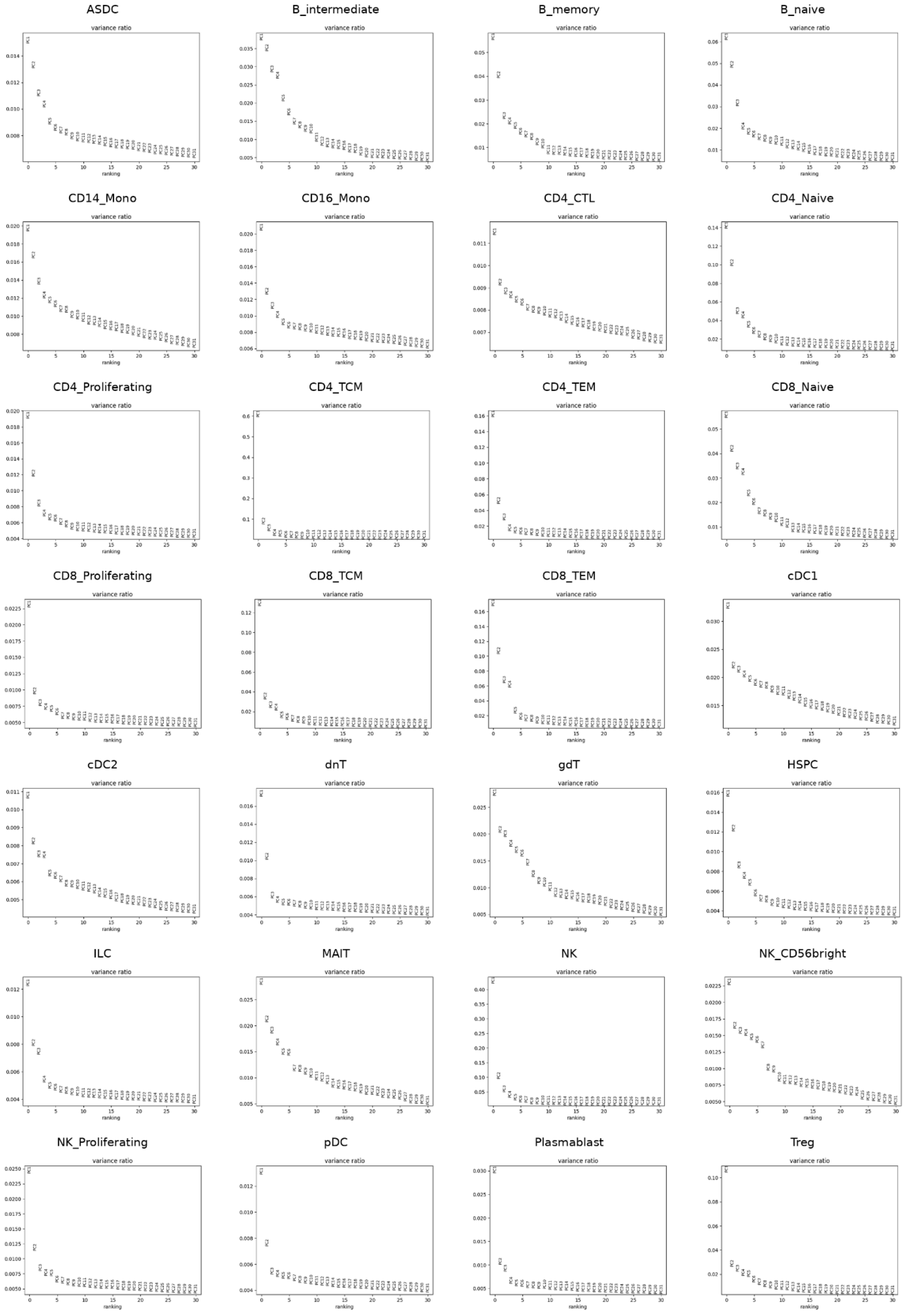
ASDC
B_intermediate
B_memory
B_naive
CD14_Mono
CD16_Mono
CD4_CTL
CD4_Naive
CD4_Proliferating
CD4_TCM
CD4_TEM
CD8_Naive
CD8_Proliferating
CD8_TCM
CD8_TEM
cDC1
cDC2
dnT
gdT
HSPC
ILC
MAIT
NK
NK_CD56bright
NK_Proliferating
pDC
Plasmablast
Treg
variance ratio
ranking

**Supplementary Figure 32: Variance ratio plots for the TOB cohort across 28 immune cell types.** Principal component analysis (PCA) was performed on the pseudobulk gene expression matrix of each immune cell type to quantify the proportion of variance explained by successive principal components. The plots depict the variance ratio for the top 31 principal components.

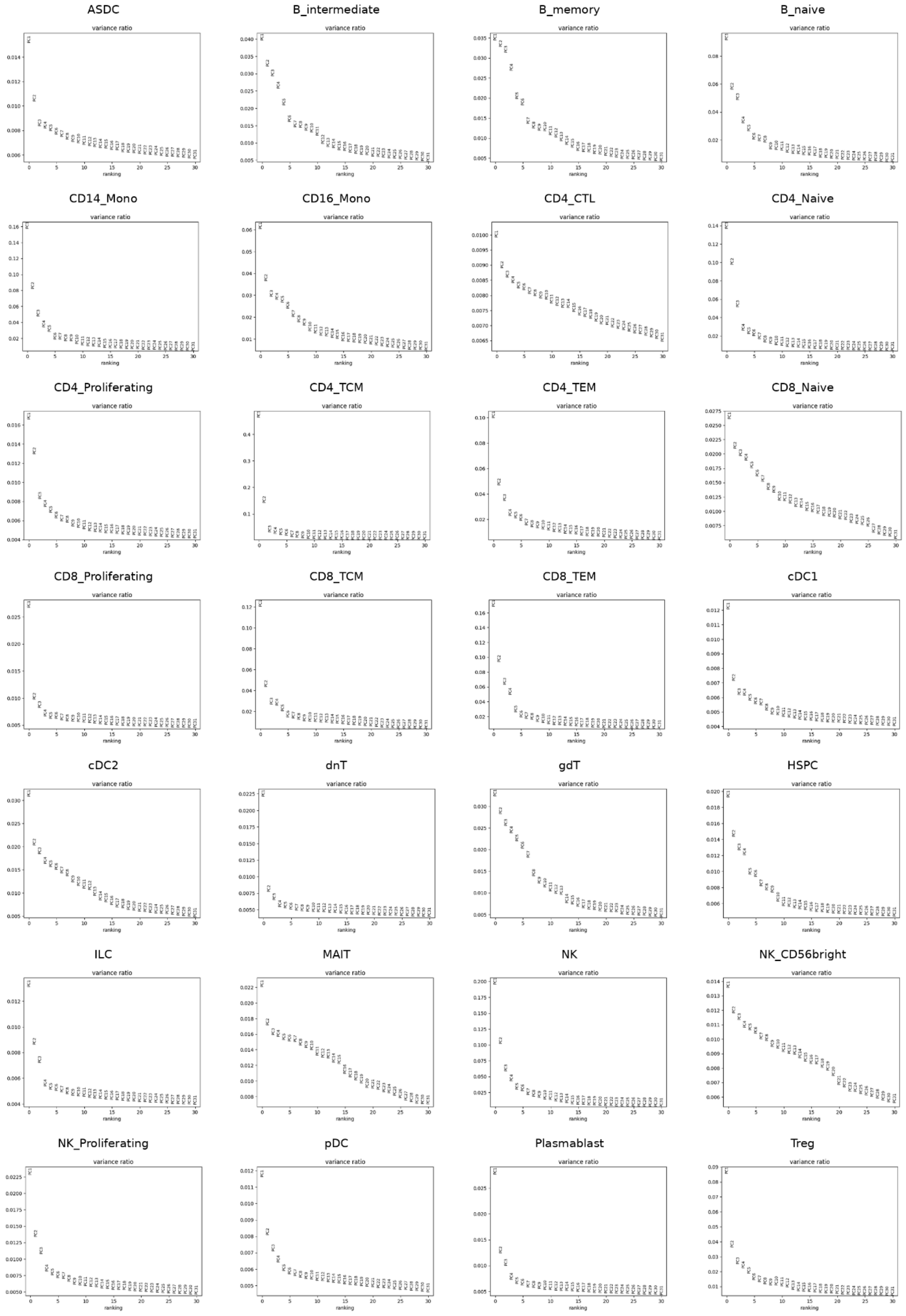


**Supplementary Figure 33: Variance ratio plots for the BioHEART cohort across 28 immune cell types.** Principal component analysis (PCA) was performed on the pseudobulk gene expression matrix of each immune cell type to quantify the proportion of variance explained by successive principal components. The plots depict the variance ratio for the top 31 principal components.

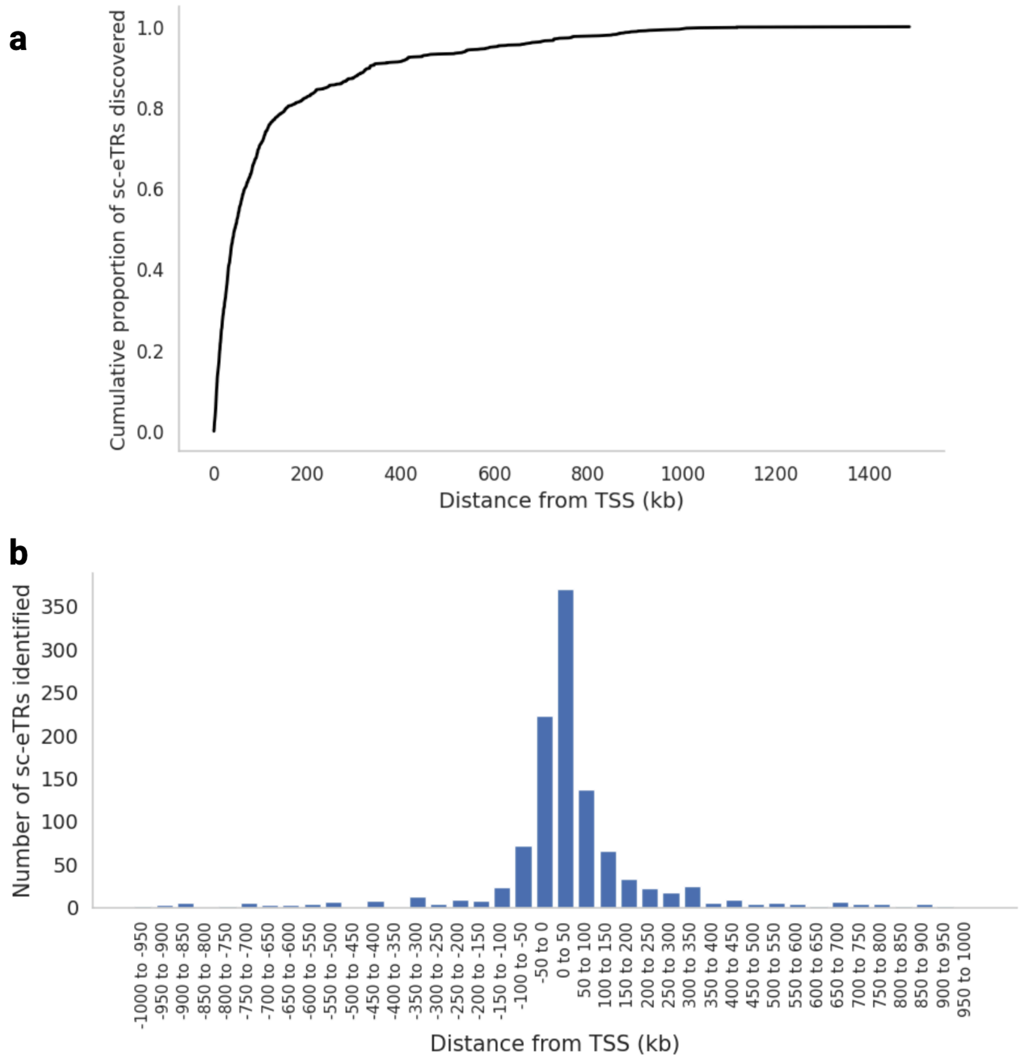


**Supplementary Figure 34: sc-eTR discovery in CD4$_{TCM}$ using +/- 1Mb *cis* window around the gene body. a**, Cumulative proportion of sc-eTRs discovered (FDR <5%) as a function of distance from the transcription start site (TSS). 71% of sc-eTRs were found within 100 kb of the TSS. **b**, Binned histogram plot of the number of sc-eTRs identified as a function of distance from the TSS.

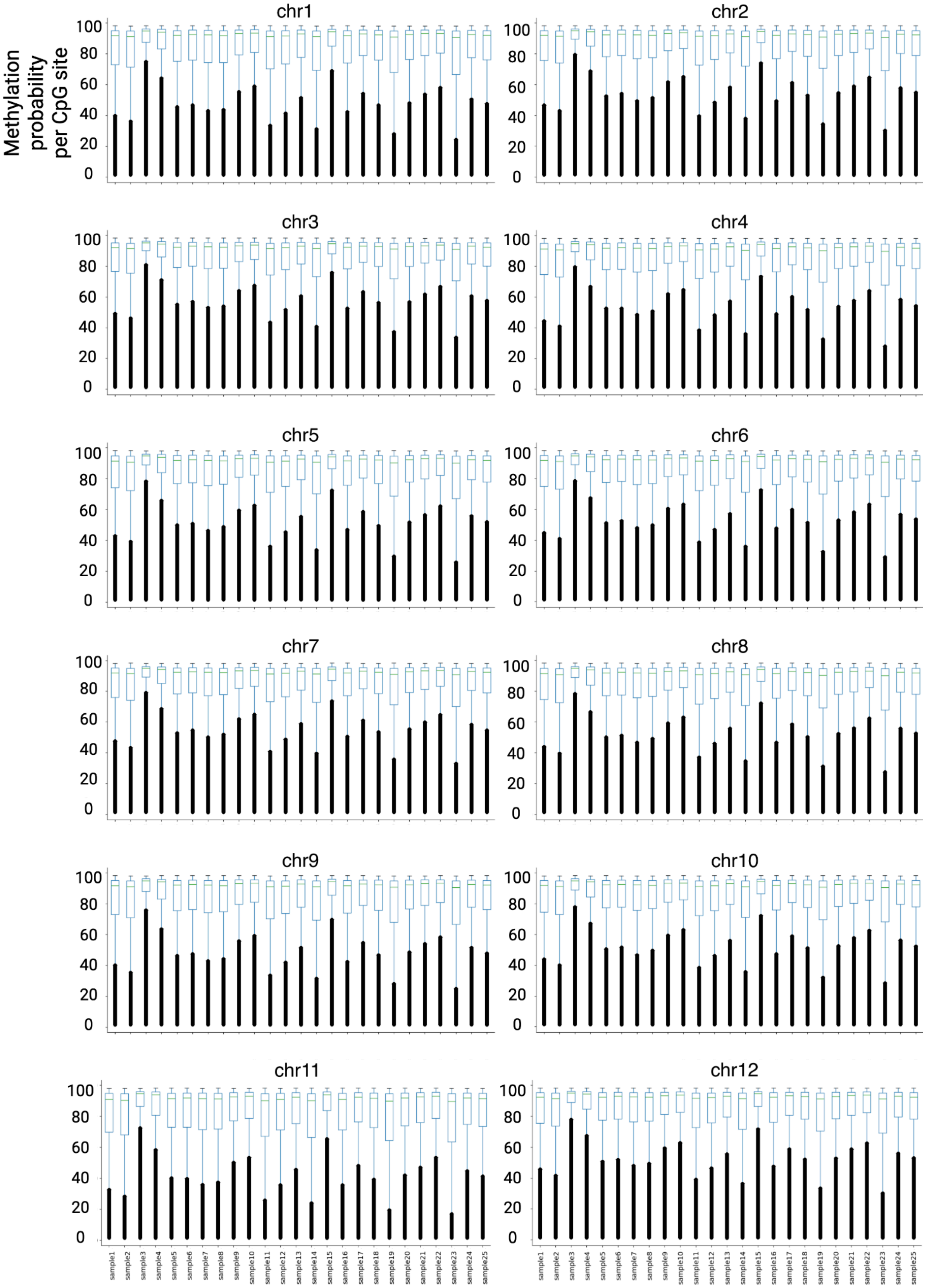
chr1
chr2
chr3
chr4
chr5
chr6
chr7
chr8
chr9
chr10
chr11
chr12
Methylation probability per CpG site
0
20
40
60
80
100
sample1
sample2
sample3
sample4
sample5
sample6
sample7
sample8
sample9
sample10
sample11
sample12
sample13
sample14
sample15
sample16
sample17
sample18
sample19
sample20
sample21
sample22
sample23
sample24
sample25

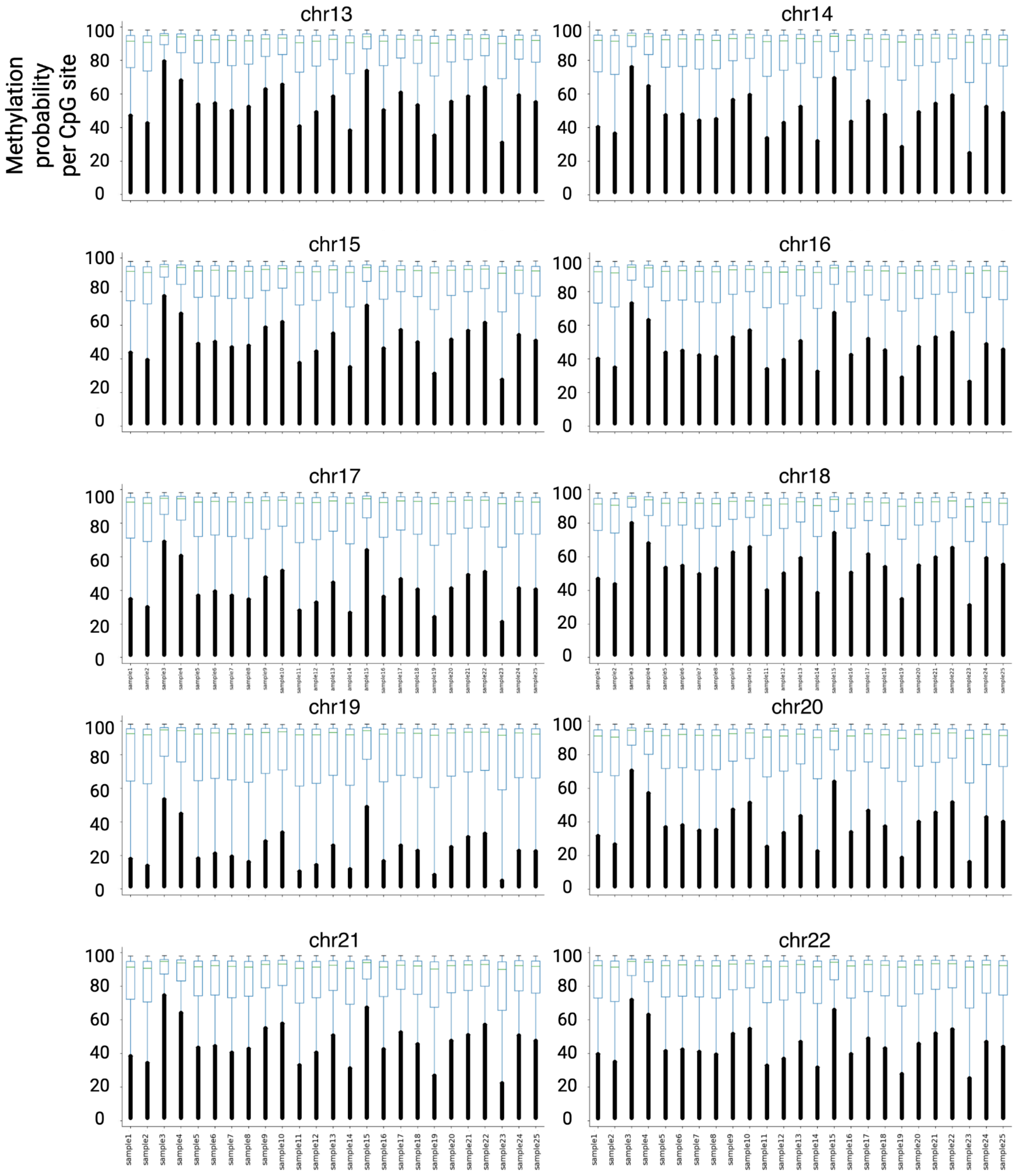


**Supplementary Figure 35: Distribution of methylation probability (per CpG site) across chromosomes and n=25 samples.** Methylation probability was derived from PacBio LR sequencing data using pb-CpG-tools (**Methods**).

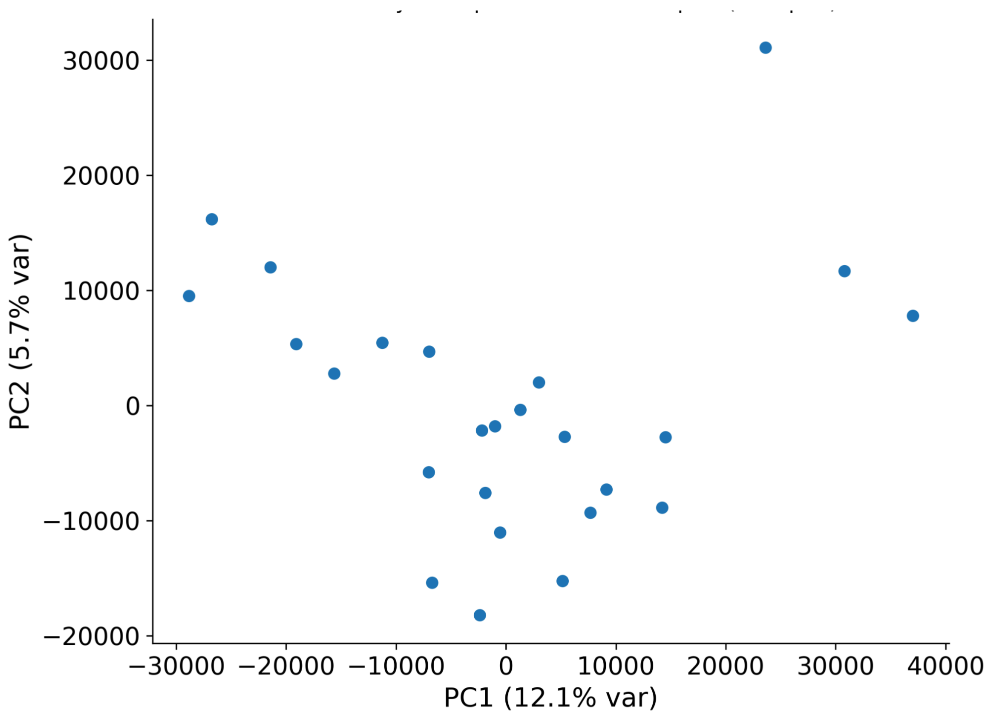


**Supplementary Figure 36: Principal component analysis of methylation probability values across 26,191,720 CpG sites in chr1-22.** Methylation probability was derived from PacBio LR sequencing data using pb-CpG-tools (**Methods**).

<u>Supplementary Tables</u>
*All supplementary tables, apart from Supplementary Table 8 below, are provided as supplementary files.*

**Supplementary Table 8: Candidate causal sc-eTRs intersecting known disease-associated loci.** Nominal p-value, effect size, and standard error (s.e.) estimates are reported for the cell type with the minimum nominal p-value (in bold). Cell types are listed in order of ascending nominal p-value.

| Coordinate (hg38) | Motif | eGene (associated disease) | Cell types | Nominal p-value | Effect size | s.e. |
|---|---|---|---|---|---|---|
| chr11:119206289-119206322 | CGG | *CBL* (Jacobsen Syndrome) | **NK**,$CD8_{TEM}$ | 2.09e-10 | -0.043825 | 0.006896 |
| chr2:190880872-190880920 | GCA | *GLS* (glutaminase deficiency) | **$CD4_{TCM}$**, CD4+ naive, $CD4_{TEM}$, NK, B naive, $CD8_{TCM}$ | 3.63e-56 | -0.040696 | 0.002577 |
| chr2:190880872-190880920 | GCA | lnc-NEMP2-1 [ENSG00000228509]<br><br>*TR locus is associated with glutaminase deficiency | **B naive**, NK, CD4+ naive, $CD4_{TCM}$, B memory, B intermediate, CD8+ naive, $CD8_{TCM}$, NK CD56bright, MAIT | 8.2715e-62 | 0.058543 | 0.003529 |
| chr12:50505001-50505022 | GGC | *DIP2B* (FRA12A type intellectual developmental disorder) | **NK** | 2.469e-81 | -0.051912 | 0.002718 |
| chr19:13207858-13207897 | CTG | *CACNA1A* (SCA6) | **NK** | 1.05e-8 | 0.044821 | 0.007833 |
| chr21:43776443-43776479 | CGC GGG GCG GGG | *CSTB* (EPM1) | **NK** | 1.895e-85 | 0.273381 | 0.013955 |
| chr5:146878727-146878757 | GCT | *PPP2R2B* (SCA12) | **Treg**,$CD4_{TCM}$ | 1.649e-26 | -0.068693 | 0.006447 |

| | | | | | | |
|---|---|---|---|---|---|---|
| chr9:27573528-27573546 | GGCCCC | *C9orf72* (ALS/FTD) | **CD4+ naive**,$CD4_{TCM}$ | 4.982e-62 | 0.064801 | 0.003899 |

**References**

1. Dolzhenko, E. *et al.* ExpansionHunter: a sequence-graph-based tool to analyze variation in short tandem repeat regions. *Bioinformatics* **35**, 4754–4756 (2019).
2. Mousavi, N., Shleizer-Burko, S., Yanicky, R. & Gymrek, M. Profiling the genome-wide landscape of tandem repeat expansions. *Nucleic Acids Res.* **47**, e90 (2019).
3. Willems, T. *et al.* Genome-wide profiling of heritable and de novo STR variations. *Nat. Methods* **14**, 590–592 (2017).
4. Mallick, S. *et al.* The Simons Genome Diversity Project: 300 genomes from 142 diverse populations. *Nature* **538**, 201–206 (2016).
5. Hinrichs, A. S. *et al.* The UCSC genome browser database: update 2006. *Nucleic Acids Res.* **34**, D590–8 (2006).
6. Benson, G. Tandem repeats finder: a program to analyze DNA sequences. *Nucleic Acids Res.* **27**, 573–580 (1999).
7. Pemberton, T. J., Sandefur, C. I., Jakobsson, M. & Rosenberg, N. A. Sequence determinants of human microsatellite variability. *BMC Genomics* **10**, 612 (2009).
8. Willems, T. *et al.* The landscape of human STR variation. *Genome Res.* **24**, 1894–1904 (2014).
9. Dolzhenko, E. *et al.* Detection of long repeat expansions from PCR-free whole-genome sequence data. *Genome Res.* **27**, 1895–1903 (2017).
10. Tanudisastro, H. A., Deveson, I. W., Dashnow, H. & MacArthur, D. G. Sequencing and characterizing short tandem repeats in the human genome. *Nat. Rev. Genet.* **25**, 460–475 (2024).
11. van Kuilenburg, A. B. P. *et al.* Glutaminase deficiency caused by short tandem repeat expansion in *GLS*. *N. Engl. J. Med.* **380**, 1433–1441 (2019).
12. Calder, P. C. & Yaqoob, P. Glutamine and the immune system. *Amino Acids* **17**, 227–241 (1999).
13. Bakhtiari, M. *et al.* Variable number tandem repeats mediate the expression of proximal

genes. *Nat. Commun.* **12**, 2075 (2021).

14. Fotsing, S. F. *et al.* The impact of short tandem repeat variation on gene expression. *Nat. Genet.* **51**, 1652–1659 (2019).
15. Gymrek, M. *et al.* Abundant contribution of short tandem repeats to gene expression variation in humans. *Nat. Genet.* **48**, 22–29 (2016).
16. Urbut, S. M., Wang, G., Carbonetto, P. & Stephens, M. Flexible statistical methods for estimating and testing effects in genomic studies with multiple conditions. *Nat. Genet.* **51**, 187–195 (2019).
17. Alquicira-Hernandez, J., Sathe, A., Ji, H. P., Nguyen, Q. & Powell, J. E. scPred: accurate supervised method for cell-type classification from single-cell RNA-seq data. *Genome Biol.* **20**, 264 (2019).
18. Kuderna, L. F. K. *et al.* A global catalog of whole-genome diversity from 233 primate species. *Science* **380**, 906–913 (2023).
19. Kuderna, L. F. K. *et al.* Identification of constrained sequence elements across 239 primate genomes. *Nature* **625**, 735–742 (2024).
20. Andrews, G. *et al.* Mammalian evolution of human cis-regulatory elements and transcription factor binding sites. *Science* **380**, eabn7930 (2023).
21. Hahne, F. & Ivanek, R. Visualizing genomic data using Gviz and Bioconductor. *Methods Mol. Biol.* **1418**, 335–351 (2016).
22. Storey, J. D. The positive false discovery rate: a Bayesian interpretation and the q-value. *Ann. Stat.* **31**, 2013–2035 (2003).
23. Liu, Y. *et al.* ACAT: A fast and powerful p value combination method for rare-variant analysis in sequencing studies. *Am. J. Hum. Genet.* **104**, 410–421 (2019).
